# Unsupervised visual learning is revealed for task-irrelevant natural scenes due to reduced attentional suppression effects in visual areas

**DOI:** 10.1101/2024.07.31.605957

**Authors:** Takeo Watanabe, Yuka Sasaki, Takuro Zama, Julian R Matthews, Daiki Ogawa, Kazuhisa Shibata

**Author notes:** Equally contributed (Takeo Watanabe, Yuka Sasaki, Takuro Zama).

## Abstract

Unsupervised learning—learning through repeated exposure without instruction or reward—is central to both machine learning and human cognition, including language acquisition and statistical learning. However, its role in visual perceptual learning (VPL) remains debated, as previous studies have not shown VPL for task-irrelevant but visible features, particularly in artificial stimuli. Here, we show that task-irrelevant exposure to natural scene images induces robust VPL, while artificial images that lack complex structure characteristics of natural scene images, known as higher-order statistics, do not. Behavioral and fMRI results suggest that although unsupervised learning underlies VPL, it can be suppressed by top-down attention. Higher-order statistics may evade this suppression, possibly because their slower processing reaches visual areas beyond V1 outside the optimal temporal window for attentional suppression. These findings suggest that unsupervised learning underlies VPL, but its occurrence depends on both higher-order stimulus structure and the brain’s attentional gating mechanisms.

## Introduction

Since birth, humans are immersed in a complex environment filled with continuous visual stimulation. Among this stream of information, frequently encountered features—such as object contours or motion patterns—are often ecologically significant. It is widely believed that the visual system becomes increasingly sensitive to such features through unsupervised learning—a process occurring without instruction, reinforcement, or feedback. This principle underpins many machine learning models and is supported across domains of cognitive science, including language acquisition^1–4^ and statistical learning^5–10^. These lines of evidence suggest that unsupervised learning plays a central role in human learning.

But does unsupervised learning also underlie visual perceptual learning (VPL)—the long-term improvement in visual performance on basic features like orientation or motion direction as a result of visual experience? Some studies suggest that feedback or top-down processing is essential for VPL (e.g., ^11,12^), raising the possibility that VPL relies on mechanisms distinct from those of general forms of human learning in which unsupervised learning occurs. However, a series of studies have demonstrated that VPL can occur through repeated presentation of sub-threshold *task-irrelevant* features during unrelated task performance^13–15^. A feature that is to be learned under such a context is referred to as a task-irrelevant feature.

Importantly, this form of VPL of a task-irrelevant feature appears to be context-dependent: learning occurs when task-irrelevant features are near or below perceptual threshold and temporally linked to task-relevant events^13,14,16–18^, or when they are spatially distant from task-relevant stimuli (e.g., ^19,20^). This contextual dependency raises a critical question: is unsupervised learning a fundamental mechanism of VPL, or does it operate only under specific conditions?

To address this question, we compared VPL of a task-irrelevant feature using natural scene versus artificial images, asking whether unsupervised learning supports both. Notably, the vast majority of VPL studies have employed artificial stimuli composed of simple structures in which different spatial frequencies and orientations are unrelated^15,21–24^. In contrast, natural scenes consist of more global and complex spatial structures in addition to the simpler structures present in artificial images. Moreover, studies demonstrating VPL of task-irrelevant features have primarily used sub-threshold features^13,14,18,25^. The significant differences in stimulus contexts—specifically, the structural complexity of artificial versus natural scene images—make it uncertain whether VPL of task-irrelevant natural scenes would occur in the same (e.g., sub-threshold) manner as VPL of task-irrelevant artificial images.

We found that repeated exposure to supra-threshold natural scene images as task-irrelevant stimuli during the performance of a main task unrelated to the image led to VPL, whereas supra-threshold artificial images presented in the same conditions did not. In contrast, passive viewing of both natural scene and artificial images without task engagement led to VPL regardless of the stimulus type (natural scene vs artificial images).

Do these differential results imply that unsupervised learning occurs only for natural scene images—in a specific stimulus context? To explore this, we examined the underlying processing that caused these findings. Results of a series of experiments suggest that natural scene images contain higher-order statistical structures from which the learned feature can be constructed and are less susceptible to attentional suppression. In contrast, no learning occurred for supra-threshold task-irrelevant artificial images or other images lacking these higher-order structures.

Further behavioral and fMRI experiments suggest how the brain played a role in differential results between VPL of task-irrelevant natural scene images and artificial images: while attentional source areas may send comparable suppression signals for both image types, the slower processing of higher-order statistics in natural scene images—particularly in visual areas beyond V1—may miss the temporal window for effective top-down suppression. In contrast, the faster processing of artificial images allows suppression to occur in time, preventing learning. This temporal mismatch may explain why VPL emerges for natural scene images but not artificial images under identical conditions.

These results suggest that unsupervised learning is fundamental to VPL, but its expression is modulated by attentional mechanisms in a context-dependent manner—attentional mechanisms can suppress learning particularly for artificial stimuli that lack higher-order statistical structure from which a feature is reconstructed.

## Results

### VPL occurs for supra-threshold task-irrelevant natural scene images, but not for artificial images

In Experiment 1 (see Stimuli and Experiment 1 in Methods for details), we examined whether exposure to natural scene (NS) images leads to VPL of the dominant orientation in a different manner than exposure to artificial images. Two groups of 12 participants were exposed to supra-threshold, task-irrelevant images: one group to NS images (NS group), and the other to artificial images with the same orientation and spatial frequency spectra, termed Fourier-scrambled (FS) images (FS group).

It is widely accepted that NS images are characterized by *higher-order statistics*, *marginal statistics*, *and lower-order statistics*. Higher-order statistics convey complex spatial relationships such as edges, textures, and contours typical of natural scenes (e.g., ^26,27^). Marginal statistics include kurtosis, which reflects the "tailedness" or "peakedness", and skewness, which indicates asymmetry of the luminance histogram of natural scenes (e.g., ^28,29^). Lower-order statistics refer to properties such as orientation and spatial frequency that are locally and independently processed in the visual system (e.g., ^30,31^).

FS images, by contrast, lack higher-order statistics such as global structures and marginal statistics such as kurtosis and skewness. Images containing the higher-order and marginal statistics are processed in visual areas beyond V1^32–34^, whereas lower-order statistics are primarily processed in V1^35–39^. Thus, theoretically these two forms of images should be processed in similar ways in V1 but differently in visual areas beyond V1. Most prior VPL studies have used artificial images with simple characteristics like those of FS images, which can be fully processed within V1.

The complete procedure with both NS and FS groups consisted of one day of the pre-test stage, 10 days of the exposure+task stage, and one day of the post-test stage (Fig. 1a).

**Fig. 1:**
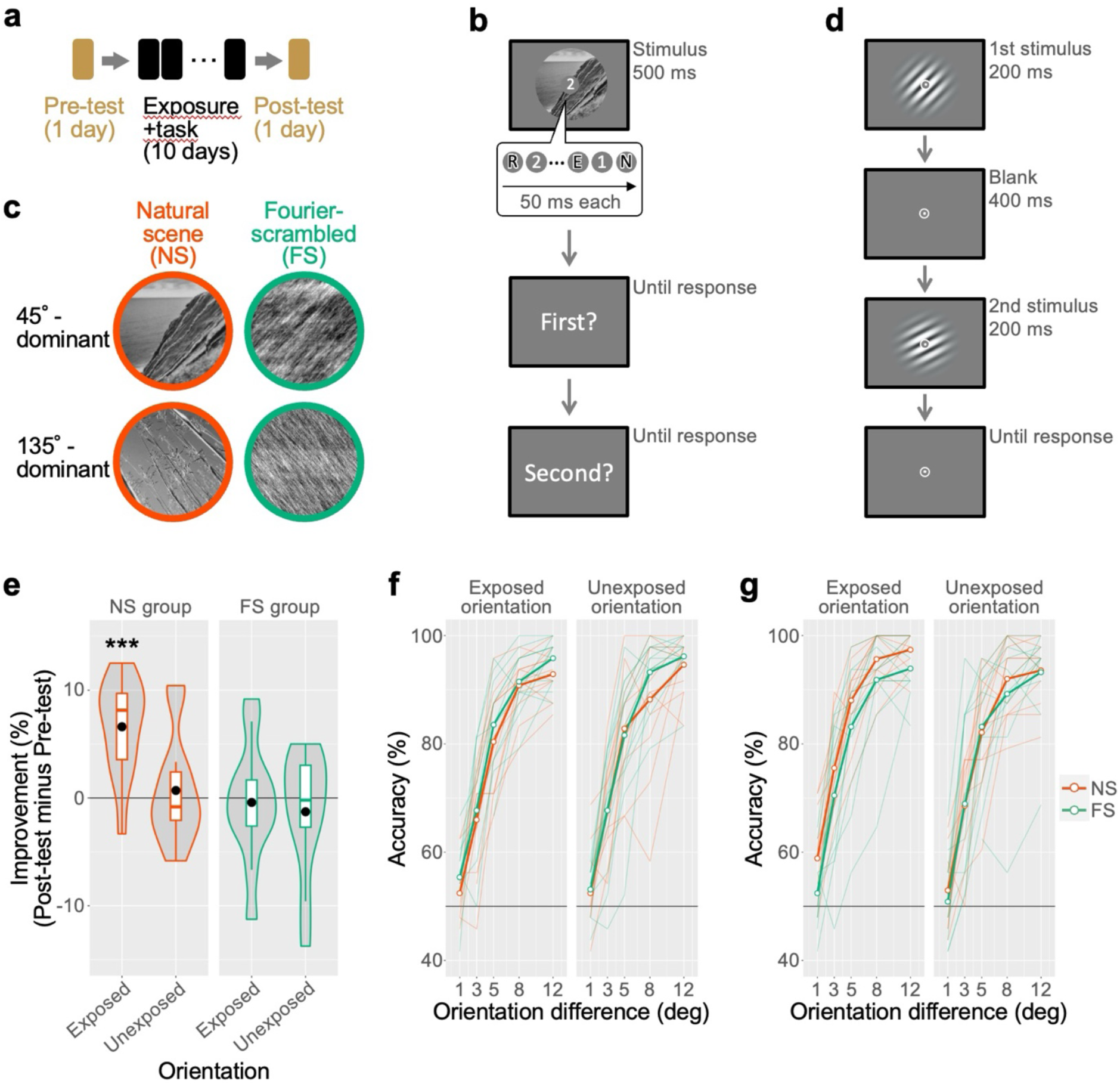
Methods and results of Experiment 1 **a**, The complete experiment consisted of a 10-day exposure+task stage, preceded and followed by 1-day pre- and post-test stages. **b**, Procedure of a single trial in the rapid serial visual presentation (RSVP) task during the exposure+task stage. While a sequence of white digits and black letters was presented, participants were asked to make two successive responses to report the two white digits in the order they appeared. **c**, Example stimuli of natural scene images (NS, highlighted with orange rings) and Fourier-scrambled images (FS, highlighted with green rings) presented as task-irrelevant during the exposure+task stage. **d**, Procedure of the orientation discrimination task in the pre-test and post-test stages. Participants reported whether the orientations of the first and second stimuli were the same or different. **e**, Improvement in performance on the orientation discrimination task (post-test minus pre-test) for the NS (left panel) and FS (right panel) groups. For the NS group, a two-way repeated-measures ANOVA revealed a significant interaction between Test (pre vs post) and Orientation (exposed vs unexposed) (see Table S1 for detailed results). Subsequent analyses showed a significant simple effect of Test at the exposed orientation (F_1,11_ = 26.5228, P = 0.0003, partial η^2^ = 0.7068, 95% CI of partial η^2^ = [0.2738–0.8931], BF_10_ = 106.29), but not at the unexposed orientation (F_1,11_ = 0.2262, P = 0.6436, partial η^2^ = 0.0202, 95% CI of partial η^2^ = [0.0000–0.1837], BF_10_ = 0.32). For the FS group, a two-way repeated-measures ANOVA showed no significant effects. Box plots are overlaid on violin plots. Black dots indicate the mean accuracies across participants, and violin plots show kernel probability densities of individual accuracies. ***P < 0.005. **f**, Performance on the orientation discrimination task in the pre-test stage. **g**, Performance in the post-test stage. Thick lines indicate mean accuracy across participants, and thin lines represent individual accuracies.

In the exposure+task stage, participants were asked to perform a rapid serial visual presentation (RSVP) task, known to suppress visible and task-irrelevant features^13,14,17,18,40,41^. In each trial, a sequence of eight alphabetical letters and two digits were presented in a random order for each participant at the center of the display (Fig. 1b). After the presentation of the sequence, the participants were instructed to make two successive responses to report the two digits, in the order they appeared (see Fig. S1a for performance in the RSVP task). While the RSVP task was performed, an NS (Fig. 1c, highlighted with orange rings) or FS (Fig. 1c, highlighted with green rings) image, as task-irrelevant stimuli, was being presented in the background to the NS or FS group, respectively. We created two sets of 40 NS images, each set exhibiting the dominant orientation power peaking at 45° or 135°, referred to as 45°-dominant and 135°-dominant NS images, respectively (Fig. S2). Perceiving the dominant orientation in NS is ecologically important for identifying or categorizing the NS^42,43^. Importantly, the FS images consisted of the identical spectra of orientations and spatial frequencies and the mean luminance and variance of luminance to those of the NS images used for the NS group (see Stimuli in Methods for details of creating the FS images). In each of the NS and FS groups, half of participants were exposed to the 45°-dominant images as task-irrelevant and the other half to the 135°-dominant images. The dominant orientation in the NS and FS images had been proven to be supra-threshold and almost equally visible (see Control Experiment in Methods and Fig. S3).

The pre-test and post-test stages examined how exposure to the task-irrelevant NS and FS images influenced the discriminability of the dominant orientations in the presented task-irrelevant images. We used Gabor patches in both the pre- and post-test stages for both the NS and FS groups to examine whether the most dominant orientation in NS and FS images becomes sensitized through exposure to these images. All participants were asked to perform a discrimination task (Fig. 1d) for the dominant orientation to which they were exposed during the exposure+task stage (exposed orientation) and the orientation orthogonal to the exposed orientation (unexposed orientation). In each trial, participants were asked to report whether the orientations of the two Gabor patches presented successively were the same or different. In half the trials, one Gabor patch was at the reference orientation, and the other patch was at an orientation differing by ±Δ degrees from the reference. In the other half, both Gabor patches were at the reference orientation. The reference orientation for each block was either exposed or unexposed orientation and alternated between blocks. The Δ value for each block was 12, 8, 5, 3, or 1 degree, decreasing every two blocks consisting of 48 trials. Performance in each of the pre- and post-stages was defined as the mean accuracy averaged across the Δ values of 12, 8, 5, 3, and 1 degree. An improvement in performance was defined by the subtraction of the performance in the pre-test stage from that in the post-test stage. We employed this combination of task and procedure, rather than the classical pairing of a two-interval forced-choice task and a staircase procedure, because a well-known previous study on task-irrelevant VPL of orientation used the same task without a staircase procedure^19^, allowing a more straightforward comparison between our results and those of the previous study.

Fig. 1e shows improvements in the NS and FS groups (see Fig. 1f and g for task accuracies before the averaging and subtraction). To test VPL of the global orientation in the NS and FS images, we applied a three-way mixed-model ANOVA on performance with Group (NS vs FS) as the between-participant factor and Test (pre vs post) and Orientation (exposed vs unexposed) as the within-participant factors (see Table S1 for detailed results of ANOVAs). Since there was a significant three-way interaction (F_1,22_ = 5.8216, P = 0.0246, partial η^2^ = 0.2092, 95% CI of partial η^2^ = [0.0036–0.4781], BF_10_ = 2.52), we proceeded to a two-way ANOVA with the factors of Test and Orientation to explore the source of the interaction in each group (See Statistics in Methods for details). In the NS group, the results of the ANOVA showed significant effects of Test (F_1,11_ = 9.2612, P = 0.0112, partial η^2^ = 0.4571, 95% CI of partial η^2^ = [0.0412–0.7403], BF_10_ = 291.85), Orientation (F_1,11_ = 14.1095, P = 0.0032, partial η^2^ = 0.5619, 95% CI of partial η^2^ = [0.0835–0.8458], BF_10_ = 25.42) and significant interaction between them (F_1,11_ = 19.3135, P = 0.0011, partial η^2^ = 0.6371, 95% CI of partial η^2^ = [0.3418–0.8191], BF_10_ = 49.76).

Subsequently, we found a significant simple effect of Test for the exposed orientation (F_1,11_ = 26.5228, P = 0.0003, partial η^2^ = 0.7068, 95% CI of partial η^2^ = [0.2738–0.8931], BF_10_ = 106.2929) in the NS group. On the other hand, in the FS group the results of the ANOVA showed no significant effect of Test (F_1,11_ = 0.3659, P = 0.5575, partial η^2^ = 0.0322, 95% CI of partial η^2^ = [0.0000–0.2383], BF_10_ = 0.20), Orientation (F_1,11_ = 0.8083, P = 0.3879, partial η^2^ = 0.0685, 95% CI of partial η^2^ = [0.0000–0.4071], BF_10_ = 0.20), or interaction between them (F_1,11_ = 0.2955, P = 0.5976, partial η^2^ = 0.0262, 95% CI of partial η^2^ = [0.0000–0.2207], BF_10_ = 0.06), indicating no significant learning in this group. These results indicate that VPL of a supra-threshold feature occurs as a result of exposure to the task-irrelevant NS images, in contrast to no performance improvement as a result of exposure to the dominant orientation in the task-irrelevant FS images.

In Experiment 1, VPL of orientation for supra-threshold NS images occurred, whereas that for FS images did not. This raises a question: is this tendency confined to VPL of orientation, or is it generalized to other features? To address this question, in Experiment 2 we focused on spatial frequency, a key feature underlying the perception of contours, textures, and spatial layouts in natural scenes^26,44–46^. Two new groups of 12 participants each (NS and FS groups) were trained and tested using the same procedures as in Experiment 1, with the following exceptions. First, from the NS and FS images used in Experiment 1, we selected images dominated by lower and higher spatial frequencies (see Experiment 2 in Methods for details and Fig. S4 for frequency spectra). Within each group, half of the participants were exposed to the lower-spatial-frequency dominant images and the other half to the higher-spatial-frequency dominant images during the exposure+task stage (see Fig. S1b for RSVP performance). Second, to assess VPL of spatial frequency, we employed a spatial frequency discrimination task (Fig. 2a) in the pre- and post-test stages (see Experiment 2 in Methods for details).

**Fig. 2:**
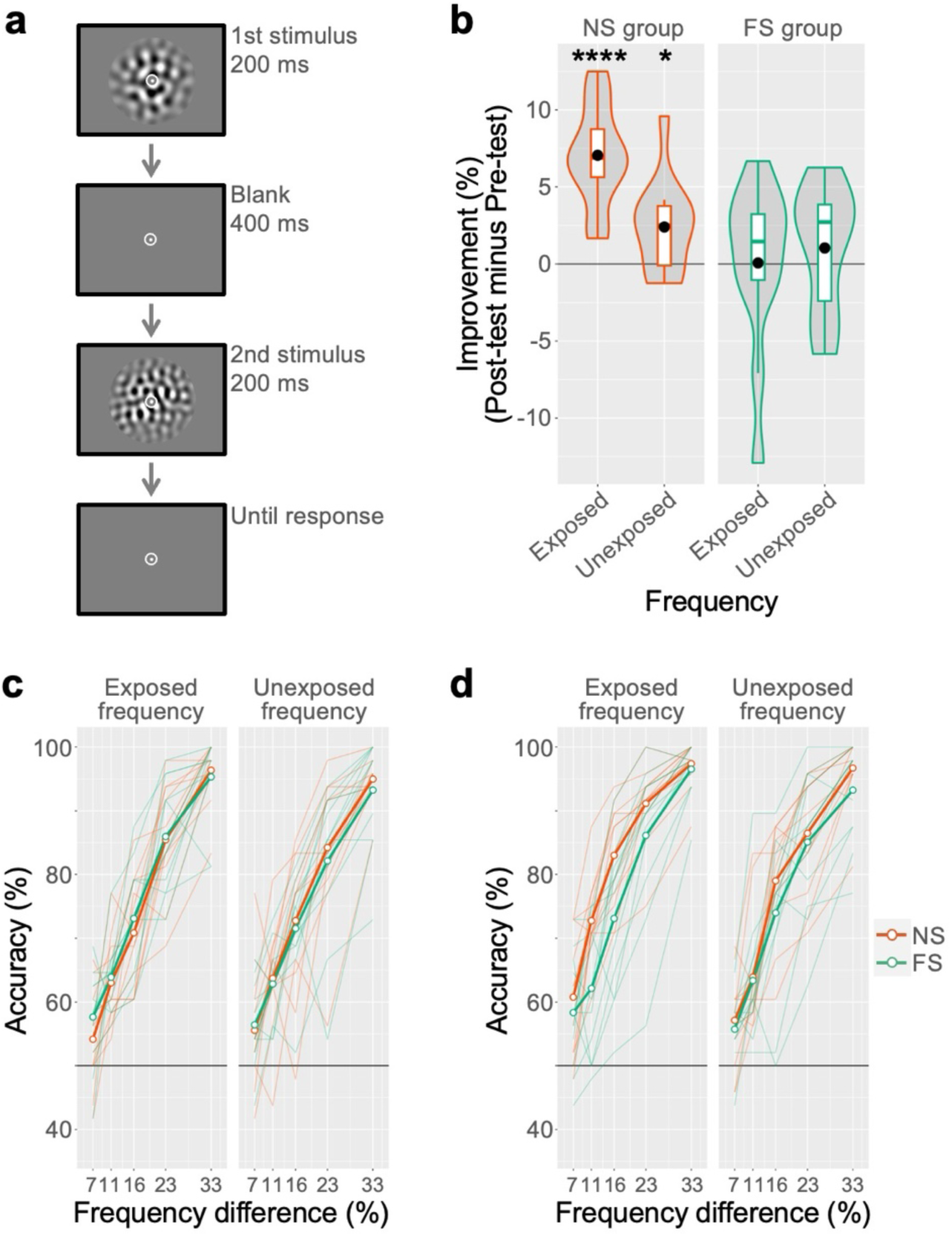
Methods and results of Experiment 2 **a**, Procedure of the spatial frequency discrimination task in the pre-test and post-test stages. Participants reported whether the spatial frequencies of the first and second stimuli were the same or different. **b**, Improvement in performance on the frequency discrimination task (post-test minus pre-test) for the NS (left panel) and FS (right panel) groups. For the NS group, a two-way repeated-measures ANOVA revealed a significant effect of Test (pre vs post) and significant interaction between Test and Frequency (exposed vs unexposed) (see Table S1 for detailed results). Subsequent analyses showed a significant simple effect of Test at the exposed (F_1,11_ = 57.5034, P < 10^-4^, partial η^2^ = 0.8394, 95% CI of partial η^2^ = [0.6791–0.9249], BF_10_ = 2013.4405) and unexposed (F_1,11_ = 7.5999, P = 0.0187, partial η^2^ = 0.4086, 95% CI of partial η^2^ = [0.0685–0.6695], BF_10_ = 3.5824) frequencies. For the FS group, a two-way repeated-measures ANOVA showed no significant effects. Box plots are overlaid on violin plots. Black dots indicate the mean accuracies across participants, and violin plots show kernel probability densities of individual accuracies. *P < 0.05, ****P < 0.001. **c**, Performance on the frequency discrimination task in the pre-test stage. **d**, Performance in the post-test stage. Thick lines indicate mean accuracy across participants, and thin lines represent individual accuracies.

Fig. 2b shows performance improvements in both the NS and FS groups (see Fig. 2c and d for task accuracy prior to the averaging and subtraction). We applied a three-way mixed-model ANOVA to performance with Group (NS vs FS) as the between-participant factor and Test (pre vs post) and Frequency (exposed vs unexposed) as the within-participant factors (see Table S1 for detailed results of ANOVAs). The analysis revealed a significant three-way interaction (F_1,22_ = 7.2760, P = 0.0132, partial η^2^ = 0.2485, 95% CI of partial η^2^ = [0.0029–0.4819], BF_10_ = 1.43), prompting a separate two-way ANOVA for each group. In the NS group, the results of the ANOVA showed a significant effect of Test (F_1,11_ = 37.2494, P = 0.0001, partial η^2^ = 0.7720, 95% CI of partial η^2^ = [0.5789–0.8790], BF_10_ > 10^4^) and significant interaction between Test and Frequency (F_1,11_ = 25.6514, P = 0.0004, partial η^2^ = 0.6999, 95% CI of partial η^2^ = [0.4495–0.8250], BF_10_ = 21.83). Subsequently, we found a significant simple effect of Test for the exposed (F_1,11_ = 57.5034, P < 10^-4^, partial η^2^ = 0.8394, 95% CI of partial η^2^ = [0.6791–0.9249], BF_10_ = 2013.4405) and unexposed orientations (F_1,11_ = 7.5999, P = 0.0187, partial η^2^ = 0.4086, 95% CI of partial η^2^ = [0.0685–0.6695], BF_10_ = 3.5824). In contrast, the FS group showed no significant effect of Test (F_1,11_ = 0.2977, P = 0.5962, partial η^2^ = 0.0264, 95% CI of partial η^2^ = [0.0000–0.2475], BF_10_ = 0.18) or Frequency (F_1,11_ = 1.3768, P = 0.2654, partial η^2^ = 0.1112, 95% CI of partial η^2^ = [0.0000–0.5819], BF_10_ = 0.35), nor a interaction between them (F_1,11_ = 0.2697, P = 0.6138, partial η^2^ = 0.0239, 95% CI of partial η^2^ = [0.0000–0.1508], BF_10_ = 0.08), indicating no evidence of learning in this group.

These results indicate that the result of Experiment 1̶VPL of orientation in NS and FS images being task-irrelevant̶is generalized to that of spatial frequency, another fundamental visual feature.

### Self-paced passive viewing enables unsupervised VPL for both natural scene and artificial images

Do the results of Experiments 1 and 2 mean that unsupervised VPL does not occur with supra-threshold artificial images? Note that in statistical learning literature, a number of studies demonstrated unsupervised learning based on passive viewing paradigms, where participants are allowed to freely observe presented stimuli without performing any given task^5–8,10^. One possible explanation for the occurrence of unsupervised learning in these prior studies is that passive viewing paradigms may not induce attentional suppression of the presented stimuli, whereas such suppression occurs with task-irrelevant images presented during active task performance to support efficient task execution^16^.

To test this possibility, we conducted Experiment 3 (Fig. 3a). Twenty-four new participants were trained and tested in the same way as in Experiment 1, with one exception. The exposure stage in Experiment 1 was replaced with an exposure stage in which participants were instructed to maintain central fixation but were allowed to freely view the presented images in a self-paced manner (Fig. 3b). Note that in this case presented images were not task-irrelevant since no task was performed during presentation of these images. Half of the participants were shown NS images (NS group), while the other half were shown the FS images (FS group).

**Fig. 3:**
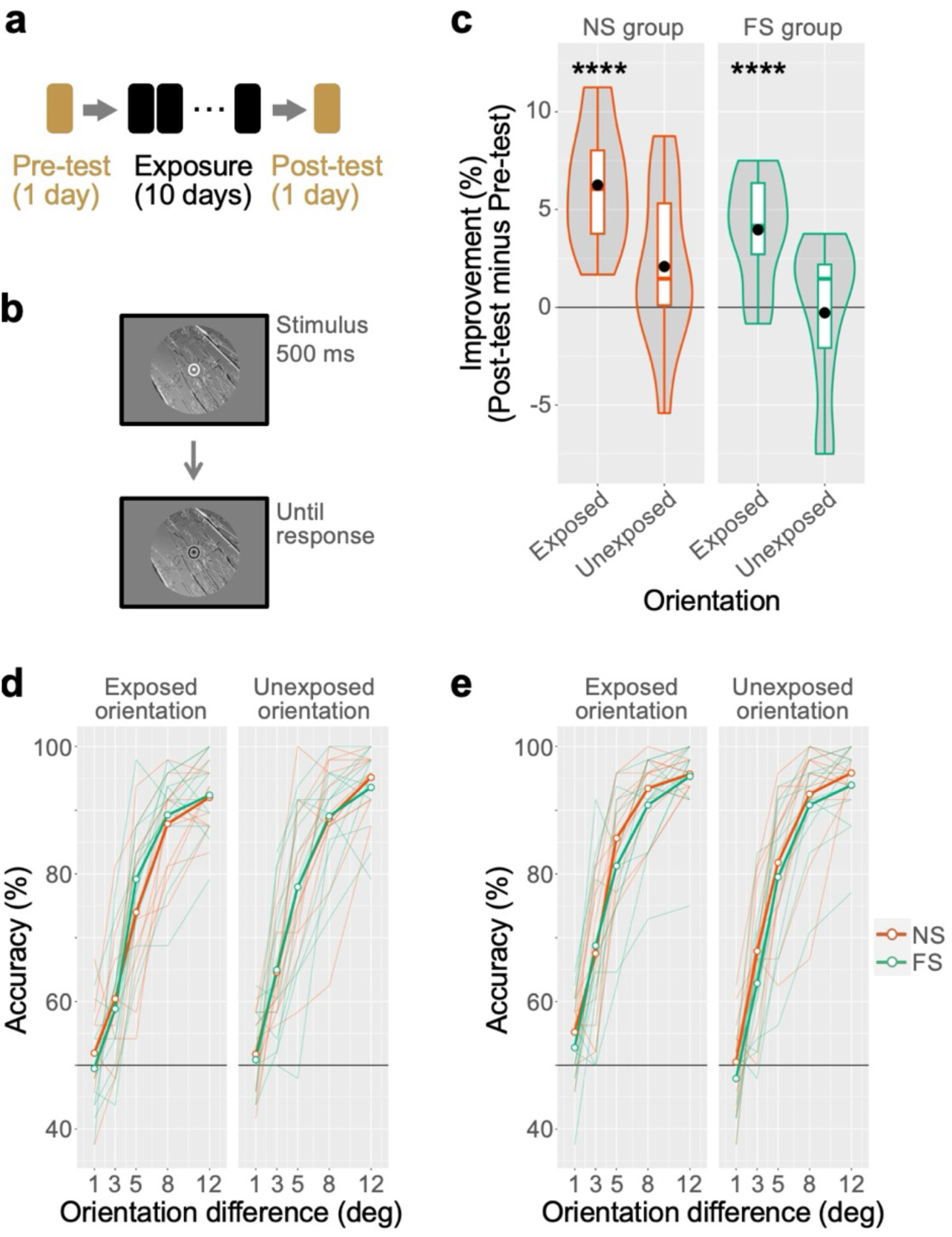
Methods and results of Experiment 3 **a**, Experiment 3 involved a 10-day exposure stage with self-paced passive viewing, instead of the exposure+task stage as in Experiment 1. The pre- and post-test stages were identical to those in Experiment 1. **b**, Procedure of a single trial during the exposure stage. Participants were instructed to observe an image at their own pace, pressing a button to proceed to the next trial once ready. **c**, Improvement in performance on the orientation discrimination task (post-test minus pre-test) for the NS (left panel) and FS (right panel) groups. A three-way mixed-model ANOVA on performance revealed significant effects of Test (pre vs post) and a two-way interaction between Test and Orientation (exposed vs unexposed), but no main effect of Orientation (see Table S1 for details). Subsequent analyses showed a significant simple effect of Test for the exposed orientation (F_1,22_ = 72.2270, P < 10^-4^, partial η^2^ = 0.7665, 95% CI of partial η^2^ = [0.6351–0.8543], BF_10_ > 10^5^), but not for the unexposed orientation (F_1,22_ = 1.2694, P = 0.2720, partial η^2^ = 0.0546, 95% CI of partial η^2^ = [0.0000–0.3333], BF_10_ = 0.37). Box plots are overlaid on violin plots. Black dots indicate mean performance improvements across participants. The violin plots show kernel probability densities of individual performance improvements. ****P < 10^-4^. **d**, Performance in the orientation discrimination task in the pre-test stage. **e**, Performance in the post-test stage. Thick lines indicate the mean accuracy across participants, and thin lines represent individual accuracies.

Fig. 3c shows improvements in both NS and FS groups (see Fig. 3d and e for task accuracy prior to the averaging and subtraction). A three-way ANOVA on performance revealed a significant effect of Test (F_1,22_ = 27.6724, P < 10^-4^, partial η^2^ = 0.5571, 95% CI of partial η^2^ = [0.2318–0.7195], BF_10_ > 10^4^) and significant interaction between Test and Orientation (F_1,22_ = 25.1603, P = 0.0001, partial η^2^ = 0.5335, 95% CI of partial η^2^ = [0.2530–0.7127], BF_10_ = 41.17) (see Table S1 for details). We found a significant simple effect of Test for the exposed orientation (F_1,22_ = 72.2270, P < 10^-4^, partial η^2^ = 0.7665, 95% CI of partial η^2^ = [0.6351–0.8543], BF_10_ > 10^5^), but not for the unexposed orientation (F_1,22_ = 1.2694, P = 0.2720, partial η^2^ = 0.0546, 95% CI of partial η^2^ = [0.0000–0.3333], BF_10_ = 0.37). These results indicate that unsupervised VPL of a supra-threshold feature occurs with both NS and FS images under a passive viewing protocol.

### Testing if marginal statistics (kurtosis and skewness) in natural scene images as task-irrelevant play a critical role in unsupervised VPL

Then, an important question arises: What enables unsupervised VPL of the dominant orientation in task-irrelevant natural scene images to occur under attentional suppression? To address this question, we further manipulated characteristics of images presented as task-irrelevant while participants engaged in the RSVP task during the exposure+task stage in the subsequent experiments (Experiments 4-7).

In Experiment 4, we tested if marginal statistics such as kurtosis and skewness, which are among distinctive characteristics of natural scene images that might enable such learning. Unlike the artificial images used in Experiment 1, which preserve lower-order statistics but lack structural regularities, and unlike higher-order statistics that capture complex spatial relationships such as edges, textures, and contours typical of natural scenes, kurtosis and skewness do not account for these rich spatial patterns^26,29,56^. It is possible that VPL of FS images as task-irrelevant does not occur because it does not have the same kurtosis and skewness in the luminance histogram as in NS images. If true, we should expect exposures to task-irrelevant FS images with the same kurtosis and skewness as NS images, referred to as kurtosis- and skewness-matched (KS) images (Fig. 4a, highlighted with purple rings), to lead to VPL of the dominant orientation in these images.

**Fig. 4:**
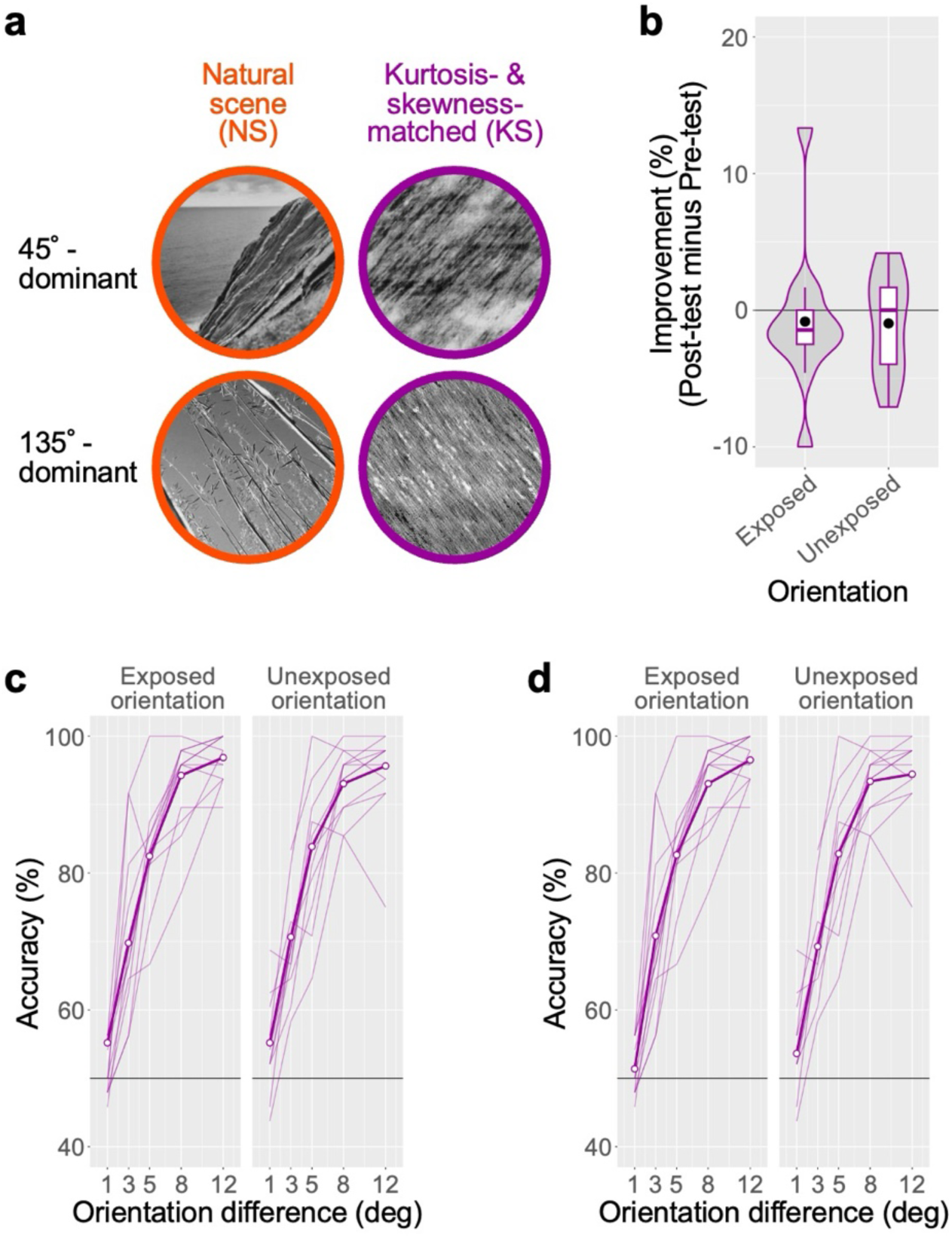
Methods and results of Experiment 4 **a**, Examples of the NS images (highlighted with orange rings) and kurtosis- & skewness-matched images (KS, highlighted with purple rings). **b**, Improvement in performance on the orientation discrimination task (post-test minus pre-test) for the KS group. A two-way repeated-measures ANOVA revealed no significant effects of Test (pre vs post), Orientation (exposed vs unexposed), or their interaction (see Table S1 for details). Box plots are overlaid on violin plots. Black dots indicate mean performance improvements across participants, while violin plots show kernel probability densities of individual performance improvements. **c**, Performance on the orientation discrimination task in the pre-test stage. **d**, Performance in the post-test stage. Thick lines indicate mean accuracy across participants, and thin lines represent individual accuracies.

In Experiment 4, 12 new participants were trained and tested in the same way as in Experiment 1, with the following exception. During the exposure+task stage (see Fig. S1c for RSVP performance), the participants were exposed to KS images (KS group) as task-irrelevant. The KS images consisted of similar local orientation and spatial frequency spectra as NS images with kurtosis and skewness that were matched to the NS images as discussed above. Fig. 4b shows performance improvements in the KS group (see Figs. 4c and d for task accuracies before the averaging and subtraction). Results of a two-way ANOVA on performance with Test (pre vs post) and Orientation (exposed vs unexposed) showed no significant effect of Test (F_1,11_ = 0.6535, P = 0.4360, partial η^2^ = 0.0561, 95% CI of partial η^2^ = [0.0000–0.4466], BF_10_ = 0.25), Orientation (F_1,11_ = 0.0234, P = 0.8812, partial η^2^ = 0.0012, 95% CI of partial η^2^ = [0.0000–0.0159], BF_10_ = 0.16), or interaction between them (F_1,11_ = 0.0092, P = 0.9254, partial η^2^ = 0.0008, 95% CI of partial η^2^ = [0.0000–0.0052], BF_10_ = 0.05). These results indicate no significant learning as a result of exposure to the task-irrelevant KS images.

### Testing if higher-order statistics in natural scene images as task-irrelevant play a critical role in unsupervised VPL

The results of Experiment 4 indicate that it is not the marginal statistics such as kurtosis or skewness that play a role in VPL of the dominant orientation presented as task-irrelevant. A growing body of research suggests that higher-order statistics derived from natural scenes are essential for perceiving and recognizing the natural scenes^57^. The higher-order statistics include *correlations* between different local orientation signals and spatial frequency signals^57–59^. In contrast, lower-order and marginal statistics include local orientation and spatial frequency signals and luminance histogram that constitute the NS as discussed above. The Portilla-Simoncelli (PS) images^27^ (Fig 5a, highlighted with blue rings; see Stimuli in Methods for details of creating the PS images) include the higher-order statistics derived from the NS images used in Experiment 1. If the higher-order statistics derived from NS images are necessary for VPL of the dominant orientation in these task-irrelevant images, then exposure to the PS images as task-irrelevant should result in VPL. In contrast, there was no evidence that VPL occurred with the KS images as task-irrelevant in Experiment 4, likely because they lack these higher-order statistical signals. Thus, in Experiment 5, to test whether higher-order statistics are necessary for VPL of the dominant orientation in NS images, we examined whether exposure to task-irrelevant supra-threshold PS images derived from these NS images would lead to VPL of that orientation. The procedures were identical to Experiment 1, except that 12 new participants were exposed to the PS images as task-irrelevant during the exposure+task stage (PS group; see Fig. S1d for RSVP performance).

**Fig. 5:**
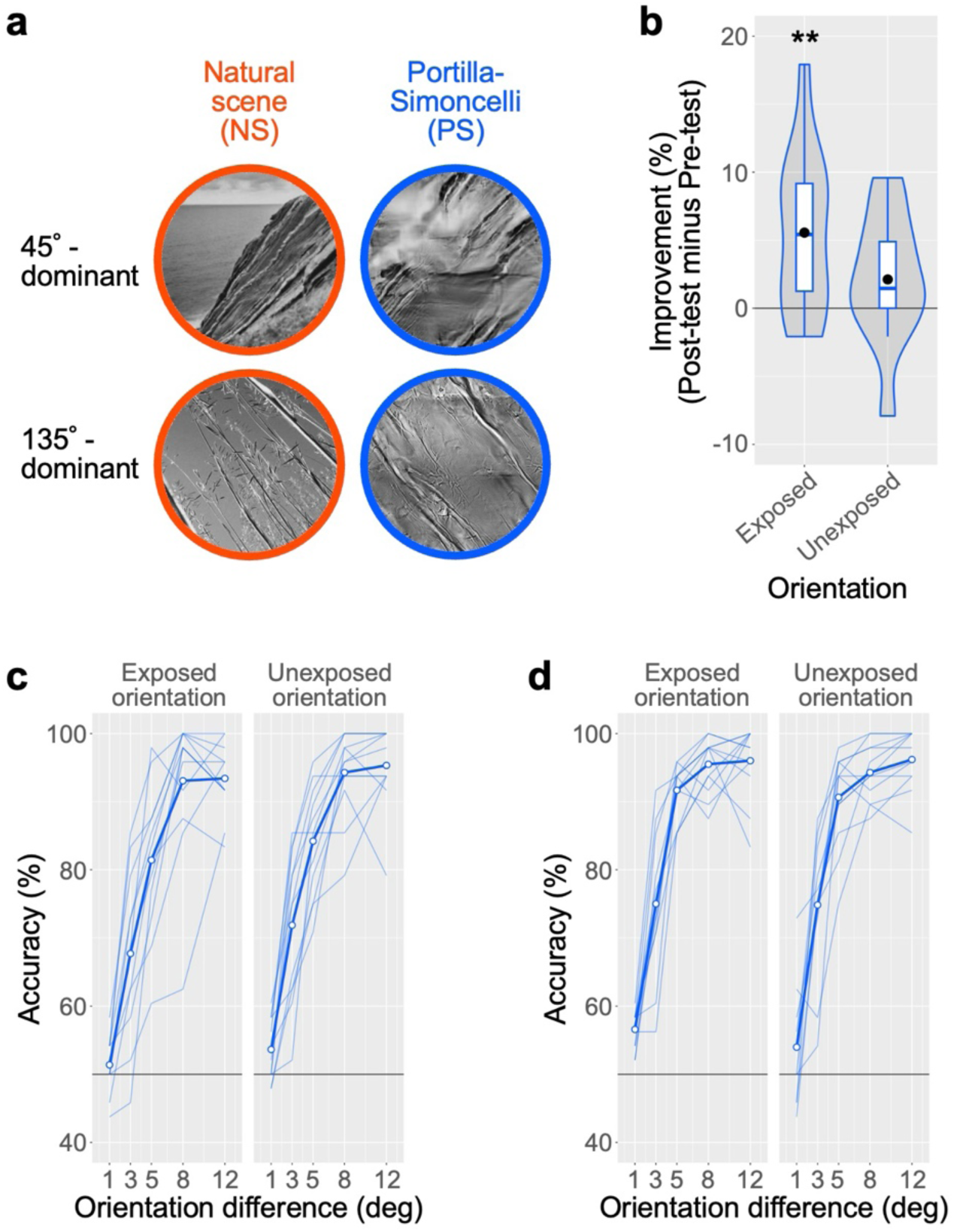
Methods and results of Experiment 5 **a**, Examples of the NS images (highlighted with orange rings) and Portilla–Simoncelli images (PS, highlighted with blue rings). **b**, Improvement in performance on the orientation discrimination task (post-test minus pre-test) for the PS group. A two-way repeated-measures ANOVA revealed a significant interaction between Test (pre vs post) and Orientation (exposed vs unexposed) (see Table S1 for details). Subsequent analyses showed a significant simple effect of Test at the exposed orientation (F_1,11_ = 10.5579, P = 0.0077, partial η^2^ = 0.4897, 95% CI of partial η^2^ = [0.1926–0.7240], BF_10_ = 7.286), but not at the unexposed orientation (F_1,11_ = 2.3348, P = 0.1547, partial η^2^ = 0.1751, 95% CI of partial η^2^ = [0.0000–0.5633], BF_10_ = 0.725). Box plots are overlaid on violin plots. Black dots indicate mean performance improvements across participants, and violin plots show kernel probability densities of individual performance improvements. **P < 0.01. **c**, Performance in the orientation discrimination task in the pre-test stage. **d**, Performance in the post-test stage. Thick lines indicate mean accuracy across participants, and thin lines represent individual accuracies.

Fig. 5b shows performance improvements in the PS group (see Figs. 5c and d for task accuracies before the averaging and subtraction). Results of a two-way ANOVA on performance with Test (pre vs post) and Orientation (exposed vs unexposed) showed significant effects of Test (F_1,11_ = 7.0718, P = 0.0222, partial η^2^ = 0.3913, 95% CI of partial η^2^ = [0.0298–0.7107], BF_10_ = 56.28) and significant interaction between Test and Orientation (F_1,11_ = 8.6700, P = 0.0133, partial η^2^ = 0.4408, 95% CI of partial η^2^ = [0.1225–0.6047], BF_10_ = 0.89), suggesting significant learning. We found a significant simple effect of Test for the exposed dominant orientation (F_1,11_ = 10.5579, P = 0.0077, partial η^2^ = 0.4897, 95% CI of partial η^2^ = [0.1926–0.7240], BF_10_ = 7.286), but not for the unexposed orientation (F_1,11_ = 2.3348, P = 0.1547, partial η^2^ = 0.1751, 95% CI of partial η^2^ = [0.0000–0.5633], BF_10_ = 0.725), demonstrating VPL as a result of exposure to the PS images as task-irrelevant.

The PS images consisted of the higher-order, marginal, and lower-order statistics, derived from NS images. Therefore, the above results indicate that PS images, consisting of these higher-order, marginal, and lower-order statistics, are sufficient to induce unsupervised VPL. We found the absence of VPL with the task-irrelevant KS images, which consisted of the lower-order and marginal statistics. Thus, if unsupervised VPL of the dominant orientation occurs only with the higher-order and marginal statistics in PS images, we may as well say that the higher-order statistics alone are sufficient for unsupervised VPL to occur.

To test if this is true, for the first time, we created images termed a higher-order statistics (HS) image, only consisting of the marginal and higher-order statistic signals derived from NS images (Fig. 6a; see Fig. S5 and Stimuli in Methods for details). In the HS image, the local orientation component of the original NS image was practically taken away by randomizing these orientations (see Fig. S6 for orientation power distribution of HS images).

**Fig. 6:**
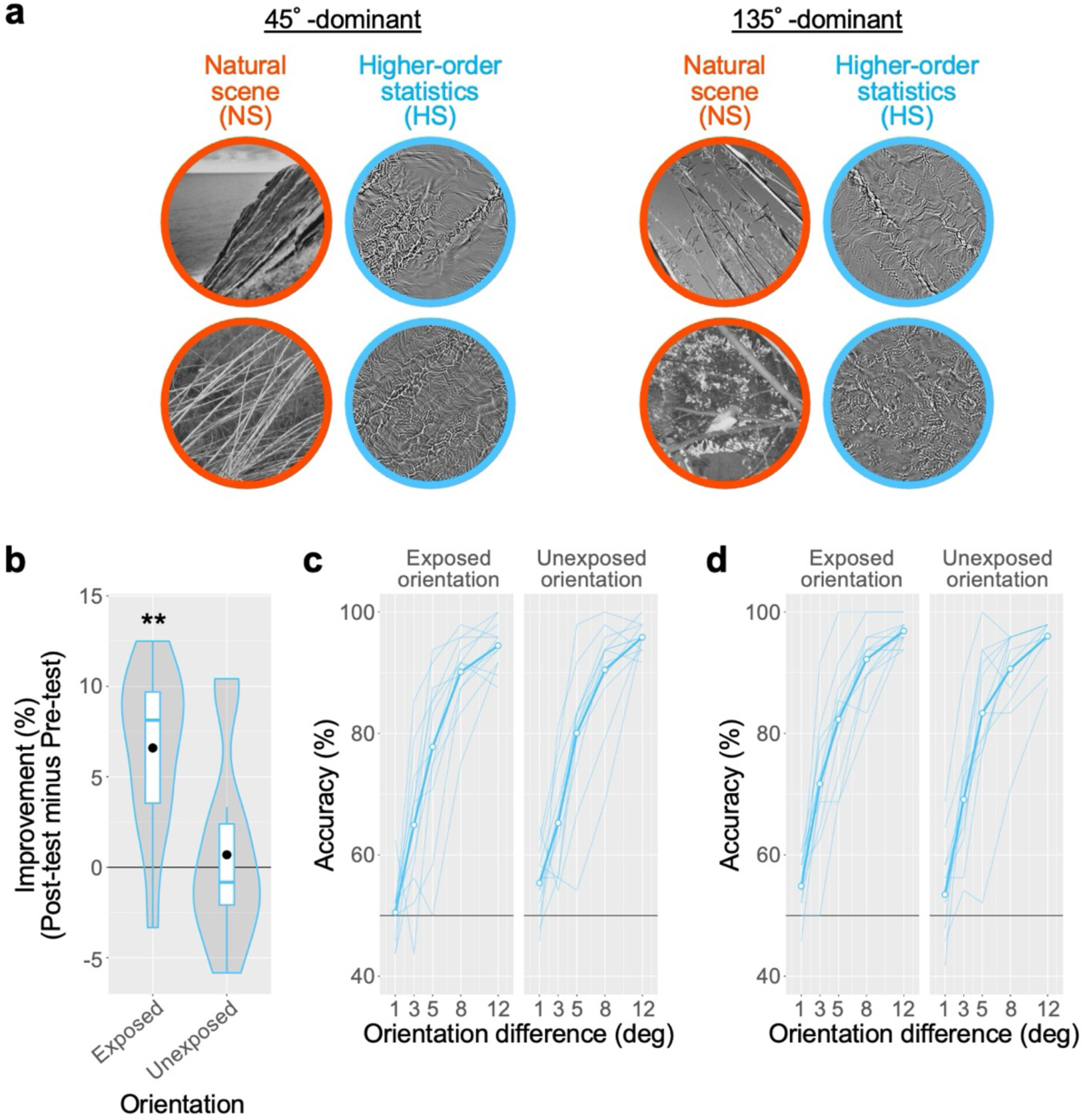
Methods and results of Experiment 6 **a**, Examples of the NS images (highlighted with orange rings) and higher-order statistics images (HS; highlighted with cyan rings). The HS images were generated to match the marginal and higher-order statistics of the NS images (see Stimuli in Methods for details). Specifically, we first computed higher-order statistics of the NS images, then randomly shuffled their pixels to create images that preserved no spatial structure. Next, we calculated the lower-order and marginal statistics from these shuffled images. Finally, we synthesized the HS images to match the computed higher-order, marginal, and lower-order statistics. See Fig. S5 for detailed graphical illustrations. **b**, Improvement in performance on the orientation discrimination task (post-test minus pre-test) for the HS group. A two-way repeated-measures ANOVA revealed significant effects of Test (pre vs post) and an interaction between Test and Orientation (exposed vs unexposed) (see Table S1 for details). Subsequent analyses showed a significant simple effect of Test at the exposed orientation (F_1,11_ = 15.9225, P = 0.0021, partial η^2^ = 0.5914, 95% CI of partial η^2^ = [0.2705–0.7647], BF_10_ = 21.28), but not at the unexposed orientation (F_1,11_ = 2.6466, P = 0.1320, partial η^2^ = 0.1939, 95% CI of partial η^2^ = [0.0003–0.5629], BF_10_ = 0.81). Box plots are overlaid on violin plots. Black dots indicate mean performance improvements across participants. Violin plots show kernel probability densities of individual performance improvements. **P < 0.01. **c**, Performance in the orientation discrimination task in the pre-test stage. **d**, Performance in the post-test stage. Thick lines indicate mean accuracy across participants, and thin lines represent individual accuracies.

Thus, in Experiment 6, we tested whether VPL of the dominant orientation occurs as a result of exposure to the HS images as task-irrelevant. The procedures were identical to Experiment 1, except that 12 new participants were exposed to the HS images as task-irrelevant during the exposure+task stage (see Fig. S1e for performance in the RSVP and orientation discrimination tasks). We applied a two-way ANOVA on performance with the factors of Test (pre-vs post-tests) and Orientation (exposed vs unexposed orientations) and found a significant effect of Test (F_1,11_ = 10.8027, P = 0.0072, partial η^2^ = 0.4955, 95% CI of partial η^2^ = [0.1096–0.7385], BF_10_ = 23.63) and significant interaction between Test and Orientation (F_1,11_ = 16.1700, P = 0.0020, partial η^2^ = 0.5951, 95% CI of partial η^2^ = [0.3204–0.7699], BF_10_ = 1.24; see Table S1 for details), suggesting learning specific to the exposed orientation (Fig. 6b; see Figs. 6c and d for task accuracies before the averaging and subtraction). We also found a significant simple effect of Test at the exposed orientation (F_1,11_ = 15.9225, P = 0.0021, partial η^2^ = 0.5914, 95% CI of partial η^2^ = [0.2705–0.7647], BF_10_ = 21.28), but not at the unexposed orientation (F_1,11_ = 2.6466, P = 0.1320, partial η^2^ = 0.1939, 95% CI of partial η^2^ = [0.0003–0.5629], BF_10_ = 0.81), indicating unsupervised VPL of a dominant orientation with the HS images. Notably, the HS images that did not contain components of lower-order statistics from original natural scene images (Fig. S6). This result supports the hypothesis that the higher-order statistics alone generate the dominant orientation based on which unsupervised VPL of the orientation can occur.

In Experiments 1 and 4, we found no significant VPL for the global orientation in artificial stimuli such as the FS and KS images presented as task-irrelevant during the RSVP task. This suggests that the global orientation information in lower-order and/or marginal statistics is suppressed by attention, leading to no VPL. It is possible that the failure to observe VPL was due to the task-irrelevant FS and KS images lacking sufficient power to induce VPL of the global orientations (e.g., 45° and 135°) assessed in pre- and post-tests.

To test this possibility, we conducted a new experiment using stimuli that had clear orientation peaks at 45 and 135 degrees (see Fig. S7a and Stimuli in Methods for details). The procedures were identical to Experiment 1, except that 12 new participants were exposed to Gabor patch (GP) images (GP group), which had prominent orientation power at 45 or 135 degrees, as task-irrelevant during the exposure+task stage (see Fig. S1f for RSVP performance).

Fig. S7b appears to show no performance improvements in the GP group (see Figs. S7c and d for task accuracies before the averaging and subtraction). To confirm the observation, a two-way repeated-measures ANOVA on performance with Test (pre vs post) and Orientation (exposed vs unexposed) was conducted. The results showed no significant effect of Test (F_1,11_ = 0.0293, P = 0.8671, partial η^2^ = 0.0027, 95% CI of partial η^2^ = [0.0000–0.0221], BF_10_ = 0.16), Orientation (F_1,11_ = 0.8781, P = 0.3689, partial η^2^ = 0.0739, 95% CI of partial η^2^ = [0.0000–0.4179], BF_10_ = 0.23), or interaction between them (F_1,11_ = 1.1918, P = 0.2983, partial η^2^ = 0.0978, 95% CI of partial η^2^ = [0.0001–0.4709], BF_10_ = 0.07), showing no significant VPL as a result of exposure to the task-irrelevant GP images which have strong orientation power at the orientation to be tested. These results indicate that if an image has merely prominent orientation power but lacks higher-order statistics, it is not sufficient to induce VPL of a task-irrelevant orientation.

In Experiments 1-7, six different types of images—NS, FS, KS, PS, HS, and GP—were exposed as task-irrelevant. The characteristics of each type are shown in Table 1. The summary of the results (Table 2) indicates that exposures to task-irrelevant images that contain the higher-order statistics from NS images (NS, PS, and HS) led to VPL of the dominant orientation of the images under attentional suppression while exposures to task-irrelevant images lacking these higher-order statistics (FS, KS, and GP) did not.

**Table 1:**
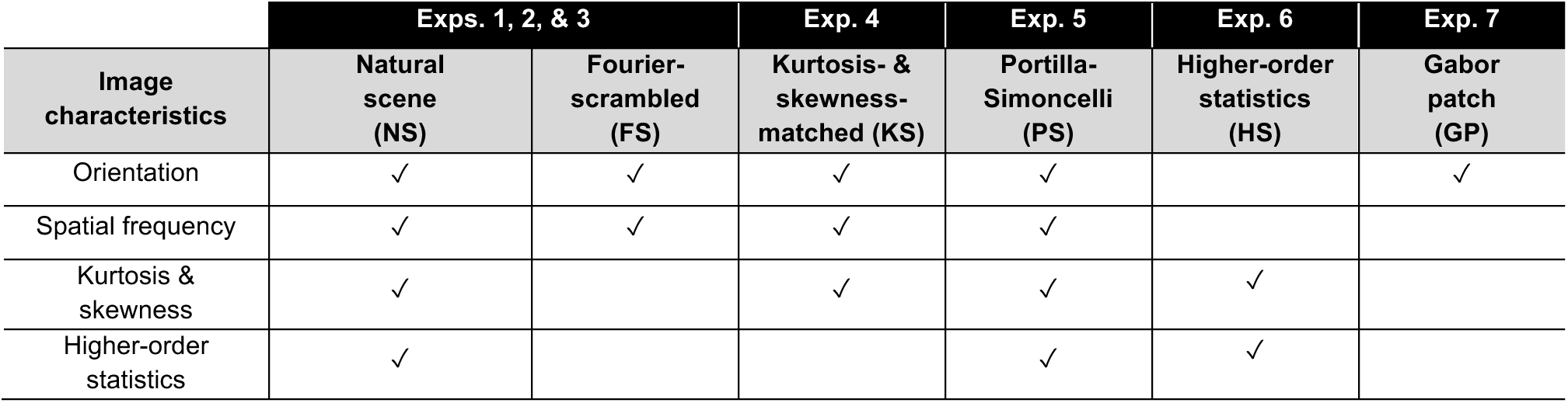
Image characteristics of 6 different image types in Experiments 1-7

**Table 2:**
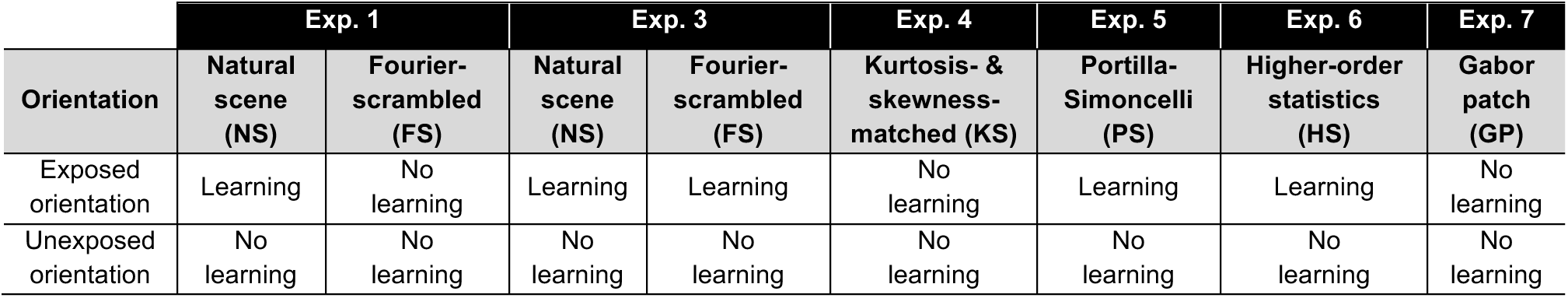
Summary results of Experiments 1, 3-7

Then, why does exposure to task-irrelevant images containing higher-order statistics from NS images lead to VPL even under conditions of attentional suppression? One possibility is that the dominant orientation generated from these higher-order statistics was sub-threshold and invisible, given that VPL of a task-irrelevant feature in an artificial stimulus occurs if the feature is sub-threshold^16,17,40,41^. That might have been why unsupervised VPL of the dominant orientation in the images containing these higher-order statistics occurred.

We conducted Experiment 8 to test whether the dominant orientation in the HS images, which contained higher-order statistics but not lower-order statistics, was sub-threshold or supra-threshold. A new group of 12 participants were asked to report whether the dominant orientation in HS images was rotated clockwise or counterclockwise from the vertical (See Experiment 8 in Methods for details). As shown in Fig. S3b, mean accuracy was significantly higher than the chance level for the 45°-dominant images (one-sample *t*-test with Bonferroni correction for multiple comparisons; t_11_ = 9.2317, corrected P < 10^-4^, Cohen’s d = 8.0307, 95% CI of accuracy = [68.9127–80.7540] %, BF_10_ = 10726.38) and the 135°-dominant images (t_11_ = 11.6124, corrected P < 10^-4^, Cohen’s d = 10.8153, 95% CI of accuracy = [68.2016–76.7151] %, BF_10_ = 83865.74). These results indicate that the HS images were supra-threshold, ruling out the possibility that the orientation in HS images used in Experiment 6 were sub-threshold, allowing VPL of the orientation to occur.

Performance in the orientation discrimination task was lower for the global orientation in HS images than for the NS and PS images (Fig. S3b). It has been found that task-irrelevant features that are at or close to threshold are not subject to attentional suppression^17,40,41^. These results collectively raise the possibility that participants were less likely to detect the orientation in the HS images than that in the other images, which may have influenced learning of the orientation of the task-irrelevant HS images. However, this possibility is unlikely. Fig. S3b shows that the orientation discriminability in the task-irrelevant HS images was much higher than threshold (BF_10_ > 10^4^; the lower bound of the confidence interval is 68.9%). Therefore, it is unlikely that the lower orientation discriminability in the task-irrelevant HS images significantly influenced the manner in which VPL in the HS images was developed.

In Experiment 6, it is possible that the lower visibility of the HS images reduced suppression by reducing competition with the RSVP stream. If this were the case, the KS images whose visibility was equivalent to that of HS images should also be subject to reduced suppression. To test the possibility, we conducted Experiment 9 using the KS images with reduced visibility (low-visibility KS images, Fig. S8a; see Experiment 9 in Methods for details). The low-visibility KS images were generated using a pixel randomization method (see Experiment 9 in Method) so that performance in the orientation discrimination task using these images (Fig. S8b) matched that with the HS images (73.7%; Fig. S3b). The procedures of Experiment 9 were identical to those of Experiment 4, except that 12 new participants were exposed to the low-visibility KS images during the exposure+task stage (see Fig. S1g for RSVP performance). We applied a two-way repeated-measured ANOVA to performance (see Fig. S8c for performance improvements and Fig. S8d and e for task accuracy prior to the averaging and subtraction) with the factors of Test (pre vs post) and Orientation (exposed vs unexposed). The analysis revealed no significant effect of Test (F_1,11_ = 1.1175, P = 0.3131, partial η^2^ = 0.0922, 95% CI of partial η^2^ = [0.0000–0.4081], BF_10_ = 0.38) or Orientation (F_1,11_ = 0.0828, P = 0.7789, partial η^2^ = 0.0075, 95% CI of partial η^2^ = [0.0000–0.0722], BF_10_ = 0.17), and no significant interaction between them (F_1,11_ = 1.8130, P = 0.2052, partial η^2^ = 0.1415, 95% CI of partial η^2^ = [0.0002–0.5720], BF_10_ = 0.10).

These results indicate that the lower visibility of the HS images does not account for their lack of susceptibility to attentional suppression in Experiment 6.

It has been reported that VPL tends to rely more on lower-level visual information as the task becomes more difficult^60^. In our study, accuracy changes (post-test minus pre-test) in the orientation discrimination task were obtained at five difficulty levels (Δ values of 12, 8, 5, and 1 degree; see Experiment 1 and Methods for details) and then averaged. Does exposure to NS, PS, or HS images—each containing higher-order statistics—produce differential learning gains across Δ = 12°, 8°, 5°, 1° that would signal a higher-order-specific effect?

To examine this, we compared accuracy changes at these five orientation differences (reflecting task difficulty) for NS, PS, and HS images, all of which included higher-order statistics (Fig. S11). We performed a three-way mixed-model ANOVA on accuracies, with Test (pre- vs post-test) and Difficulty level (12, 8, 5, and 1) as within-participant factors, and Image type (NS images in Experiment 1, PS images in Experiment 5, and HS images in Experiment 6) as the between-participant factor. The results showed that accuracy improved at all difficulty levels, but the patterns of improvement did not differ significantly across image types (see Fig. S11 and its legend for detailed analysis procedures). Accuracy improved rather uniformly across all Δ values and did not differ among NS, PS, and HS images. One possibility is that VPL induced by task-irrelevant images with higher-order statistics manifests as a general enhancement of low-level orientation processing rather than a change restricted to higher-order representations.

In Experiments 1-7 and 9, performance in the pre- and post-stages was defined as the mean accuracy averaged across the Δ values (see Experiments 1 and 2 in Methods for details). To validate the results based on this performance measure (Figs. 1e, 2b, 3c, 4b, 5b, 6b, 7b, 8b, and Table S1), we also calculated two alternative performance indices: the perceptual threshold (see Curve fitting in the Methods section for details) and d’. The same pattern of results was obtained when performance improvements were quantified using perceptual thresholds (see Fig. S9 for performance improvements, Table S2 for ANOVA details on perceptual thresholds, and Table S3 for ANOVA details on slope) and d’ values (see Fig. S10 for performance improvements, Table S4 for ANOVA details on d’ values, and Table S5 for ANOVA details on biases). These convergent results further support the conclusion that unsupervised VPL occurs as a result of exposure to task-irrelevant images containing the higher-order statistics of natural scenes.

### Higher-order statistics in natural scene images are less susceptible to attentional suppression

Studies have consistently shown that exposure to supra-threshold task-irrelevant artificial stimuli during task performance, including Gabor patches, does not lead to VPL due to attentional suppression on the stimuli^16,17,40,41,47–55^. It is possible that the orientation derived from these higher-order statistics from NS images is less susceptible to attentional suppression than the orientation derived from the lower-order statistics, as the processing of the former may be significantly different from that of the latter^61–63^. If true, the reduced effect of attentional suppression on the higher-order statistics may lead to VPL of task-irrelevant NS, PS, and HS images but not of FS, KS, or GP images.

To test this hypothesis, in Experiment 10, we used an established experimental paradigm designed to measure the effect of attentional suppression on task-irrelevant features^64,65^. Specifically, we measured participants’ performance in an orientation discrimination task under two conditions. In the focused attention condition, the participants focused their attention on the orientation of stimuli. In the diverted attention condition, the participants were unexpectedly required to report the orientation of stimuli while their attention was directed away from the orientation. The effects of attentional suppression were compared based on performance in the diverted attention and focused attention conditions. We also employed a third condition (RSVP condition; details below) to confirm that the participants indeed diverted their attention from the orientation of the stimuli in the diverted attention condition.

A new group of 12 participants underwent these three conditions (Fig. 7a; see Experiment 10 in Methods for details). In the focused attention condition, participants performed the orientation discrimination task on the dominant orientation in both KS and HS images. Since KS images include the lower-order and marginal statistics derived from NS images, while HS images include the higher-order and marginal statistics from these NS images, using these two image types is sufficient to compare the effects of attentional suppression on task-irrelevant orientations arising from lower-order versus higher-order statistics.

**Fig. 7:**
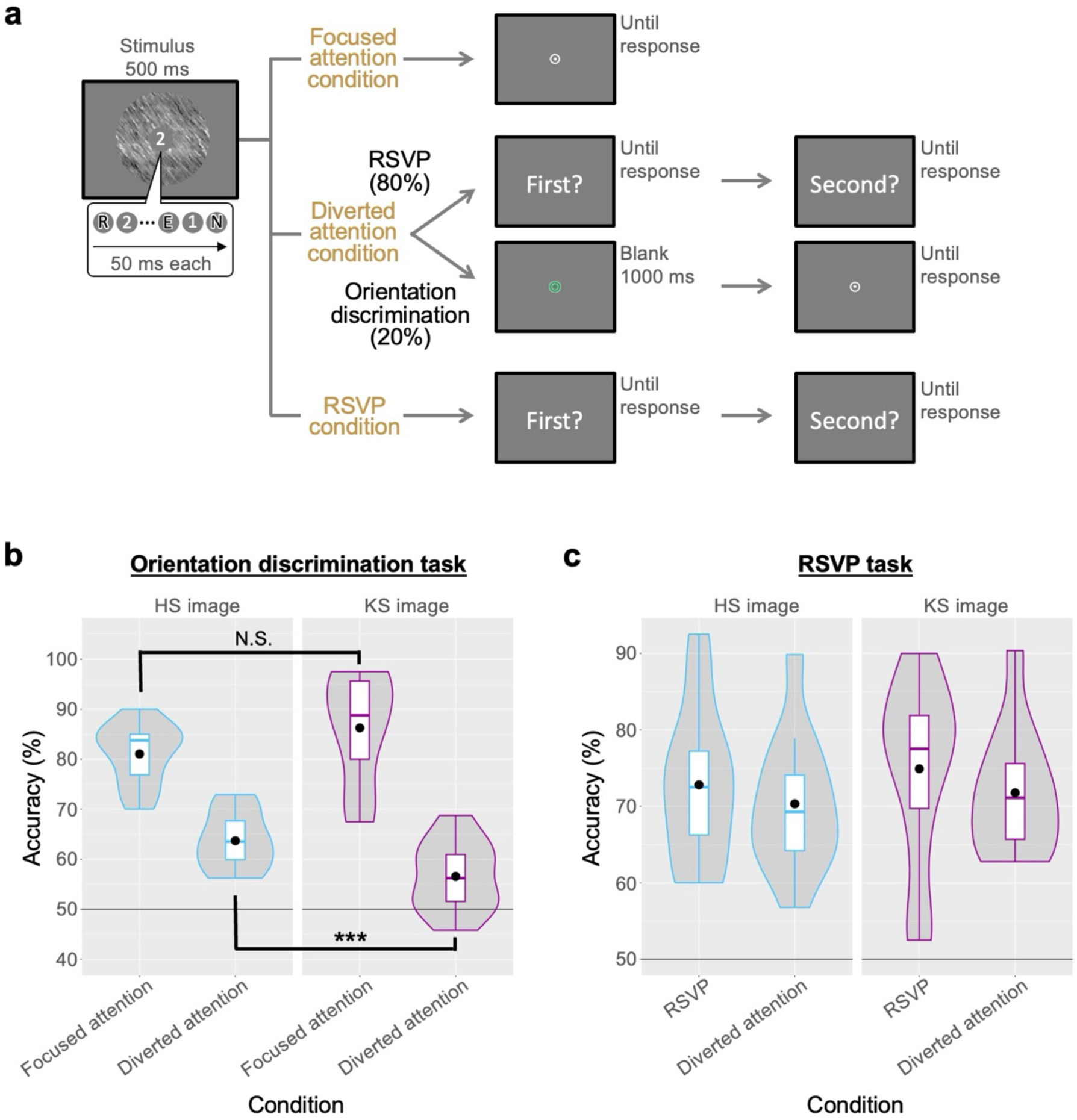
Procedures and results of Experiment 10 **a**, Illustrative timeline of trials in the focused attention, diverted attention, and RSVP conditions. In the focused attention condition, participants performed an orientation discrimination task, reporting whether the dominant orientation of a presented image was rotated clockwise or counterclockwise from the vertical. In the diverted attention condition, 80% of trials were assigned to an RSVP task, in which participants indicated two digits in a sequence of characters. In the remaining 20% of trials, they performed the orientation discrimination task. Notably, participants were informed of the task only after the stimulus offset and were instructed to prioritize the RSVP task over the orientation discrimination task. In the RSVP condition, participants performed only the RSVP task. **b**, Accuracies of the orientation discrimination task in the focused and diverted attention conditions. A two-way repeated-measures ANOVA on accuracies was conducted with the factors of Image (HS vs KS) and Condition (focused vs diverted attention). We found significant effects of Condition (F_1,11_ = 176.9310, P < 10^-4^, partial η^2^ = 0.9415, 95% CI of partial η^2^ = [0.8985–0.9662], BF_10_ < 10^11^) and an interaction between Image and Condition (F_1,11_ = 9.0928, P = 0.0118, partial η^2^ = 0.4525, 95% CI of partial η^2^ = [0.0366–0.7114], BF_10_ = 9.45), but no main effect of Image (F_1,11_ = 0.1774, P = 0.6818, partial η^2^ = 0.0159, 95% CI of partial η^2^ = [0.000–0.1390], BF_10_ = 2.10). Subsequent analyses revealed a significant simple effect of Image in the diverted attention condition (F_1,11_ = 15.5779, P = 0.0023, partial η^2^ = 0.5861, 95% CI of partial η^2^ = [0.1413–0.7705], BF_10_ = 19.99), but not in the focused attention condition (F_1,11_ = 1.7633, P = 0.2111, partial η^2^ = 0.1382, 95% CI of partial η^2^ = [0.0002–0.5646], BF_10_ = 0.59). **c**, Accuracies of the RSVP task in the RSVP and diverted attention conditions. A two-way repeated-measures ANOVA on accuracies was conducted with the factors of Image (HS vs KS) and Condition (RSVP vs diverted attention). None of main effects of Image (F_1,11_ = 2.0062, P = 0.1843, partial η^2^ = 0.1542, 95% CI of partial η^2^ = [0.0001–0.5296], BF_10_ = 0.29), Condition (F_1,11_ = 2.0800, P = 0.1771, partial η^2^ = 0.1590, 95% CI of partial η^2^ = [0.0007–0.6082], BF_10_ = 0.66), or their interaction (F_1,11_ = 0.0569, P = 0.8159, partial η^2^ = 0.0051, 95% CI of partial η^2^ = [0.0000–0.0511], BF_10_ = 0.15) was significant. Box plots are overlaid on violin plots. Black dots represent mean accuracies across participants, and violin plots show kernel probability densities of individual accuracies. ***P < 0.005.

In the diverted attention condition, they were asked to perform the RSVP task for 80% of the whole trials and orientation discrimination task for the remaining 20% of trials in a random order. At the beginning of each trial, participants were not informed of whether they would be asked to perform the RSVP or orientation discrimination task. The task for each trial was only revealed after the stimulus offset (see Experiment 10 in Methods for details). We instructed the participants to prioritize the RSVP task over the orientation discrimination task, ensuring that they diverted their attention away from the orientation at the time of the stimulus presentation. In the RSVP condition, the participants were asked to perform the RSVP task only. This RSVP condition was designed to confirm whether participants indeed prioritize the RSVP task in the diverted attention condition.

Fig. 7b shows accuracies in the orientation discrimination task for the focused attention and diverted attention conditions. A two-way repeated-measures ANOVA was applied to accuracy with the factors of Image (HS vs KS) and Condition (focused vs diverted attention). We found significant effects of Condition (F_1,11_ = 176.9310, P < 10^-4^, partial η^2^ = 0.9415, 95% CI of partial η^2^ = [0.8985–0.9662], BF_10_ < 10^11^) and an interaction between Image and Condition (F_1,11_ = 9.0928, P = 0.0118, partial η^2^ = 0.4525, 95% CI of partial η^2^ = [0.0366–0.7114], BF_10_ = 9.45), but no significant main effect of Image (F_1,11_ = 0.1774, P = 0.6818, partial η^2^ = 0.0159, 95% CI of partial η^2^ = [0.000–0.1390], BF_10_ = 2.10). Subsequent simple-effect analyses revealed a significant simple effect of Image in the diverted attention condition (F_1,11_ = 15.5779, P = 0.0023, partial η^2^ = 0.5861, 95% CI of partial η^2^ = [0.1413–0.7705], BF_10_ = 19.99), but not in the focused attention condition (F_1,11_ = 1.7633, P = 0.2111, partial η^2^ = 0.1382, 95% CI of partial η^2^ = [0.0002–0.5646], BF_10_ = 0.59). This finding supports the hypothesis that the supra-threshold dominant orientation in task-irrelevant HS images is less susceptible to attentional suppression compared to that in KS images. Notably, KS images comprise the lower-order and marginal statistics of NS images, while HS images comprise the marginal and higher-order statistics of the NS images. Furthermore, the results of Experiments 1 and 4-7 did not indicate any interaction in performance improvements across the lower-order, marginal, and higher-order statistics (see Table 2). Based on these findings, the results of Experiment 10 suggest that attentional suppression is less effective on task-irrelevant NS, PS, and HS, which include the higher-order statistics, than on FS, KS, and GP, which lack these statistics.

One possible concern is that the reduced effect of attentional suppression on the HS images might have resulted from improving their performance in the orientation discrimination task by deprioritizing the RSVP task in the diverted attention condition specifically when the HS images were presented in the background. If so, RSVP performance in the diverted attention condition with the HS images should be significantly lower than that in other conditions. To test this possibility, a two-way repeated-measures ANOVA was applied to accuracy (Fig. 7c) with the factors of Image (HS vs KS) and Condition (RSVP vs diverted attention). None of the main effects of Image (F_1,11_ = 2.0062, P = 0.1843, partial η^2^ = 0.1542, 95% CI of partial η^2^ = [0.0001–0.5296], BF_10_ = 0.29), Condition (F_1,11_ = 2.0800, P = 0.1771, partial η^2^ = 0.1590, 95% CI of partial η^2^ = [0.0007–0.6082], BF_10_ = 0.66), or their interaction (F_1,11_ = 0.0569, P = 0.8159, partial η^2^ = 0.0051, 95% CI of partial η^2^ = [0.0000–0.0511], BF_10_ = 0.15) was significant. These results indicate that the RSVP task was not deprioritized in the diverted attention condition when HS images were presented in the background.

Monkey studies have shown that the higher-order statistics derived from NS images are processed at V2 and higher areas, rather than in V1^32–34^. If neural activation representing the reduced effect of suppression on the task-irrelevant higher-order statistics indeed occurs as the above hypothesis indicates, the suppression should be observed in the areas corresponding to those observed in the monkey studies.

In Experiment 11, we tested this hypothesis. Specifically, we examined whether the degree of attentional suppression with HS images differs from that with KS images, using fMRI to measure blood-oxygen-level-dependent (BOLD) signals, while a new group of 28 participants was exposed to HS and KS images as task-irrelevant (Fig. 8a). During this measurement, participants were engaged in easy and harder RSVP tasks (Fig. 8a). It has been established that BOLD signals in response to task-irrelevant visual stimuli are more strongly suppressed during the harder RSVP task than the easy RSVP task^66,67^. We employed the easy and harder RSVP tasks to compare the effects of attentional suppression on BOLD responses to the task-irrelevant HS and KS images. The HS images are derived from the marginal and higher-order statistics of the NS images while the KS images are composed of the lower-order and marginal statistics of the NS images (see Table 1). Thus, differences in the effects of attentional suppression on BOLD signals between the task-irrelevant HS and KS images should reflect differences in neural processing related to the suppression on the higher- and lower-order task-irrelevant statistics in the brain. We performed five different analyses as below. Detailed experimental procedures are shown in Methods.

**Fig. 8:**
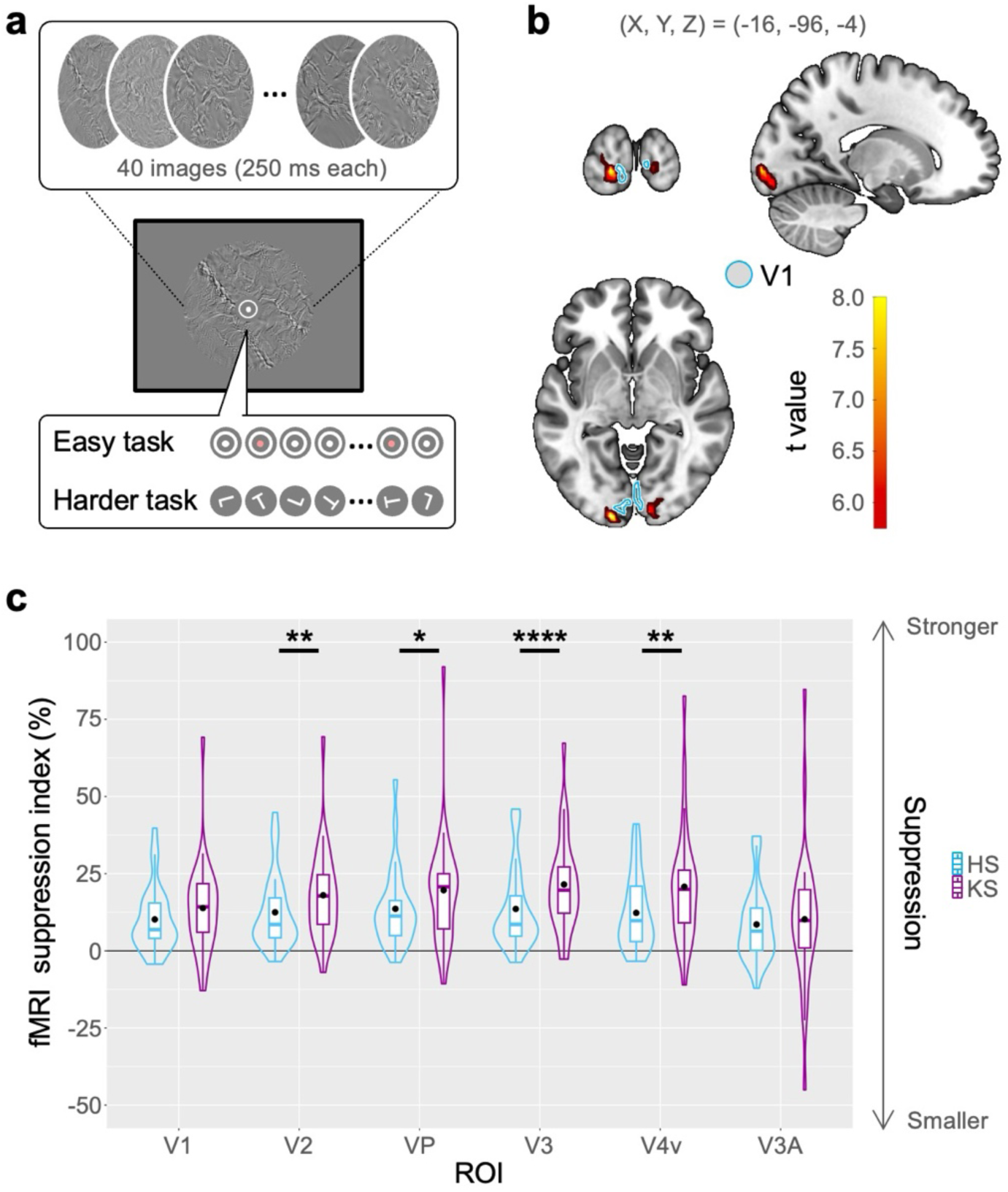
Procedures and results of Experiment 11 **a**, Illustrative timeline of the stimulus period during which participants performed easy and harder RSVP tasks while being exposed to HS and KS images as task-irrelevant (see Methods for details). In the easy RSVP task, participants indicated whether the number of presentations of a red fixation point was even or odd. In the harder RSVP task, they reported whether the number of presentations of a “T” was even or odd. **b**, Stronger responses to HS images than to KS images. Colored voxels show areas where BOLD signal amplitudes in response to task-irrelevant HS images were significantly higher than those to KS images (paired t-test, P < 0.05 after Bonferroni correction for multiple comparisons; see Methods for details). Regions enclosed by blue lines represent voxels identified as belonging to V1 in more than half of the 28 participants. **c**, The fMRI suppression indices for the task-irrelevant HS and KS images in each region of interest (ROI). The fMRI suppression index was defined as the percentage reduction in BOLD signal amplitudes during the harder RSVP task, compared to the easy RSVP task (see Methods for detailed calculation). A smaller index value indicates a smaller effect of attentional suppression on a BOLD response to a task-irrelevant image. A two-way repeated-measures ANOVA on fMRI suppression indices with factors of ROI (V1, V2, VP, V3, V4v, vs V3A) and Image (HS vs KS) revealed significant main effects of ROI (F_2.29,61.95_ = 6.5843, P = 0.0017, partial η^2^ = 0.1961, 95% CI of partial η^2^ = [0.0940–0.3243], BF_10_ = 821.36), Image (F_1,27_ = 7.0106, P = 0.0134, partial η^2^ = 0.2061, 95% CI of partial η^2^ = [0.0039–0.4220], BF_10_ > 10^4^), as well as a significant interaction between them (F_2.43,65.67_ = 3.2439, P = 0.0363, partial η^2^ = 0.1073, 95% CI of partial η^2^ = [0.0370–0.2135], BF_10_ = 0.13). Subsequent analyses revealed a significant simple effects of Image at V2 (F_1,27_ = 8.5033, P = 0.0071, partial η^2^ = 0.2395, 95% CI of partial η^2^ = [0.0224–0.4695], BF_10_ = 6.17), VP (F_1,27_ = 6.4939, P = 0.0168, partial η^2^ = 0.1939, 95% CI of partial η^2^ = [0.0090–0.4086], BF_10_ = 2.98), V3 (F_1,27_ = 20.5149, P = 0.0001, partial η^2^ = 0.4318, 95% CI of partial η^2^ = [0.1696–0.6453], BF_10_ = 245.80), and V4v (F_1,27_ = 8.6490, P = 0.0066, partial η^2^ = 0.2426, 95% CI of partial η^2^ = [0.0310–0.3883], BF_10_ = 6.52), but not at V1 (F_1,27_ = 3.8517, P = 0.0601, partial η^2^ = 0.1248, 95% CI of partial η^2^ = [0.0001–0.3402], BF_10_ = 1.06) or V3A (F_1,27_ = 0.2216, P = 0.6416, partial η^2^ = 0.0081, 95% CI of partial η^2^ = [0.0000–0.0788], BF_10_ = 0.22). Box plots are overlaid on violin plots. Black dots indicate mean values across participants, and violin plots show kernel probability densities of individual participant data. VP: ventral posterior area. *P < 0.05, **P < 0.01, ****P < 0.001.

First, we performed a whole-brain analysis to identify brain regions that respond more strongly to the HS images than the KS images. Higher-visual areas beyond V1 showed significantly stronger BOLD signal amplitudes in response to the HS images than the KS images (Fig. 8b; see Fig. S12a for the regions that responded more strongly to the KS images). These regions are consistent with those reported to be involved in processing higher-order statistics derived from natural scene images in previous studies^32–34^.

Second, we tested whether the effects of attentional suppression on BOLD responses differ between task-irrelevant HS and KS images in higher-visual areas such as V2, VP (ventral posterior area), V3, V4v, and V3A, as well as V1, as the regions of interest (ROIs). These visual areas were retinotopically defined for each participant (see Methods for details). We used easy and harder RSVP tasks to compare the effects of attentional suppression on BOLD signals in response to the HS and KS images for each ROI. In particular, we calculated the fMRI suppression index separately for each ROI and for each of the HS and KS image conditions (Fig. 8c). This index was defined as the percentage reduction in BOLD signal amplitudes under the harder RSVP task compared to the easy RSVP task (see Methods for detailed calculation). A smaller value of the index indicates a lower effect of attentional suppression on BOLD signals. As shown in Fig. 8c (see also Fig. S13a for BOLD signal amplitudes prior to the calculation of suppression indices), the areas showing reduced effects of suppression on task-irrelevant HS images compared to KS images match the areas in monkeys that respond to the higher-order statistics derived from NS images^32–34^. A two-way repeated-measures ANOVA on fMRI suppression indices with factors of ROI (V1, V2, VP, V3, V4v, vs V3A) and Image (HS vs KS) revealed significant main effects of ROI (F_2.29,61.95_ = 6.5843, P = 0.0017, partial η^2^ = 0.1961, 95% CI of partial η^2^ = [0.0940–0.3243], BF_10_ = 821.36), Image (F_1,27_ = 7.0106, P = 0.0134, partial η^2^ = 0.2061, 95% CI of partial η^2^ = [0.0039–0.4220], BF_10_ > 10^4^), as well as a significant interaction between them (F_2.43,65.67_ = 3.2439, P = 0.0363, partial η^2^ = 0.1073, 95% CI of partial η^2^ = [0.0370–0.2135], BF_10_ = 0.13). Subsequent analyses revealed a significant simple effects of Image at V2 (F_1,27_ = 8.5033, P = 0.0071, partial η^2^ = 0.2395, 95% CI of partial η^2^ = [0.0224–0.4695], BF_10_ = 6.17), VP (F_1,27_ = 6.4939, P = 0.0168, partial η^2^ = 0.1939, 95% CI of partial η^2^ = [0.0090–0.4086], BF_10_ = 2.98), V3 (F_1,27_ = 20.5149, P = 0.0001, partial η^2^ = 0.4318, 95% CI of partial η^2^ = [0.1696–0.6453], BF_10_ = 245.80), and V4v (F_1,27_ = 8.6490, P = 0.0066, partial η^2^ = 0.2426, 95% CI of partial η^2^ = [0.0310–0.3883], BF_10_ = 6.52), but not at V1 (F_1,27_ = 3.8517, P = 0.0601, partial η^2^ = 0.1248, 95% CI of partial η^2^ = [0.0001–0.3402], BF_10_ = 1.06) or V3A (F_1,27_ = 0.2216, P = 0.6416, partial η^2^ = 0.0081, 95% CI of partial η^2^ = [0.0000–0.0788], BF_10_ = 0.22). Notably, the reduced effect of attentional suppression on the HS images cannot be attributed to differences in RSVP task performance, as no significant difference in RSVP performance was observed between the HS and KS image conditions (Fig. S13b).

Third, we asked whether effects of attentional suppression are reflected on the pattern of BOLD signals in voxels of each visual area, not just the overall amplitude. We computed correlations of BOLD signal patterns between the easy and harder RSVP tasks using representation similarity analysis (RSA^68^), performed for each image type (HS, KS) within each ROI (V1, V2, VP, V3, V4v, V3A). If the reduced effect of attentional suppression on task-irrelevant HS images relative to KS images is reflected in the higher visual areas, the difference in correlation coefficients between the HS and KS images should be larger in these areas. The results of the RSA showed that the correlation coefficients were significantly larger for HS than KS images across ROIs (Fig. S14a). We also found significantly or markedly larger differences in the correlation coefficients in higher-visual areas than for V1 (Fig. S14b). These results further support the conclusion that higher-visual areas beyond V1 are primarily involved in the reduced effect of attentional suppression on the task-irrelevant higher-order statistics compared to lower-order statistics (Fig. 7b).

Fourth, we examined whether attention-related regions in the parietal and frontal cortices play a crucial role. The analyses above suggested the reduced effect of attentional suppression on task-irrelevant higher-order statistics in higher-visual areas. Another possibility is that the degrees of attentional suppression from the source regions in the parietal and frontal cortices differ depending on the image type presented as task-irrelevant (HS or KS images). To test this possibility, a whole-brain analysis was performed to identify regions associated with enhanced attentional suppression. Fig. S12b shows stronger BOLD signal amplitudes during the harder RSVP task than the easy RSVP task in the bilateral intraparietal sulci (IPS) and frontal eye field (FEF), which are regarded as source regions for attentional suppression^69–73^ (see Fig. S12c for regions activated more strongly during the easy RSVP task). Importantly, these regions did not show differential BOLD signal amplitudes depending on the image type presented as task-irrelevant (HS or KS images), suggesting that the degree of attention was not different for HS and KS images.

Fifth, we tested whether functional connectivity between each of the four source regions associated with attentional suppression and the six visual areas differed depending on the task and image type, using a generalized psychophysiological interaction technique^74^ (see Methods for details). We applied a four-way ANOVA on functional connectivity with the within-participant factors of Source region (left IPS, right IPS, left FEF, vs right FEF), Visual area (V1, V2, VP, V3, V4v, vs V3A), Task (easy vs harder RSVP), and Image type (HS vs KS). We found a significant effect of Source region on functional connectivity, but not significant effects of the other factors or their interactions (Fig. S15; see Table S5 for detailed results of the ANOVA).

The analyses of BOLD signals and functional connectivity collectively suggest that the source regions of attentional suppression, such as the IPS and FEF, modulate the suppression of task-irrelevant images. However, this modulation does not depend on the type of task-irrelevant image. Instead, the reduced effects of attentional suppression on task-irrelevant higher-order statistics signals appear to arise within higher visual areas, rather than through top-down modulation from attentional control regions.

Why does the reduced effect of attentional suppression occur for a task-irrelevant feature generated from higher-order statistics compared to one derived from lower-order statistics? One possible explanation is that signals based on higher-order statistical information from the HS images are processed outside the temporally optimal window for attentional suppression. In a behavioral experiment (Experiment 12), we addressed this question. We psychophysically measured the time to detect the dominant orientations in the HS images and KS images (Fig. 9a; see Experiment 12 in Methods). This measurement employed a deadline paradigm, in which participants were asked to report the perceived orientation by a specific point in time, termed the “deadline,” on each trial^75–77^. We measured orientation discrimination performance at various deadlines and calculated the time required for performance to reach a certain level, termed “processing time,” for both HS and KS images (see Methods for detailed calculation).

**Fig 9:**
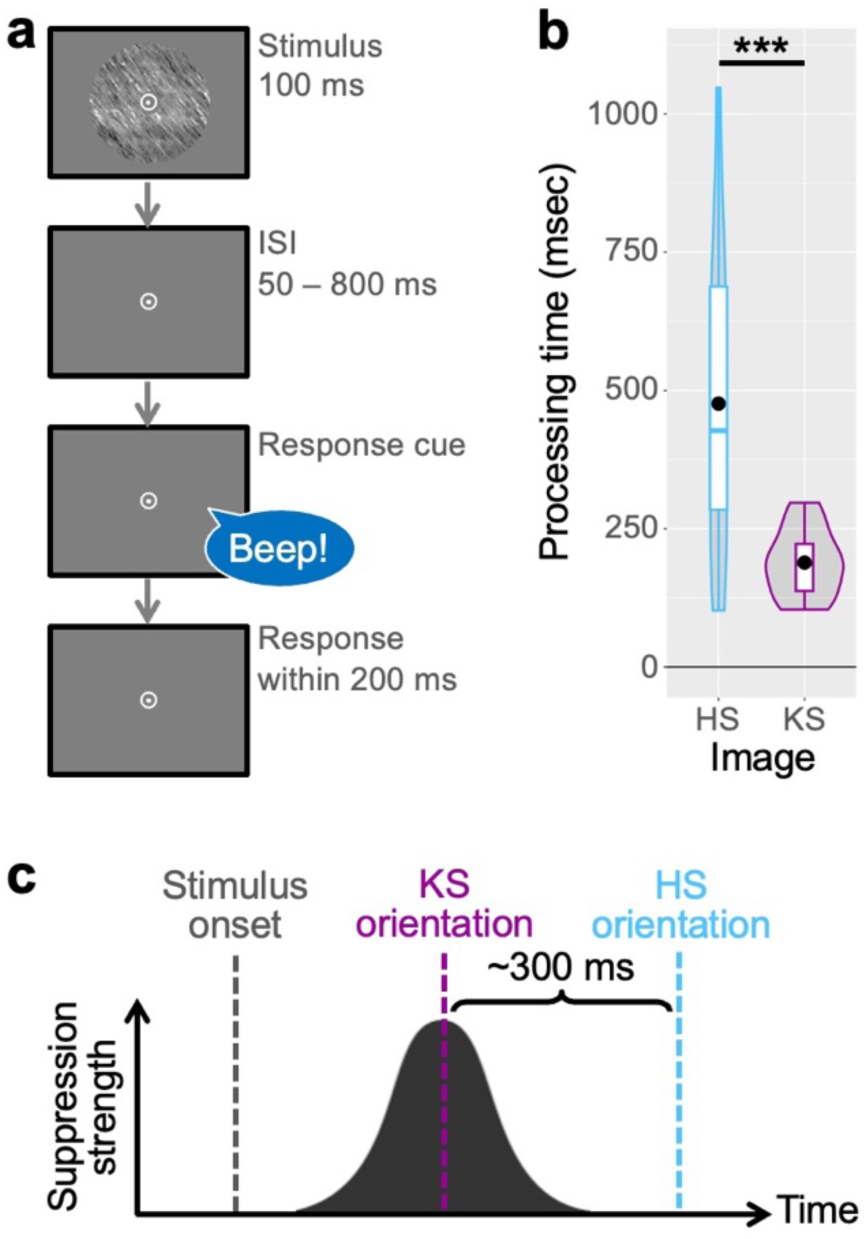
Procedures and results of Experiment 12 **a**, Illustrative timeline of a single trial in the orientation discrimination task, incorporating a response deadline. Participants were asked to report whether the dominant orientation of a presented image was tilted clockwise or counterclockwise relative to the vertical axis within 200 ms after an auditory response cue. **b**, Processing time computed from accuracies of the orientation discrimination task at different inter-stimulus intervals (ISIs; see Methods for details). The mean processing time for HS images was significantly longer—by 287.3 ms—compared to KS images (paired *t*-test; t_11_ = 3.6352, P = 0.0039, Cohen’s d = 1.0494, 95% CI of the difference in processing time = [113.3561–461.2772] ms, BF_10_ = 12.76). Box plots are overlaid on violin plots. Black dots indicate mean processing times across participants, and violin plots show kernel probability densities of individual processing times. ***P < 0.005. **c**, Graphical explanation of how suppressive signals interact with orientation information derived from KS images (“KS orientation”, magenta) and HS images (“HS orientation”, cyan). The results indicate that it took approximately 300 ms longer to detect the orientation in HS images compared to KS images.

The mean processing time for the HS images was significantly longer—by 287.3 ms—compared to the KS images (Fig. 9b; paired *t*-test; t_11_ = 3.6352, P = 0.0039, Cohen’s d = 1.0494, 95% CI of the difference in processing time = [113.3561–461.2772] ms, BF_10_ = 12.76). Thus, we estimate that it took approximately 300 ms longer for orientations in HS images to be detected compared to for those in KS images. Given that attentional suppression for artificial stimuli is unlikely to persist beyond 150 ms^78–83^, our result supports the hypothesis that dominant orientation representation from higher-order statistics emerges after the time window during which attentional suppression is most effective (Fig. 9c). However, it is important to note that this processing time is an estimate, and direct measurements of brain activity in future studies will be necessary to obtain a more precise assessment.

Based on the above results, we suggest that visible task-irrelevant features can still drive learning if they contain higher-order statistics. However, NS, HS, and PS images may have captured attention more strongly than KS, FS, or GP images, even when they were ”task-irrelevant.” Such enhanced salience could reduce attentional suppression rather than evaded it via statistical structure per se. If this bottom-up capture hypothesis is correct, NS, HS, and PS images could elicit greater eye-movement amplitudes than KS, FS, and GP images during the RSVP task.

To test this possibility, we conducted Experiment 13 with 12 new participants (see Experiment 13 in Methods for details). We recorded participants’ gaze positions and pupil diameters while they performed the RSVP task (Fig. 1b) under six conditions of task-irrelevant background images (NS, HS, PS, KS, FS, and GP). No significant differences were observed in eye movements or RSVP performance across the image conditions, as detailed below. First, from the time series of horizontal (Fig. S16a) and vertical (Fig. S16b) gaze positions, we calculated the radius encompassing the gaze-position distribution with a 95% confidence range (Fig. S16c). A one-way repeated-measures ANOVA on this radius revealed no significant effect of Image (NS, HS, PS, KS, FS, vs GP; F_4.02,44.21_ = 0.4153, P = 0.7976, partial η^2^ = 0.0364, 95% CI of partial η^2^ = [0.0037–0.0619], BF_10_ = 0.0346). Second, using the time series of pupil diameter (Fig. S16d), we computed mean changes in pupil diameter (Fig. S16e). A one-way ANOVA showed no significant differences in mean changes in pupil diameter across the image conditions (F_3.37,37.10_ = 1.2619, P = 0.3022, partial η^2^ = 0.1029, 95% CI of partial η^2^ = [0.0178–0.1921], BF_10_ = 0.1374). Finally, we compared RSVP performance across the image types (Fig. S16f). A previous study reported that stronger deterioration of RSVP performance occurs when task-irrelevant stimuli are less effectively suppressed^16^. However, a one-way ANOVA revealed no significant effect of Image (F_3.36,36.99_ = 1.4036, P = 0.2554, partial η^2^ = 0.1132, 95% CI of partial η^2^ = [0.0113–0.1985], BF_10_ = 0.1725). Taken together, these results are inconsistent with the bottom-up capture hypothesis.

## Discussion

Unsupervised learning is one of the most prevailing computational^84–89^ and biological principles of learning^1–10,13,14,18^. Nevertheless, studies on VPL have indicated that VPL does not occur as a result of exposure to visible task-irrelevant features, due to attentional suppression of those features^17,40,41,48^. This has raised the controversy over whether unsupervised learning occurs in the context of VPL. When visible images were presented as task-irrelevant during task performance, VPL occurred only for natural scene images—not for artificial ones (Figs. 1e and 2b). In contrast, we found that VPL occurred for *both* visible natural scene and artificial images as a result of free viewing (Fig. 3c). Further experiments demonstrated that VPL occurred when the exposed task-irrelevant images contained higher-order statistics derived from natural scenes (Figs. 5b and 6b), whereas VPL did not occur when the images lacked such statistics (Figs. 4b, S7b, and S8c). Results from subsequent psychophysical and fMRI experiments showed that task-irrelevant features containing higher-order statistics are less susceptible to attentional suppression than those containing only lower-order statistics (Figs. 7b and 8c). Such reduced susceptibility was particularly observed in visual areas beyond V1 (Figs. 8c and S14b).

Empirical^90^ and theoretical studies^8,10^ suggest that the form of VPL depends on contexts composed by tasks and stimuli during training. The results of some experiments in this study appear to indicate that it is true: Under a natural scene context, unsupervised VPL occurs from mere exposure^91^. In contrast, under an artificial stimulus context, unsupervised VPL is not observed, since visible, task-irrelevant features appear to be suppressed by the inhibitory attentional system involved in a main task^16^, leading no VPL. However, the results of our behavioral and fMRI experiments suggest that unsupervised learning underlies VPL of both natural scenes and artificial images, but attentional suppression effects are dependent on the exposed form of a stimulus.

It is notable that the FS and KS images also include some higher-order statistics. However, because the FS images disrupt the spatial phase relationships that normally organize structures in natural scenes, and the KS images adjust only pixel luminance distributions without preserving spatial arrangements, the higher-order statistics in these images are much less spatially organized than those in natural scene (NS) images. This weaker spatial structure in the FS and KS images may not be sufficient to produce a clear dominant orientation, unlike the richly organized structures of the NS images. In contrast, the higher-order statistics derived from the NS images have distinctive and meaningful structures of correlations between different orientation channels and different spatial frequency channels at various locations, enabling them to generate dominant orientation signals.

One possible explanation for the results of Experiments 12 and 13 is that signals based on higher-order statistics from the NS images are processed outside the temporally optimal window for attentional suppression, allowing them to escape such suppression (Fig. 9c). However, it should be noted that this account has not yet been substantiated by sufficient existing evidence and therefore remains speculative. Another possibility is that attentional suppression effects are stronger for lower-order statistics than for higher-order statistics. It has been shown that first-order motion, defined by changes in luminance, tends to be more readily suppressed by attention compared to second-order motion, which is defined by changes in texture or contrast^92^. This suggests that simpler, lower-order stimuli are more susceptible to attentional modulation than more complex, higher-order stimuli. However, this possibility does not appear to be plausible if we consider the possibility that the attentional suppression effect found in our RSVP paradigm may generalize to other types of attentional processing tested by different paradigms. This is because strong attentional modulation effects—such as attentional blindness^93,94^ and Eureka effects^95,96^—are known to strongly influence the processing of higher-order stimuli. In sum, additional systematic experiments will be necessary to clarify the mechanisms underlying the present findings.

Although our results indicate that VPL of supra-threshold artificial stimuli presented as task-irrelevant during unrelated task performance did not occur, there are two studies that reported VPL even under such conditions^19,20^. As mentioned above, in these studies, the task target and supra-threshold task-irrelevant artificial stimuli were significantly spatially distant. VPL occurred perhaps because the task-irrelevant images were located beyond the suppressive attentional skirts centered on the task target.

What are the implications of our findings for statistical learning research, in which unsupervised learning plays a central role? Statistical learning studies have typically employed passive viewing paradigms, allowing participants to freely observe presented stimuli without performing any concurrent task^5–8,10^. In contrast, our standard paradigm exposed participants to task-irrelevant images while they performed the RSVP task, which prevented VPL for task-irrelevant images containing only the lower-order statistics of the NS images due to attentional suppression. This raises the question of whether this paradigm disrupts typical statistical learning observed in classical paradigms^5–8,10^. Typical statistical learning may remain unaffected by the RSVP task. Since statistical learning is unlikely to rely on low-level learning processes^5,8,10^, it may escape attentional suppression.

The results of the present study may have important implications for the advancement of AI. Most AI models do not account for substantial differences in processing depending on stimulus type. Our findings indicate that whether the input consists of natural scenes or artificial images can determine different interactions between top-down attentional processes and stimulus processing in visual areas, leading to opposite outcomes: the presence or absence of learning. This underscores that the statistical structure of the stimulus—particularly the presence of higher-order statistics typical of natural scenes—can critically influence learning processes and outcomes. Incorporating sensitivity to such structure in AI models may enhance their ability to generalize, adapt, and learn from real-world data more effectively, especially under unsupervised or weakly supervised conditions.

In our study, we found that although free viewing of both supra-threshold artificial and natural scene images led to VPL of a feature of the images, exposure to these images as task-irrelevant during task performance resulted in VPL of the feature only from natural scene images. The results of further experiments, using behavioral and brain imaging measurements, suggest that higher-order statistics derived from natural scene images, when task-irrelevant, are less susceptible to attentional suppression than those lacking such statistics, perhaps because the slower processing speed of these statistics signals causes them to miss the optimal temporal window for attentional suppression exerted by attentional source areas. These results suggest that unsupervised learning is fundamental to VPL but can be hindered by factors such as attentional suppression.

## Methods

### Participants

A total of 250 naïve participants with normal or corrected-to-normal vision were enrolled in this study. The experimental protocols were approved by the Institutional Review Board. Prior to participation, all participants provided written informed consent. Data from 18 participants were subsequently excluded from the analyses (see Exclusion of participants for details). Consequently, the final analyses included data from 232 participants (18 to 41 years old; 147 males and 85 females).

In Experiments 1-7, and 9, we predetermined the sample size as 12 participants per group, aligning with prior studies that employed analogous designs and stimuli, as conducted by our group and other groups^18,19,97^. The sample size for the fMRI experiment (Experiment 11) was set to 28 participants, which was selected to be consistent with that used in previous fMRI studies that examined attentional modulation using ROI-based analyses^98–102^.

### Stimuli

Visual stimuli were presented in a grayscale within a visual annulus of 0.75° to 6.5° extending from the center of a uniform gray background (luminance = 0.50).

We employed oriented Gabor patch (GP) images (Fig. 1d; spatial frequency = 1 cycle/degree, contrast = 100%, Gaussian filter sigma = 3°, random spatial phase) to assess participants’ sensitivity to orientation. The chosen spatial frequency closely approximated the mean normalized spatial frequency observed in the natural scene images used in this study (∼1 cycle/degree).

Regarding the natural scene (NS) images (Fig. 1c, highlighted with orange rings), we curated two sets of 40 images from multiple databases^103–106^, with each set exhibiting a dominant orientation power peaking at 45° and 135°, referred to as 45°-dominant and 135°-dominant NS images, respectively (Fig. S2). Orientation power was computed using a Sobel filter^107^.

Fourier-scrambled (FS) images (Fig. 1c, highlighted with green rings) were synthesized to replicate the orientation and spatial frequency distributions of the NS images, as well as the features derived from luminance histograms of the NS images, such as the mean and variance. This was achieved by Fourier transforming the NS images, randomizing the phase spectra, and then applying an inverse Fourier transformation^108^.

Kurtosis- and skewness-matched (KS) images (Fig. 4a, highlighted with purple rings) were generated as random images that preserved the lower-order statistics (orientation and spatial frequency distributions) and marginal statistics (the mean, variance, kurtosis, and skewness of luminance histograms) of the NS images. These images were synthesized using a pyramid-based synthesis algorithm^109^ (# orientation = 4, # spatial scale = 5, window size = 15-pixel square).

The Portilla-Simoncelli (PS) images (Fig. 5a, highlighted with blue rings) were generated to obtain the higher-order statistics (the correlation among the outputs of filters tuned to different positions, orientations, and scales) of the NS images, in addition to the statistics that KS images have. The PS images were synthesized using the Portilla-Simoncelli algorithm^27^ (# orientation = 4, # spatial scale = 5, window size = 15-pixel square).

Synthesized images such as FS, KS, and PS images were commonly used in prior studies for the fields of computer vision and cognitive neuroscience^26,27,32,45,57,108^. The use of these images allowed researchers to isolate the contributions of lower-order, marginal, and higher-order statistics to perception and cognition. By employing these images in the current study, our findings can be compared with those in the prior studies.

The HS images (Fig. 6a, highlighted with cyan rings) were tailored to match the marginal and higher-order statistics of the NS images but differed in terms of the lower-order statistics (see Fig. S5 for graphical illustration). These images were created via the following steps. First, higher-order statistics of the NS images were computed. Second, pixels in the NS images were randomly shuffled in the pixel space. Third, lower-order statistics (orientation and spatial frequency distributions) and marginal statistics (the mean, variance, kurtosis, and skewness of luminance histograms) were computed using the shuffled images. Finally, each HS image was synthesized based on the computed lower-order, marginal, and higher-order statistics via the Portilla-Simoncelli algorithm. Although derived from the 45°-dominant and 135°-dominant NS images, the HS images did not display a prominent orientation power at 45° or 135° (Fig. S6).

### Experiment 1

The objective of Experiment 1 was to test whether visual perceptual learning (VPL) of orientation occurs as a result of exposures to supra-threshold NS and FS images as task-irrelevant. One group of 12 participants was exposed to the NS images (NS group), while the other group of 12 participants was exposed to the FS images (FS group). In each group, half of the participants were exposed to the 45°-dominant images, and the other half were exposed to the 135°-dominant images. Experiment 1 consisted of a 10-day exposure+task stage, with one-day pre- and post-test stages (Fig. 1a). The purpose of the exposure+task stage was to expose participants to the orientation features of the NS and FS images as task-irrelevant stimuli during performance of a given task (Fig. 1b). To ensure that these images remained task-irrelevant, we assigned a demanding task on the fovea to participants^13,17,40,110^. In the rapid serial visual presentation (RSVP) task, participants were required to identify two digits from a rapid sequence of ten characters presented in the center of a display (Fig. 1b). Each trial began with a 500-ms fixation period. Then, ten characters were presented within a 1°-diameter circle. Each character was displayed for 33 ms, followed by a 17-ms blank. Of these characters, two were digits (light gray, adjusted for each participant), and the remaining eight were alphabets (dark gray, luminance = 0.24). The digits and alphabets were randomly chosen from 1, 2, and 3 and A, C, D, E, G, H, J, K, M, N, P, R, S, U, V, W, and X for each trial, respectively. After the stimulus presentation, participants were required to make two successive responses by pressing buttons on a keyboard to indicate the first and second digits in the order they appeared (no time limit). In each trial, a task-irrelevant image was presented at the periphery. Forty images were presented in a random order for each block of 40 trials. Participants completed 15 blocks of the RSVP tasks per day, with brief breaks between blocks upon a participant’s request.

To ensure that the RSVP task remained sufficiently demanding during the 10-day exposure+task stage, we adjusted the difficulty of the RSVP task in the following two ways. First, after the pre-test stage and on the same day, the participants practiced the RSVP task, and difficulty adjustments were performed. This involved testing multiple luminance levels for the digits until accuracy reached approximately 70%, which was selected individually for each participant. Second, if a participant recorded an accuracy of over 90% on any given day of the exposure+task stage, the difficulty was recalibrated by modifying the luminance level, with the aim of achieving an accuracy of approximately 70% on the subsequent day.

The aim of the pre- and post-test stages was to measure changes in participants’ sensitivities to orientation before and after the exposure+task stage. Specifically, we adopted an orientation discrimination task (Fig. 1d) used in a previous study^19^, which aligns with the aim. Each trial was initiated with a 500-ms fixation period, followed by the presentation of two Gabor patches for 200 ms each. These presentations were separated by a 400-ms inter-stimulus interval. Participants were asked to report whether the orientations of the two Gabor patches were the same or different by pressing a button on a keyboard (no time limit). Each block consisted of 48 trials. In half of the trials per block, one Gabor patch was at the reference orientation (45 or 135 degrees), and the other patch was at an orientation differing by ±Δ degrees from the reference. In the other half, both Gabor patches were at the reference orientation. The Δ value was fixed within a block, and the numbers of “same” and “different” trials were equal, ensuring a balanced design. The Δ value for each block was 12, 8, 5, 3, or 1 degree, with a decrease every two blocks. The reference orientation for each block was either 45 or 135 degrees, and the orientation was alternated between blocks. Participants completed ten blocks, with brief breaks between blocks upon a participant’s request. Performance in each of the pre- and post-stages was defined as the mean accuracy averaged across the Δ values of 12, 8, 5, 3, and 1 degree. Both “same” and “different” trials were used for the calculation of accuracy. An improvement in performance (Fig. 1e) was defined by the subtraction of the performance in the pre-test stage from that in the post-test stage. Note that we employed this combination of task and procedure, rather than the classical pairing of a two-interval forced-choice task and a staircase procedure, because a well-known previous study on task-irrelevant VPL of orientation used the same task without a staircase procedure^19^, enabling a direct comparison of our results with those reported in that study.

### Experiment 2

The objective of Experiment 2 was to test whether VPL of spatial frequency occurs as a result of exposure to supra-threshold NS and FS images. The procedures were identical to those in Experiment 1, with the following two exceptions.

First, from the 80 NS images used in this study, we selected 20 images for the exposure+task stage based on their spatial frequency. Specifically, the 80 NS images were sorted according to their normalized spatial frequency, irrespective of their dominant orientations. The 10 images with the lowest values were defined as lower-spatial-frequency dominant images, and the 10 images with the highest values as higher-spatial frequency dominant images (see Fig. S4 for frequency spectra). The mean value of the normalized spatial frequency was 0.63 Hz for the lower-spatial-frequency dominant images and 1.48 Hz for the higher-spatial-frequency dominant images. In the NS group (N = 12), half of the participants were exposed to the lower-spatial-frequency dominant images, and the other half to the higher-spatial-frequency dominant images. Correspondingly, for the FS group (N = 12), we selected 10 lower-spatial-frequency and 10 higher-spatial-frequency dominant FS images. Each block of the exposure+task stage thus contained 10 images, each presented four times in random order.

Second, participants performed a spatial frequency discrimination task (Fig. 2a) during the pre- and post-test stages. The time-course of each trial is identical to that of the orientation discrimination task (Fig. 1d), except that band-pass filtered noise stimuli (see below for details) were presented, instead of Gabor patches. In half of the 48 trials per block, one stimulus was at the reference frequency (0.63 or 1.48 Hz), and the other at a frequency differing by +Δ Hz from the reference. In the remaining half, both stimuli were presented at the reference frequency. The Δ value was fixed within a block, and the numbers of “same” and “different” trials were equal, ensuring a balanced design. The Δ value for each block was 0.33, 0.23, 0.16, 0.11, or 0.07 Hz, decreasing every two blocks. The reference frequency for each block was either 0.63 or 1.48 Hz, alternating between blocks. As in the orientation discrimination task, performance was defined as the mean accuracy averaged across the Δ values.

We generated two band-pass filtered noise stimuli for each trial (Fig. 2a), as follows. First, two white noise images (100% contrast) were created. Second, each image was transformed into Fourier space, where a band-pass filter (Gaussian kernel; sigma = 0.05 × center frequency) was applied to the frequency components. The filtered images were then transformed back into pixel space using an inverse Fourier transformation. Finally, each filtered image was multiplied by a Gaussian envelope (sigma = 3°).

### Experiment 3

The objective of Experiment 3 was to test whether VPL of orientation occurs under a self-paced passive viewing protocol using supra-threshold NS and FS images. The procedures were identical to those in Experiment 1, except that the exposure+task stage in Experiment 1 was replaced with an exposure stage (Fig. 3a). During the exposure stage, participants were instructed to maintain central fixation and were allowed to view presented images at their own pace (Fig. 3b). One group of 12 participants viewed the NS images (NS group), while the other group of 12 participants viewed the FS images (FS group). In each trial, a white fixation point was presented at the center of the display for 500 ms, followed by the presentation of an image around the fixation point for another 500 ms. Then, the color of the fixation point turned black, indicating that participants were able to proceed to the next image by pressing a button on a keyboard.

### Experiments 4-7

The objective of Experiments 4-7 was to test whether VPL of orientation occurs as a result of exposures to the task-irrelevant KS, PS, HS, and GP images, respectively. The procedures were identical to those in Experiment 1, except that the KS (Experiment 4), PS (Experiment 5), HS (Experiment 6), or GP (Experiment 7) images were used in the exposure+task stage. A different group of 12 participants was used in each experiment. Half of the 12 participants were exposed to the images created from the 45°-dominant NS images, while the other half were exposed to the images created from the 135°-dominant NS images. See Stimuli in Methods for details of how GP, KS, PS, and HS images were created.

### Experiment 8

The objective of Experiment 8 was to examine whether the orientation information contained in the HS images was sub-threshold or supra-threshold. Specifically, 12 participants engaged in an orientation discrimination task using the HS images (Fig. 6a, highlighted with cyan rings).

In each trial, following a 500-ms fixation period, an HS image was presented for 500 ms. Participants were instructed to indicate whether the predominant orientation of the HS image was tilted in a clockwise or counterclockwise direction with respect to the vertical axis by pressing a button on a keyboard (no time limit). Each block consisted of 80 trials. In half of the trials, the HS images were created from the 45°-dominant NS images. In the other half, the HS images were created from the 135°-dominant NS images. Each participant completed five task blocks, with brief breaks between blocks upon a participant’s request.

### Experiment 9

The objective of Experiment 9 (N = 12) was to test whether VPL of orientation in the KS images occurs due to their reduced visibility (low-visibility KS images, Fig. S8a; see below for details). The procedures were identical to those in Experiment 4, except that the low-visibility KS images were used in the exposure+task stage.

To reduce the visibility of orientation in the KS images, we adopted a pixel randomization method in which a certain proportion of image pixels were randomly selected and shuffled (Fig. S8a). This method allowed us to systematically manipulate accuracy in the orientation discrimination task as a function of the proportion of preserved pixels, while maintaining the kurtosis and skewness of the KS images.

We conducted a pilot experiment to determine the proportion of preserved pixels that would yield mean accuracy in the orientation discrimination task equivalent to that obtained with the HS images (73.7%; Fig. S3b). Twelve new participants were asked to perform the orientation discrimination task (Fig. S3a) using the low-visibility KS images with five levels of pixel preservation (90, 70, 50, 30, and 10%). Each participant completed five task blocks, each consisting of 80 trials, with brief breaks between blocks upon a participant’s request. The proportion of preserved pixels decreased across blocks in the order of 90, 70, 50, 30, and 10%. The mean accuracy equivalent to that for the HS images was achieved when 25% of the pixels were preserved (Fig. S8b). Accordingly, Experiment 9 employed low-visibility KS images with 25% of the pixels preserved during the exposure+task stage.

### Experiment 10

The objective of Experiment 10 was to test whether effects of attentional suppression are weaker on images containing the higher-order statistics of the NS images (i.e., HS images) than the images lacking the higher-order statistics (i.e., KS images). Twelve participants took part in an experiment using a modified version of a distractor suppression paradigm (Fig. 7a), which is typically combined with visual search tasks^64,65^.

This paradigm involved the same stimulus presentation as that in the exposure+task stage of Experiment 1, where a sequence of characters was presented on the fovea while an image was presented at the periphery. Each block consisted of 40 trials. The exception in Experiment 10 was that half of the trials in each block were based on 45°-dominant images, and the other half were based on 135°-dominant images. Using this stimulus, each participant performed tasks under three conditions (Fig. 6a; focused attention, RSVP, and diverted attention) in a given order. In the focused attention condition, as in Experiment 8, participants were asked to indicate whether the predominant orientation of the presented image was clockwise or counterclockwise compared to vertical, ignoring the sequence of characters on the fovea. This condition was conducted over two blocks, with one block presenting the HS images and the other the KS images, in a random order for each participant. A brief break was provided between the blocks upon a participant’s request.

In the RSVP condition, the participants performed the same RSVP task as in Experiment 1 for two blocks, with a brief break between the blocks upon a participant’s request. The luminance of the digits in the RSVP task was individually set for each participant, as in Experiment 1. As task-irrelevant stimuli, the HS images were presented in one block and the KS images were presented in another, with the order randomized for each participant.

In the diverted attention condition, participants performed the orientation discrimination task for 20% (8 trials) of each block and the RSVP task for the remaining 80% (32 trials). At the beginning of each trial, participants were not informed of which task would be assigned. The participants were informed of the task for each trial only after the offset of the stimulus. In trials with the orientation discrimination task, a green fixation point was presented for one second following the offset of the visual stimulus. After the color of the fixation point returned to white, the participants were asked to report the predominant orientation of the image. The RSVP trials were identical to those in the original RSVP condition. Participants were instructed to prioritize the RSVP task as the primary task. A total of eight blocks were conducted in the diverted attention condition, with the HS and KS images alternately presented in each block. The order of the image presentation was randomized for each participant.

### Experiment 11

The objective of Experiment 11 was to identify the brain areas/networks that manifest differential effects of attentional suppression on brain responses to the higher- vs lower-order statistics of the NS images. In this experiment, 28 participants were enrolled.

We used an established paradigm^66,67^ to measure the effects of attentional suppression on BOLD signal responses to task-irrelevant stimuli (Fig. 8a). In this paradigm, while images were presented at the periphery, participants were instructed to count the number of target stimulus presentations on the fovea under two task conditions. Under one condition (easy RSVP task), the target was a fixation point that randomly turned from white to pink for 250 ms during a 10-sec stimulus period. The number of color changes during each stimulus period was randomly determined between three and six times. In the other condition (harder RSVP task), the target was the letter ’T’. During the stimulus period, a randomly rotated ’T’ or ’L’ was presented every 250 ms. The number of ’T’s presented during the stimulus period was randomly selected between eight and fourteen. Under both conditions, participants were asked to report whether the number of target appearances was odd or even by pressing a button during the fixation period following the stimulus period. Previous research has shown that the harder RSVP task results in greater attentional suppression in BOLD signal responses in the visual cortex than does the easy RSVP task^66,67^.

For the fMRI measurements, a block design was employed. During each session, participants were instructed to focus on a white fixation point at the center of a display. Each session consisted of a 6-sec fixation period followed by 16 repetitions of a set of a 10-sec stimulus and 10-sec fixation periods (326 sec in total). During the stimulus period, 40 task-irrelevant images were presented for 250 ms each in a random order. The stimulus periods were classified into eight types based on the image type (HS or KS), the dominant orientation of the images (45°-or 135°-dominant), and the task (easy or harder RSVP). These eight types were repeated twice in random order during the 16 stimulus periods. During the initial 4-sec interval of the fixation period, the fixation point was colored green, and participants were asked to press a button within this interval. The fixation point was turned white for the subsequent 5-sec interval, and in the last second, it changed to a ’C’ or ’T.’ A ’C’ indicated that the next stimulus period would be under the easy RSVP task, while a ’T’ indicated the harder RSVP task. Each participant underwent eight sessions, with brief breaks between the sessions upon a participant’s request.

After the main fMRI session above, retinotopic mapping^111–115^ was also conducted to delineate regions of interest (ROIs) within the visual cortex (V1, V2, VP, V3, V4v, and V3A). We used standard retinotopic stimuli and localizer stimuli corresponding to the visual cortical areas responsive to the images used in the study.

MRI data were acquired using a 3T MRI system (Siemens, Prisma) with a 64-channel head coil. Functional images were obtained using an echo-planar imaging (EPI) sequence with an echo time (TE) of 30 ms, a repetition time (TR) of 1,000 ms, and a flip angle (FA) of 64°. The field of view was set at 192×192 mm^2^, with a matrix configuration of 64×64, and a slice thickness of 3 mm. A total of 51 contiguous slices, aligned parallel to the AC-PC line, were obtained utilizing an acceleration factor of 2 via the GRAPPA parallel imaging technique^116^ and a multiband factor of 3. We confirmed that participants were awake by monitoring their eyes through a video camera throughout the sessions. Physiological monitoring was conducted to record participants’ respiratory and cardiac signals using a pressure sensor and a pulse oximeter, respectively. For the purpose of surface reconstruction, high-resolution anatomical images of the entire brain were obtained using a 3D T1-weighted MPRAGE sequence^117^ with a resolution of 0.7 × 0.7 × 0.7 mm^3^ (TR = 2,180 ms, TE = 2.95 ms, inversion time = 1,100 ms, FA = 8°).

In the first stage of fMRI analysis, functional images were preprocessed by the following steps. First, head motion correction was performed using the AFNI command 3dvolreg^118^ (http://afni.nimh.nih.gov/afni). Second, physiological noise was removed based on the participants’ respiratory and cardiac signals collected during the sessions^119^. Third, components related to slow drift and head motion were regressed out based on a general linear model (GLM) analysis using in-house software. Fourth, slice timing correction was performed with the first slice as a reference. Subsequent preprocessing steps were performed using SPM12 (https://www.fil.ion.ucl.ac.uk/spm/). The anatomical images were normalized to a standard space. The functional images were coregistered to the anatomical image for each participant and spatially smoothed with an isotropic Gaussian kernel of 6 mm (full-width at half-maximum).

In the second stage, we performed five analyses to achieve the objective of Experiment 11, as detailed below. First, we applied whole-brain GLM analysis to the spatially smoothed fMRI data obtained from the main experiment using SPM12 (https://www.fil.ion.ucl.ac.uk/spm/). For each participant, we modeled the four conditions for the two types of images (HS and KS) and tasks (easy and harder RSVP), as well as the fixation period and button response, as the regressors. Note that the components related to head motion had already been removed in the preprocessing stage. In each voxel, time series of BOLD signals was regressed against a time series of predicted fMRI responses created by convolving a canonical hemodynamic response function with the regressors. For the second-level analysis, we applied paired t-test to a mean contrast value between the conditions across participants for each voxel. The alpha-level threshold of 0.05 was adjusted by Bonferroni correction for multiple comparisons. No cluster-level assessment was employed.

Second, we identified the voxels that were responsive to the spatial extent stimulated by the HS and KS images in the visual areas (V1, V2, VP, V3, V4v, and V3A) and defined these areas as regions of interest (ROIs) for each participant. To achieve this, we applied GLM to fMRI data obtained based on the retinotopic and localizer stimuli and delineated the ROIs on flattened cortical maps using FreeSurfer (https://surfer.nmr.mgh.harvard.edu/). The ROIs were projected to a standard space for subsequent analyses.

Third, we applied ROI-based GLM analysis to the non-smoothed fMRI data obtained from the main experiment using SPM12. In each ROI, signal time series were extracted from voxels within the ROI and regressed against the time series of predicted fMRI responses for each ROI, as in the first analysis. The subtractions of the regression coefficient for the fixation period from those for the four conditions in the stimulus period were defined as the BOLD signal amplitudes for these conditions. Then, the BOLD signal amplitudes were averaged across the voxels in each ROI. We calculated the fMRI suppression index separately for each participant, ROI (V1, V2, VP, V3, V4v, and V3A), and image type (HS, KS). The index was defined as the percentage reduction of the BOLD signal amplitudes under the harder RSVP task compared to the easy RSVP task based on the following formula.

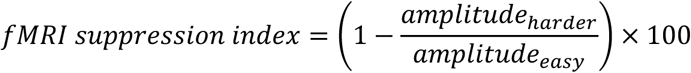

Here, amplitude*_easy_* and amplitude*_harder_* represent BOLD signal amplitudes under the easy and harder RSVP tasks, respectively.

Fourth, we performed representational similarity analysis (RSA^68^) based on the ROI-based GLM analysis (described above) using Decoding Toolbox^120^ (https://sites.google.com/site/tdtdecodingtoolbox/). In RSA, we calculated a correlation coefficient of BOLD signal amplitude patterns of voxels between the easy and harder RSVP tasks for each participant, ROI, and image type.

Fifth, we performed generalized psychophysiological interaction (gPPI^74^) analysis based on the ROI-based GLM analysis using CONN toolbox^121^ (https://web.conn-toolbox.org/home). The bilateral IPS and FEF were used as source regions. Functional connectivity was defined as a regression coefficient of one of the source regions for a target ROI.

### Experiment 12

The objective of Experiment 12 was to test whether the time required for orientation processing based on the higher-order statistics of the NS images (i.e., HS images) differed from that based on the lower-order statistics of the NS images (i.e., KS images). For this objective, 12 participants took part in an orientation discrimination task based on a deadline paradigm^75–77^.

In this paradigm, a ’deadline’ was defined as the allocated time interval from the stimulus offset to the execution of a button response. The measurement of accuracy for various deadlines, coupled with an analysis of the time required for accuracy to reach an asymptote, enabled the quantification of time required to process a certain stimulus.

In each trial (Fig. 9a), following a 1-sec fixation period, an image was presented for 100 ms. After a predetermined inter-stimulus interval (ISI), a 50-ms high-pitched auditory cue (640 Hz) was presented, requiring participants to report the predominant orientation of the image (clockwise or counterclockwise relative to vertical) within 200 ms. Trials in which the button response exceeded this deadline were excluded from subsequent analysis. Such deadline violations occurred in 4.47 ± 0.84% (mean ± standard error) of trials across participants. Each block consisted of 40 trials. Half of the trials in each block included 45°-dominant images, and the other half included 135°-dominant images. Each participant completed 24 task blocks, with brief breaks between the blocks upon a participant’s request. Two types of images (HS and KS images) and six ISIs (50, 100, 200, 300, 500, 800 ms) were paired and assigned to each block. Across the 24 blocks, 12 pairs of the image types and ISIs were presented twice in a random order.

In the analysis, curve fitting was performed based on accuracies at the six ISI levels using the following equation:

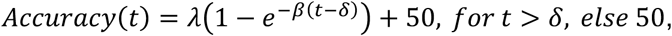

where 𝑡 represents an ISI. Curve fitting was performed for each image type and each participant. The processing time shown in Fig. 9b was defined as 𝛿 + 𝛽^-1.^, corresponding to the number of milliseconds required for the accuracy to reach the 67% asymptote.

### Experiment 13

The objective of Experiment 13 was to test the bottom-up capture hypothesis that NS, HS, and PS images should elicit greater eye-movement amplitudes than KS, FS, and GP images during the RSVP task. To this end, we recorded participants’ eye-movements while they performed the RSVP task (Fig. 1b) under six conditions of the background images (NS, HS, PS, KS, FS, and GP). Twelve new participants completed 12 task blocks, each consisting of 40 trials, with brief breaks between blocks upon a participant’s request. Across blocks, 12 combinations of the two dominant orientations (45 and 135 deg) and the six image conditions (NS, HS, PS, KS, FS, and GP) were presented in random order. As in Experiment 1, the difficulty of the RSVP task was individually adjusted to yield an accuracy of approximately 70% for each participant.

During each task block, participants’ gaze position and pupil diameter were continuously recorded from their left eye using EyeLink 1000 Plus system (SR Research, Canada) operated with EyeLink software (version 5.15) at a sampling rate of 1000 Hz. A standard five-point calibration procedure provided by the EyeLink system was performed at the start of each block and repeated until the average and maximum gaze position errors were less than 0.5 deg and 1.0 deg, respectively. Participants were instructed to refrain from blinking during the 500-ms stimulus period (Fig. 1b) and were allowed to blink only during the response period.

For each block, the entire time series of gaze position and pupil diameter were preprocessed according to the guidelines proposed by prior studies^122,123^. First, a low-pass filter (cutoff frequency = 4 Hz) was applied to the pupil-diameter time series. Second, time points from 50 ms before to 50 ms after blinks detected by the EyeLink system were marked as missing for both gaze-position and pupil-diameter data. Third, outlier detection was performed based on the first derivative of the pupil-diameter time series. Specifically, a time point 𝑡 was defined as an outlier if

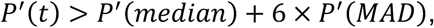

where 𝑃^/^(𝑡) denotes the first derivative at 𝑡, and 𝑃^/^(𝑚𝑒𝑑𝑖𝑎𝑛) and 𝑃^/^(𝑀𝐴𝐷) represent the median and median absolute deviation of the first derivative, respectively. For both gaze-position and pupil-diameter time series, time points within ±120 ms of an outlier were also marked as missing. Fourth, a piecewise cubic Hermite interpolation polynomial (PCHIP) was applied to each cluster of missing data shorter than 250 samples. The remaining missing data were excluded from the calculation of means and standard deviations in subsequent analyses. Fifth, the entire time series of gaze position and pupil diameter were segmented into trials spanning –200 to 700 ms relative to the onset of the 500-ms stimulus period. Finally, only trials that met the following criteria were included in subsequent analyses: (a) fewer than 30% missing data points during the baseline period (–200 to 0 ms relative to the stimulus onset), (b) fewer than 30% missing data points across the entire trial, (c) no blinks during the baseline period, and (d) gaze positions confined within 0.75 deg of the display center (corresponding to the fixation disk) during the baseline period. As a result, 94.9% of the trials were retained for further analyses.

Figs. S16a, b, and d show the mean time series of horizontal gaze position, vertical gaze position, and changes in pupil diameter, respectively. For each trial, gaze positions were baseline-normalized by subtracting the mean gaze position during the baseline period from the entire time series of that trial. This normalization was performed separately for the horizontal and vertical axes. Because pupil diameter varied substantially across participants, pupil-diameter data for each trial were converted to percent changes relative to the baseline period according to the following equation:

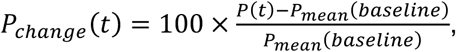

where 𝑃(𝑡) denotes the pupil diameter at each time point, 𝑃*_mean_*(𝑏𝑎𝑠𝑒𝑙𝑖𝑛𝑒) the mean pupil diameter during the baseline period, and 𝑃*_change_*(𝑡) the percent change in pupil diameter at time *t*.

The gaze-position and pupil-diameter data were analyzed and compared across the image conditions as follows. First, for each trial, we calculated the radius encompassing the gaze-position distribution during the 500-ms stimulus period with a 95% confidence range. To assess potential differences in eye movements, trial-averaged values of this radius were compared across the six image conditions (Fig. S16c). Second, to examine possible differences in pupil dilation among the six image types, we computed the mean change in pupil diameter during 200–500 ms after the stimulus onset, considering the typical latency of pupil responses (approximately 200 ms^123,124^). The trial-averaged changes in pupil diameter were then compared across the conditions (Fig. S16e).

We also identified long-distance saccades directed toward the background images during the 500-ms stimulus period (see below for details). Only four trials (0.007%) contained such saccades. Long-distance saccades were defined as follows. First, the EyeLink system detected all saccades regardless of their amplitude. Second, a detected saccade was classified as a long-distance saccade if it met the following criteria: (a) the gaze position was confined within an area corresponding to the fixation disk (0.75 deg from the display center) for at least 40 ms before the saccade onset, and (b) the gaze position remained outside the fixation area for at least 30 ms within the time window extending from the saccade onset to 40 ms after the saccade offset.

### Control Experiment

The objective of Control Experiment was to test the possibility that the results of Experiments 1, 3-5 could be explained by differences in the strength of orientation information contained in the images. To examine this possibility, 12 participants performed the same orientation discrimination task (Fig. S3a) as in Experiment 8 with the NS, FS, KS, and PS images. Each block consisted of 80 trials, and the same image type was used throughout each block. Each participant completed one task block for each image type. The order of the image types was randomized for each participant. Forty images were 45°-dominant, and the other 40 images were 135°-dominant. The order of the images was randomized for each block. Brief breaks were provided between blocks upon a participant’s request.

### Apparatus

All experiments were conducted in a dim room, with the experimental setup controlled via Psychtoolbox 3^125^ in MATLAB. In the behavioral experiments (Experiments 1-10, 12, 13, and Control Experiment), visual stimuli were presented on an LED monitor (1,920×1,080 resolution). For Experiments 1-10, 13, and Control Experiment, the monitors operated at a refresh rate of 60 Hz, whereas for Experiment 12, a higher refresh rate of 240 Hz was employed. In Experiment 11, visual stimuli were projected onto a frosted glass screen (1,920×1,080 resolution) positioned above participants’ heads with a refresh rate of 60 Hz.

### Exclusion of participants

In this study, data from 18 participants were excluded from the analyses for the following reasons. In Experiments 1, 4-6, four participants were unable to continue due to health-related concerns in the middle of the exposure+task stage. In Experiment 2, one participant reported after completing the experiment that the participant had attended to and attempted to memorize the task-irrelevant images. Another participant showed unexpectedly high accuracy (> 90%) in the RSVP task for 6 out of 10 days during the exposure+task stage. These two participants were therefore excluded. One participant in Experiment 8 who had previously participated in a different experiment on the same day was excluded to mitigate potential biases introduced by fatigue. In Experiments 8, 10, 12, and Control Experiment, data from five participants were excluded since the experiments were terminated due to program errors. In Experiment 9, data from one participant was excluded due to nearly chance-level performance in the RSVP task during the exposure+task stage. In Experiment 13, data from four participants were excluded due to unstable recording of gaze positions and pupil diameters. One participant exhibited near-chance-level performance in the RSVP task under the experimental setting with the eye tracker.

### Curve fitting

We calculated the perceptual threshold and slope of the accuracy data for each participant, separately for the pre- and post-test stages and for the exposed and unexposed features. Thresholds and slopes were obtained by fitting the accuracy data with a Weibull function using psignfit toolbox^126^. The threshold was defined as the Δ value (see Experiments 1 and 2 for details) corresponding to 80% task accuracy, which was pre-determined following a previous study^127^.

One exceptionally high threshold value (> mean + 6 SD) for the unexposed orientation in the post-test stage of one participant in the FS group of Experiment 1 was excluded from subsequent analyses.

### Statistics

All the statistical tests performed in this study were two-tailed. The significance threshold was set to an alpha level of 0.05. *t*-tests and ANOVA were performed utilizing the function anovakun in the R statistical software package. When a three-way ANOVA showed a significant three-way interaction, we conducted a two-way ANOVA per each level of a factor^128^ to identify simple interaction effects via subsequent analysis. Because the initial three-way ANOVA indicated a three-way interaction, this approach protected against inflating the Type I error rate in the subsequent analyses. This is a statistically well-established and valid way to avoid type I and type II errors^129–132^. Effect size for *t*-tests was quantified using Cohen’s d, while partial η^2^ served as the measure of effect size for ANOVAs. The computation of the 95% confidence intervals was based on a bias-corrected and accelerated (BCa) method over 2,000 bootstrap repetitions. For ANOVAs, we applied Mendoza’s multi-sample sphericity test to evaluate the sphericity assumption. When necessary, we adjusted the degrees of freedom using Greenhouse-Geisser’s ε to correct any violations.

We computed Bayes factors in R using the BayesFactor (https://richarddmorey.github.io/BayesFactor/) and bayesTestR packages^133^. We employed the Jeffreys-Zellner-Siow prior with a “wide” scale (r = 2/2) for fixed effects and a “nuisance” scale (r = 1) for random effects (i.e., participant-specific intercepts). These parameters were selected because they share properties with common frequentist tests^134^. We tested interaction effects across matched models that included all underlying main effects^135^.

## Acknowledgements

We thank M. Kaminokado, K. Haruhana, K. Uchida, and M. Leung for their assistance with behavioral data collection and K. Ueno, C. Suzuki, and K. Kunieda from the Support Unit for Functional Magnetic Resonance Imaging at RIKEN Center for Brain Science for their assistance with MRI measurements. This work was supported by NSF-BSF 2241417, R01EY027841, R01EY019466 (to T.W.), R01 EY031705 (to Y.S.), JSPS Kakenhi Grants 19H01041 and 20H05715, and JST Moonshot R&D Grant JPMJMS2013 (to K.S.).

## Notes

### Competing Interest Statement

The authors have declared no competing interest.

### Summary of Updates

Five additional experiments and related analyses were added; All sections were revised accordingly; Author list updated; Supplementary files updated.

