## Supplementary Tables and Figures for "Unsupervised visual learning is revealed for task-irrelevant natural scenes due to reduced attentional suppression effects in visual areas"

1 Department of Cognitive and Psychological Sciences, Brown University
2 Center for Brain Science, RIKEN
3 Department of Literature, Nagoya University
4 Equally contributed
5 Corresponding Author

### Supplementary Figures and Tables

**Table S1: Results of ANOVA on performance of the pre- and post-test stages in Experiments 1-7, 9**

| **Experiment 1** | | | | | | | |
|---|---|---|---|---|---|---|---|
| **Three-way ANOVA for NS and FS images** | | **F value** | **P value** | **Partial $\eta^2$** | **95% CI of partial $\eta^2$** | | **$BF_{10}$** |
| Main effects | Group | $F_{1,22}$ = 0.0459 | 0.8323 | 0.0021 | [0.0000–0.0200] | | 2.00 |
| | Test | $F_{1,22}$ = 2.2891 | 0.1445 | 0.0942 | [0.0001–0.4076] | | 3.53 |
| | Orientation | $F_{1,22}$ = 7.7903 | 0.0106 | 0.2615 | [0.0138–0.5074] | * | 1.90 |
| Two-way interactions | Group × Test | $F_{1,22}$ = 5.9240 | 0.0235 | 0.2121 | [0.0061–0.4397] | * | 7.58 |
| | Group × Orientation | $F_{1,22}$ = 1.6381 | 0.2139 | 0.0693 | [0.0000–0.3623] | | 0.85 |
| | Test × Orientation | $F_{1,22}$ = 10.5287 | 0.0037 | 0.3237 | [0.0560–0.5515] | *** | 3.35 |
| Three-way interaction | Group × Test × Orientation | $F_{1,22}$ = 5.8216 | 0.0246 | 0.2092 | [0.0036–0.4781] | * | 2.52 |
| **Two-way ANOVA for NS image** | | **F value** | **P value** | **Partial $\eta^2$** | **95% CI of partial $\eta^2$** | | **$BF_{10}$** |
| Main effects | Test | $F_{1,11}$ = 9.2612 | 0.0112 | 0.4571 | [0.0412–0.7403] | * | 291.85 |
| | Orientation | $F_{1,11}$ = 14.1095 | 0.0032 | 0.5619 | [0.0835–0.8458] | *** | 25.42 |
| Two-way interaction | Test × Orientation | $F_{1,11}$ = 19.3135 | 0.0011 | 0.6371 | [0.3418–0.8191] | *** | 49.76 |
| **Two-way ANOVA for FS image** | | **F value** | **P value** | **Partial $\eta^2$** | **95% CI of partial $\eta^2$** | | **$BF_{10}$** |
| Main effects | Test | $F_{1,11}$ = 0.3659 | 0.5575 | 0.0322 | [0.0000–0.2383] | | 0.20 |
| | Orientation | $F_{1,11}$ = 0.8083 | 0.3879 | 0.0685 | [0.0000–0.4071] | | 0.20 |
| Two-way interaction | Test × Orientation | $F_{1,11}$ = 0.2955 | 0.5976 | 0.0262 | [0.0000–0.2207] | | 0.06 |
| **Experiment 2** | | | | | | | |
| **Three-way ANOVA for NS and FS images** | | **F value** | **P value** | **Partial $\eta^2$** | **95% CI of partial $\eta^2$** | | **$BF_{10}$** |
| Main effects | Group | $F_{1,22}$ = 0.8180 | 0.3756 | 0.0358 | [0.0000–0.2366] | | 3.05 |
| | Test | $F_{1,22}$ = 17.038 | 0.0004 | 0.4364 | [0.1032–0.6829] | *** | 137.27 |
| | Frequency | $F_{1,22}$ = 4.8781 | 0.0379 | 0.1815 | [0.0010–0.4885] | * | 2.07 |
| Two-way interactions | Group × Test | $F_{1,22}$ = 10.6166 | 0.0036 | 0.3255 | [0.0641–0.5106] | *** | 10.63 |
| | Group × Frequency | $F_{1,22}$ = 0.1382 | 0.7136 | 0.0062 | [0.0000–0.0685] | | 0.56 |
| | Test × Frequency | $F_{1,22}$ = 3.1151 | 0.0914 | 0.1240 | [0.0000–0.4543] | | 1.03 |
| Three-way interaction | Group × Test × Frequency | $F_{1,22}$ = 7.2760 | 0.0132 | 0.2485 | [0.0029–0.4819] | * | 1.43 |
| **Two-way ANOVA for NS image** | | **F value** | **P value** | **Partial $\eta^2$** | **95% CI of partial $\eta^2$** | | **$BF_{10}$** |
| Main effects | Test | $F_{1,11}$ = 37.2494 | 0.0001 | 0.7720 | [0.5789–0.8790] | *** | > $10^4$ |
| | Frequency | $F_{1,11}$ = 4.2980 | 0.0624 | 0.2810 | [0.0004–0.6012] | | 12.92 |

| | | | | | | | |
|---|---|---|---|---|---|---|---|
| Two-way interaction | Test × Frequency | $F_{1,11}$ = 25.6514 | 0.0004 | 0.6999 | [0.4495–0.8250] | *** | 21.83 |
| **Two-way ANOVA for FS image** | | **F value** | **P value** | **Partial $\eta^2$** | **95% CI of partial $\eta^2$** | | **BF$_{10}$** |
| Main effects | Test | $F_{1,11}$ = 0.2977 | 0.5962 | 0.0264 | [0.0000–0.2475] | | 0.18 |
| | Frequency | $F_{1,11}$ = 1.3768 | 0.2654 | 0.1112 | [0.0000–0.5819] | | 0.35 |
| Two-way interaction | Test × Frequency | $F_{1,11}$ = 0.2697 | 0.6138 | 0.0239 | [0.0000–0.1508] | | 0.08 |
| **Experiment 3** | | | | | | | |
| **Three-way ANOVA for NS and FS images** | | **F value** | **P value** | **Partial $\eta^2$** | **95% CI of partial $\eta^2$** | | **BF$_{10}$** |
| Main effects | Group | $F_{1,22}$ = 0.2260 | 0.6392 | 0.0102 | [0.0000–0.1222] | | 0.66 |
| | Test | $F_{1,22}$ = 27.6724 | $< 10^{-4}$ | 0.5571 | [0.2318–0.7195] | *** | $> 10^{4}$ |
| | Orientation | $F_{1,22}$ = 0.0840 | 0.7747 | 0.0038 | [0.0000–0.0422] | | 8.13 |
| Two-way interactions | Group × Test | $F_{1,22}$ = 4.1505 | 0.0538 | 0.1587 | [0.0030–0.4200] | | 1.58 |
| | Group × Orientation | $F_{1,22}$ = 0.6583 | 0.4258 | 0.0291 | [0.0000–0.2724] | | 0.44 |
| | Test × Orientation | $F_{1,22}$ = 25.1603 | 0.0001 | 0.5335 | [0.2530–0.7127] | *** | 41.17 |
| Three-way interaction | Group × Test × Orientation | $F_{1,22}$ = 0.0017 | 0.9673 | 0.0001 | [0.0000–0.0001] | | 0.49 |
| **Experiment 4** | | | | | | | |
| **Two-way ANOVA for KS image** | | **F value** | **P value** | **Partial $\eta^2$** | **95% CI of partial $\eta^2$** | | **BF$_{10}$** |
| Main effects | Test | $F_{1,11}$ = 0.6535 | 0.4360 | 0.0561 | [0.0000–0.4466] | | 0.25 |
| | Orientation | $F_{1,11}$ = 0.0234 | 0.8812 | 0.0021 | [0.0000–0.0159] | | 0.16 |
| Two-way interaction | Test × Orientation | $F_{1,11}$ = 0.0092 | 0.9254 | 0.0008 | [0.0000–0.0052] | | 0.05 |
| **Experiment 5** | | | | | | | |
| **Two-way ANOVA for PS image** | | **F value** | **P value** | **Partial $\eta^2$** | **95% CI of partial $\eta^2$** | | **BF$_{10}$** |
| Main effects | Test | $F_{1,11}$ = 7.0718 | 0.0222 | 0.3913 | [0.0298–0.7107] | * | 56.28 |
| | Orientation | $F_{1,11}$ = 1.7385 | 0.2141 | 0.1365 | [0.0005–0.5035] | | 0.37 |
| Two-way interaction | Test × Orientation | $F_{1,11}$ = 8.6700 | 0.0133 | 0.4408 | [0.2115–0.6047] | * | 0.89 |
| **Experiment 6** | | | | | | | |
| **Two-way ANOVA for HS image** | | **F value** | **P value** | **Partial $\eta^2$** | **95% CI of partial $\eta^2$** | | **BF$_{10}$** |
| Main effects | Test | $F_{1,11}$ = 10.8027 | 0.0072 | 0.4955 | [0.1096–0.7385] | ** | 23.63 |
| | Orientation | $F_{1,11}$ = 0.2066 | 0.6583 | 0.0184 | [0.0000–0.1153] | | 0.42 |
| Two-way interaction | Test × Orientation | $F_{1,11}$ = 16.1700 | 0.0020 | 0.5951 | [0.3204–0.7699] | *** | 1.24 |
| **Experiment 7** | | | | | | | |
| **Two-way ANOVA for GP image** | | **F value** | **P value** | **Partial $\eta^2$** | **95% CI of partial $\eta^2$** | | **BF$_{10}$** |
| Main effects | Test | $F_{1,11}$ = 0.0293 | 0.8671 | 0.0027 | [0.0000–0.0221] | | 0.16 |
| | Orientation | $F_{1,11}$ = 0.8781 | 0.3689 | 0.0739 | [0.0000–0.4179] | | 0.23 |
| Two-way interaction | Test × Orientation | $F_{1,11}$ = 1.1918 | 0.2983 | 0.0978 | [0.0001–0.4709] | | 0.07 |

| Experiment 9 | | | | | | | |
|---|---|---|---|---|---|---|---|
| **Two-way ANOVA for lower-visibility KS image** | | **F value** | **P value** | **Partial $\eta^2$** | **95% CI of partial $\eta^2$** | | **$BF_{10}$** |
| Main effects | Test | $F_{1,11} = 1.1175$ | 0.3131 | 0.0922 | [0.0000–0.4081] | | 0.38 |
| | Orientation | $F_{1,11} = 0.0828$ | 0.7789 | 0.0075 | [0.0000–0.0722] | | 0.17 |
| Two-way interaction | Test × Orientation | $F_{1,11} = 1.8130$ | 0.2052 | 0.1415 | [0.0002–0.5720] | | 0.10 |
| *P < 0.05, **P < 0.01, *** P < 0.005 | | | | | | | |

#### Table S2: Results of ANOVA on threshold of the pre- and post-test stages in Experiments 1-7, 9

| Experiment 1 | | | | | | | |
|---|---|---|---|---|---|---|---|
| **Three-way ANOVA for NS and FS images** | | **F value** | **P value** | **Partial $\eta^2$** | **95% CI of partial $\eta^2$** | | **$BF_{10}$** |
| Main effects | Group | $F_{1,22} = 0.3058$ | 0.5861 | 0.0144 | [0.0000–0.1523] | | 0.45 |
| | Test | $F_{1,21} = 4.9368$ | 0.0374 | 0.1903 | [0.0068–0.3927] | * | 1.19 |
| | Orientation | $F_{1,21} = 4.4305$ | 0.0475 | 0.1742 | [0.0030–0.4588] | * | 0.26 |
| Two-way interactions | Group × Test | $F_{1,21} = 2.2926$ | 0.1449 | 0.0984 | [0.0001–0.3151] | | 1.20 |
| | Group × Orientation | $F_{1,21} = 4.5660$ | 0.0445 | 0.1786 | [0.0029–0.4514] | * | 0.40 |
| | Test × Orientation | $F_{1,21} = 6.9904$ | 0.0152 | 0.2497 | [0.0025–0.5202] | * | 0.39 |
| Three-way interaction | Group × Test × Orientation | $F_{1,21} = 4.8006$ | 0.0399 | 0.1861 | [0.0007–0.4981] | * | 0.74 |
| **Two-way ANOVA for NS image** | | **F value** | **P value** | **Partial $\eta^2$** | **95% CI of partial $\eta^2$** | | **$BF_{10}$** |
| Main effects | Test | $F_{1,11} = 5.2753$ | 0.0423 | 0.3241 | [0.0313–0.5761] | * | 12.81 |
| | Orientation | $F_{1,11} = 7.9455$ | 0.0167 | 0.4194 | [0.0856–0.7005] | * | 2.46 |
| Two-way interaction | Test × Orientation | $F_{1,11} = 13.1042$ | 0.0040 | 0.5436 | [0.0292–0.8333] | *** | 4.56 |
| **Two-way ANOVA for FS image** | | **F value** | **P value** | **Partial $\eta^2$** | **95% CI of partial $\eta^2$** | | **$BF_{10}$** |
| Main effects | Test | $F_{1,10} = 0.4148$ | 0.5340 | 0.0398 | [0.0000–0.2872] | | 0.15 |
| | Orientation | $F_{1,10} = 0.0006$ | 0.9807 | 0.0001 | [0.0000–0.0000] | | 0.16 |
| Two-way interaction | Test × Orientation | $F_{1,10} = 0.0915$ | 0.7685 | 0.0091 | [0.0000–0.0957] | | 0.04 |

| Experiment 2 | | | | | | | |
|---|---|---|---|---|---|---|---|
| **Three-way ANOVA for NS and FS images** | | **F value** | **P value** | **Partial $\eta^2$** | **95% CI of partial $\eta^2$** | | **$BF_{10}$** |
| Main effects | Group | $F_{1,22} = 0.9761$ | 0.3339 | 0.0425 | [0.0000–0.2414] | | 0.70 |
| | Test | $F_{1,22} = 23.4840$ | 0.0001 | 0.5163 | [0.2304–0.7099] | *** | 40.13 |
| | Frequency | $F_{1,22} = 6.1970$ | 0.0208 | 0.2198 | [0.0038–0.4698] | * | 11.12 |
| Two-way interactions | Group × Test | $F_{1,22} = 7.6801$ | 0.0111 | 0.2588 | [0.0135–0.4974] | * | 1.48 |
| | Group × Frequency | $F_{1,22} = 0.1287$ | 0.7232 | 0.0058 | [0.0000–0.0580] | | 0.34 |

| | | | | | | | |
|---|---|---|---|---|---|---|---|
| | Test × Frequency | $F_{1,22}$ = 2.2912 | 0.1443 | 0.0943 | [0.0000–0.3465] | | 0.77 |
| Three-way interaction | Group × Test × Frequency | $F_{1,22}$ = 5.2111 | 0.0325 | 0.1915 | [0.0000–0.4848] | * | 0.31 |
| **Two-way ANOVA for NS image** | | **F value** | **P value** | **Partial $\eta^2$** | **95% CI of partial $\eta^2$** | | **$BF_{10}$** |
| Main effects | Test | $F_{1,11}$ = 32.9931 | 0.0001 | 0.7500 | [0.4869–0.8970] | *** | 3759.7 |
| | Frequency | $F_{1,11}$ = 4.6403 | 0.0543 | 0.2967 | [0.0024–0.6374] | | 9.25 |
| Two-way interaction | Test × Frequency | $F_{1,11}$ = 19.8750 | 0.0010 | 0.6437 | [0.3657–0.7922] | *** | 11.12 |
| **Two-way ANOVA for FS image** | | **F value** | **P value** | **Partial $\eta^2$** | **95% CI of partial $\eta^2$** | | **$BF_{10}$** |
| Main effects | Test | $F_{1,11}$ = 1.9205 | 0.1933 | 0.1486 | [0.0002–0.5790] | | 0.25 |
| | Frequency | $F_{1,11}$ = 2.6846 | 0.1296 | 0.1962 | [0.0011–0.5894] | | 1.42 |
| Two-way interaction | Test × Frequency | $F_{1,11}$ = 0.1806 | 0.6790 | 0.0162 | [0.0000–0.1626] | | 0.19 |
| **Experiment 3** | | | | | | | |
| **Three-way ANOVA for NS and FS images** | | **F value** | **P value** | **Partial $\eta^2$** | **95% CI of partial $\eta^2$** | | **$BF_{10}$** |
| Main effects | Group | $F_{1,22}$ = 0.3179 | 0.5786 | 0.0142 | [0.0000–0.1060] | | 0.72 |
| | Test | $F_{1,22}$ = 17.6510 | 0.0004 | 0.4452 | [0.1457–0.6550] | *** | 1411.1 |
| | Orientation | $F_{1,22}$ = 0.0099 | 0.9216 | 0.0005 | [0.0000–0.0018] | | 0.55 |
| Two-way interactions | Group × Test | $F_{1,22}$ = 3.9599 | 0.0592 | 0.1525 | [0.0022–0.3662] | | 1.9 |
| | Group × Orientation | $F_{1,22}$ = 0.6543 | 0.4273 | 0.0289 | [0.0000–0.3187] | | 0.28 |
| | Test × Orientation | $F_{1,22}$ = 17.5269 | 0.0004 | 0.4434 | [0.1087–0.6827] | *** | 2.47 |
| Three-way interaction | Group × Test × Orientation | $F_{1,22}$ = 0.0637 | 0.8031 | 0.0029 | [0.0000–0.0338] | | 0.32 |
| **Experiment 4** | | | | | | | |
| **Two-way ANOVA for KS image** | | **F value** | **P value** | **Partial $\eta^2$** | **95% CI of partial $\eta^2$** | | **$BF_{10}$** |
| Main effects | Test | $F_{1,11}$ = 0.6466 | 0.4384 | 0.0555 | [0.0000–0.3705] | | 0.25 |
| | Orientation | $F_{1,11}$ = 0.6012 | 0.4545 | 0.0518 | [0.0000–0.3502] | | 0.19 |
| Two-way interaction | Test × Orientation | $F_{1,11}$ = 0.1522 | 0.7011 | 0.0139 | [0.0000–0.1290] | | 0.07 |
| **Experiment 5** | | | | | | | |
| **Two-way ANOVA for PS image** | | **F value** | **P value** | **Partial $\eta^2$** | **95% CI of partial $\eta^2$** | | **$BF_{10}$** |
| Main effects | Test | $F_{1,11}$ = 4.9716 | 0.0476 | 0.3113 | [0.0024–0.6222] | * | 9.75 |
| | Orientation | $F_{1,11}$ = 0.6375 | 0.4415 | 0.0548 | [0.0001–0.2793] | | 0.25 |
| Two-way interaction | Test × Orientation | $F_{1,11}$ = 3.5886 | 0.0848 | 0.2460 | [0.0128–0.4323] | | 0.45 |
| **Experiment 6** | | | | | | | |
| **Two-way ANOVA for HS image** | | **F value** | **P value** | **Partial $\eta^2$** | **95% CI of partial $\eta^2$** | | **$BF_{10}$** |
| Main effects | Test | $F_{1,11}$ = 6.1759 | 0.0303 | 0.3596 | [0.0155–0.6529] | * | 3.66 |
| | Orientation | $F_{1,11}$ = 0.0930 | 0.7661 | 0.0084 | [0.0000–0.0792] | | 0.25 |
| Two-way interaction | Test × Orientation | $F_{1,11}$ = 5.1656 | 0.0441 | 0.3195 | [0.0443–0.5667] | * | 0.47 |

| Experiment 7 | | | | | | | |
|---|---|---|---|---|---|---|---|
| **Two-way ANOVA for GP image** | | **F value** | **P value** | **Partial η²** | **95% CI of partial η²** | | **$BF_{10}$** |
| Main effects | Test | $F_{1,11}$ = 0.0479 | 0.8307 | 0.0043 | [0.0000–0.0434] | | 0.16 |
| | Orientation | $F_{1,11}$ = 0.1859 | 0.6747 | 0.0166 | [0.0000–0.1418] | | 0.18 |
| Two-way interaction | Test × Orientation | $F_{1,11}$ = 4.9866 | 0.0473 | 0.3119 | [0.0025–0.6450] | * | 0.07 |
| **Experiment 9** | | | | | | | |
| **Two-way ANOVA for lower-visibility KS image** | | **F value** | **P value** | **Partial η²** | **95% CI of partial η²** | | **$BF_{10}$** |
| Main effects | Test | $F_{1,11}$ = 0.8863 | 0.3667 | 0.0746 | [0.0000–0.4000] | | 0.33 |
| | Orientation | $F_{1,11}$ = 0.1091 | 0.7473 | 0.0098 | [0.0000–0.0955] | | 0.16 |
| Two-way interaction | Test × Orientation | $F_{1,11}$ = 1.0150 | 0.3353 | 0.0845 | [0.0000–0.4513] | | 0.08 |

*P < 0.05, **P < 0.01, *** P < 0.005

#### Table S3: Results of ANOVA on slope of the pre- and post-test stages in Experiments 1-7, 9

| Experiment 1 | | | | | | |
|---|---|---|---|---|---|---|
| **Three-way ANOVA for NS and FS images** | | **F value** | **P value** | **Partial η²** | **95% CI of partial η²** | **$BF_{10}$** |
| Main effects | Group | $F_{1,22}$ = 0.0380 | 0.8472 | 0.0018 | [0.0000–0.0148] | 0.13 |
| | Test | $F_{1,22}$ = 3.1960 | 0.0883 | 0.1321 | [0.0002–0.3479] | 0.27 |
| | Orientation | $F_{1,22}$ = 0.0863 | 0.7718 | 0.0041 | [0.0000–0.0357] | 0.08 |
| Two-way interactions | Group × Test | $F_{1,22}$ = 1.9148 | 0.1810 | 0.0836 | [0.0001–0.3168] | 0.12 |
| | Group × Orientation | $F_{1,22}$ = 0.1790 | 0.6766 | 0.0085 | [0.0000–0.0760] | 0.03 |
| | Test × Orientation | $F_{1,22}$ = 2.3185 | 0.1428 | 0.0994 | [0.0005–0.2796] | 0.12 |
| Three-way interaction | Group × Test × Orientation | $F_{1,22}$ = 1.8847 | 0.1843 | 0.0824 | [0.0000–0.2669] | 0.03 |
| **Experiment 2** | | | | | | |
| **Three-way ANOVA for NS and FS images** | | **F value** | **P value** | **Partial η²** | **95% CI of partial η²** | **$BF_{10}$** |
| Main effects | Group | $F_{1,22}$ = 0.1531 | 0.6993 | 0.0069 | [0.0000–0.0706] | 0.13 |
| | Test | $F_{1,22}$ = 3.1012 | 0.0921 | 0.1235 | [0.0002–0.3907] | 0.26 |
| | Frequency | $F_{1,22}$ = 2.1633 | 0.1555 | 0.0895 | [0.0000–0.3875] | 0.31 |
| Two-way interactions | Group × Test | $F_{1,22}$ = 0.0000 | 0.9972 | 0.0000 | [0.0000–0.0000] | 0.05 |
| | Group × Frequency | $F_{1,22}$ = 0.0298 | 0.8644 | 0.0014 | [0.0000–0.0108] | 0.06 |
| | Test × Frequency | $F_{1,22}$ = 0.6397 | 0.4324 | 0.0283 | [0.0000–0.1991] | 0.10 |
| Three-way interaction | Group × Test × Frequency | $F_{1,22}$ = 0.0939 | 0.7621 | 0.0043 | [0.0000–0.0473] | 0.002 |
| **Experiment 3** | | | | | | |
| **Three-way ANOVA for NS and FS images** | | **F value** | **P value** | **Partial η²** | **95% CI of partial η²** | **$BF_{10}$** |

| | | F value | P value | Partial $\eta^2$ | 95% CI of partial $\eta^2$ | | $BF_{10}$ |
|---|---|---|---|---|---|---|---|
| Main effects | Group | $F_{1,22} = 0.0011$ | 0.9739 | 0.0000 | [0.0000–0.0000] | | 0.17 |
| | Test | $F_{1,22} = 10.1870$ | 0.0042 | 0.3165 | [0.0192–0.5990] | *** | 2.05 |
| | Orientation | $F_{1,22} = 1.8396$ | 0.1888 | 0.772 | [0.0000–0.3563] | | 0.17 |
| Two-way interactions | Group × Test | $F_{1,22} = 1.2316$ | 0.2791 | 0.0530 | [0.0000–0.3087] | | 0.16 |
| | Group × Orientation | $F_{1,22} = 1.0565$ | 0.3152 | 0.0458 | [0.0000–0.3024] | | 0.06 |
| | Test × Orientation | $F_{1,22} = 1.5627$ | 0.2244 | 0.0663 | [0.0000–0.3402] | | 0.28 |
| Three-way interaction | Group × Test × Orientation | $F_{1,22} = 0.0699$ | 0.7939 | 0.0032 | [0.0000–0.0278] | | 0.01 |
| **Experiment 4** | | | | | | | |
| **Two-way ANOVA for KS image** | | **F value** | **P value** | **Partial $\eta^2$** | **95% CI of partial $\eta^2$** | | **$BF_{10}$** |
| Main effects | Test | $F_{1,11} = 0.3488$ | 0.5668 | 0.0307 | [0.0000–0.1415] | | 0.23 |
| | Orientation | $F_{1,11} = 2.6620$ | 0.1310 | 0.1948 | [0.0005–0.6146] | | 0.28 |
| Two-way interaction | Test × Orientation | $F_{1,11} = 0.8886$ | 0.3661 | 0.0747 | [0.0001–0.4270] | | 0.09 |
| **Experiment 5** | | | | | | | |
| **Two-way ANOVA for PS image** | | **F value** | **P value** | **Partial $\eta^2$** | **95% CI of partial $\eta^2$** | | **$BF_{10}$** |
| Main effects | Test | $F_{1,11} = 1.2241$ | 0.2922 | 0.1001 | [0.0000–0.4873] | | 0.35 |
| | Orientation | $F_{1,11} = 1.5190$ | 0.2435 | 0.1213 | [0.0004–0.4510] | | 0.31 |
| Two-way interaction | Test × Orientation | $F_{1,11} = 1.0726$ | 0.3226 | 0.0888 | [0.0000–0.5811] | | 0.14 |
| **Experiment 6** | | | | | | | |
| **Two-way ANOVA for HS image** | | **F value** | **P value** | **Partial $\eta^2$** | **95% CI of partial $\eta^2$** | | **$BF_{10}$** |
| Main effects | Test | $F_{1,11} = 2.1468$ | 0.1709 | 0.1633 | [0.0001–0.3637] | | 0.50 |
| | Orientation | $F_{1,11} = 0.6716$ | 0.4299 | 0.0575 | [0.0001–0.3504] | | 0.22 |
| Two-way interaction | Test × Orientation | $F_{1,11} = 1.6826$ | 0.2211 | 0.1327 | [0.0003–0.3599] | | 0.20 |
| **Experiment 7** | | | | | | | |
| **Two-way ANOVA for GP image** | | **F value** | **P value** | **Partial $\eta^2$** | **95% CI of partial $\eta^2$** | | **$BF_{10}$** |
| Main effects | Test | $F_{1,11} = 1.4495$ | 0.2539 | 0.1164 | [0.0001–0.4873] | | 0.38 |
| | Orientation | $F_{1,11} = 0.5796$ | 0.4625 | 0.0501 | [0.0000–0.3780] | | 0.20 |
| Two-way interaction | Test × Orientation | $F_{1,11} = 1.1444$ | 0.3076 | 0.0942 | [0.0001–0.3897] | | 0.13 |
| **Experiment 9** | | | | | | | |
| **Two-way ANOVA for lower-visibility KS image** | | **F value** | **P value** | **Partial $\eta^2$** | **95% CI of partial $\eta^2$** | | **$BF_{10}$** |
| Main effects | Test | $F_{1,11} = 0.5573$ | 0.4710 | 0.0482 | [0.0000–0.4561] | | 0.20 |
| | Orientation | $F_{1,11} = 0.6428$ | 0.4397 | 0.0552 | [0.0000–0.3162] | | 0.21 |
| Two-way interaction | Test × Orientation | $F_{1,11} = 0.1153$ | 0.7405 | 0.0104 | [0.0000–0.0939] | | 0.06 |

*P < 0.05, **P < 0.01, *** P < 0.005

#### Table S4: Results of ANOVA on d' of the pre- and post-test stages in Experiments 1-7, 9

| | | | | | | | |
|---|---|---|---|---|---|---|---|
| **Experiment 1** | | | | | | | |
| **Three-way ANOVA for NS and FS images** | | **F value** | **P value** | **Partial $\eta^2$** | **95% CI of partial $\eta^2$** | | **$BF_{10}$** |
| Main effects | Group | $F_{1,22} = 0.0016$ | 0.9683 | 0.0001 | [0.0000–0.0002] | | 3.45 |
| | Test | $F_{1,22} = 1.6104$ | 0.2177 | 0.0682 | [0.0001–0.3593] | | 5.80 |
| | Orientation | $F_{1,22} = 6.0508$ | 0.0222 | 0.2157 | [0.0073–0.4915] | * | 3.04 |
| Two-way interactions | Group × Test | $F_{1,22} = 6.5395$ | 0.0180 | 0.2291 | [0.0065–0.4693] | * | 13.20 |
| | Group × Orientation | $F_{1,22} = 3.9644$ | 0.0590 | 0.1527 | [0.0002–0.4395] | | 1.68 |
| | Test × Orientation | $F_{1,22} = 13.4139$ | 0.0014 | 0.3788 | [0.1069–0.5545] | *** | 9.32 |
| Three-way interaction | Group × Test × Orientation | $F_{1,22} = 4.4971$ | 0.0455 | 0.1697 | [0.0006–0.4507] | * | 4.82 |
| **Two-way ANOVA for NS image** | | **F value** | **P value** | **Partial $\eta^2$** | **95% CI of partial $\eta^2$** | | **$BF_{10}$** |
| Main effects | Test | $F_{1,11} = 8.0677$ | 0.0161 | 0.4231 | [0.0076–0.7292] | * | 257.26 |
| | Orientation | $F_{1,11} = 13.6385$ | 0.0035 | 0.5535 | [0.0579–0.8395] | *** | 43.35 |
| Two-way interaction | Test × Orientation | $F_{1,11} = 21.8768$ | 0.0007 | 0.6654 | [0.3552–0.8585] | *** | 88.60 |
| **Two-way ANOVA for FS image** | | **F value** | **P value** | **Partial $\eta^2$** | **95% CI of partial $\eta^2$** | | **$BF_{10}$** |
| Main effects | Test | $F_{1,11} = 0.7594$ | 0.4021 | 0.0646 | [0.0000–0.3604] | | 0.27 |
| | Orientation | $F_{1,11} = 0.0863$ | 0.7747 | 0.0078 | [0.0000–0.0722] | | 0.16 |
| Two-way interaction | Test × Orientation | $F_{1,11} = 0.9620$ | 0.3478 | 0.0804 | [0.0000–0.3834] | | 0.08 |
| **Experiment 2** | | | | | | | |
| **Three-way ANOVA for NS and FS images** | | **F value** | **P value** | **Partial $\eta^2$** | **95% CI of partial $\eta^2$** | | **$BF_{10}$** |
| Main effects | Group | $F_{1,22} = 1.0565$ | 0.3152 | 0.0458 | [0.0000–0.2887] | | 1.40 |
| | Test | $F_{1,22} = 25.4838$ | $< 10^{-4}$ | 0.5367 | [0.2492–0.7034] | *** | 340.16 |
| | Frequency | $F_{1,22} = 4.1986$ | 0.0526 | 0.1603 | [0.0013–0.4646] | | 1.50 |
| Two-way interactions | Group × Test | $F_{1,22} = 9.2660$ | 0.0060 | 0.2964 | [0.0628–0.4985] | ** | 4.15 |
| | Group × Frequency | $F_{1,22} = 0.0607$ | 0.8076 | 0.0028 | [0.0000–0.0263] | | 0.36 |
| | Test × Frequency | $F_{1,22} = 1.0273$ | 0.3218 | 0.0446 | [0.0000–0.3437] | | 0.53 |
| Three-way interaction | Group × Test × Frequency | $F_{1,22} = 5.7173$ | 0.0258 | 0.2063 | [0.0002–0.4293] | * | 0.50 |
| **Two-way ANOVA for NS image** | | **F value** | **P value** | **Partial $\eta^2$** | **95% CI of partial $\eta^2$** | | **$BF_{10}$** |
| Main effects | Test | $F_{1,11} = 42.7962$ | $< 10^{-4}$ | 0.7955 | [0.6623–0.8885] | *** | $> 10^4$ |
| | Frequency | $F_{1,11} = 3.9674$ | 0.0718 | 0.2651 | [0.0008–0.6484] | | 5.34 |
| Two-way interaction | Test × Frequency | $F_{1,11} = 16.8037$ | 0.0018 | 0.6044 | [0.1185–0.8190] | *** | 7.13 |
| **Two-way ANOVA for FS image** | | **F value** | **P value** | **Partial $\eta^2$** | **95% CI of partial $\eta^2$** | | **$BF_{10}$** |
| Main effects | Test | $F_{1,11} = 1.6262$ | 0.2285 | 0.1288 | [0.0000–0.5344] | | 0.26 |

| | | | | | | | |
|---|---|---|---|---|---|---|---|
| | Frequency | $F_{1,11}$ = 1.2162 | 0.2937 | 0.0996 | [0.0000–0.5335] | | 0.36 |
| Two-way interaction | Test × Frequency | $F_{1,11}$ = 0.5732 | 0.4649 | 0.0495 | [0.0000–0.2521] | | 0.12 |
| **Experiment 3** | | | | | | | |
| **Three-way ANOVA for NS and FS images** | | **F value** | **P value** | **Partial $\eta^2$** | **95% CI of partial $\eta^2$** | | **$BF_{10}$** |
| Main effects | Group | $F_{1,22}$ = 0.2251 | 0.6398 | 0.0101 | [0.0000–0.1081] | | 0.33 |
| | Test | $F_{1,22}$ = 21.3555 | 0.0001 | 0.4926 | [0.1677–0.6856] | *** | 1697.8 |
| | Orientation | $F_{1,22}$ = 0.0032 | 0.9553 | 0.0001 | [0.0000–0.0003] | | 2.83 |
| Two-way interactions | Group × Test | $F_{1,22}$ = 1.5948 | 0.2199 | 0.0676 | [0.0001–0.2859] | | 0.41 |
| | Group × Orientation | $F_{1,22}$ = 0.9238 | 0.3469 | 0.0403 | [0.0000–0.2984] | | 0.30 |
| | Test × Orientation | $F_{1,22}$ = 19.8375 | 0.0002 | 0.4742 | [0.1173–0.7011] | *** | 14.29 |
| Three-way interaction | Group × Test × Orientation | $F_{1,22}$ = 0.0137 | 0.9081 | 0.0006 | [0.0000–0.0037] | | 0.19 |
| **Experiment 4** | | | | | | | |
| **Two-way ANOVA for KS image** | | **F value** | **P value** | **Partial $\eta^2$** | **95% CI of partial $\eta^2$** | | **$BF_{10}$** |
| Main effects | Test | $F_{1,11}$ = 0.2023 | 0.6616 | 0.0181 | [0.0000–0.1992] | | 0.18 |
| | Orientation | $F_{1,11}$ = 0.0190 | 0.8928 | 0.0017 | [0.0000–0.0118] | | 0.15 |
| Two-way interaction | Test × Orientation | $F_{1,11}$ = 0.0005 | 0.9818 | 0.0000 | [0.0000–0.0000] | | 0.04 |
| **Experiment 5** | | | | | | | |
| **Two-way ANOVA for PS image** | | **F value** | **P value** | **Partial $\eta^2$** | **95% CI of partial $\eta^2$** | | **$BF_{10}$** |
| Main effects | Test | $F_{1,11}$ = 5.5973 | 0.0374 | 0.3372 | [0.0015–0.6856] | * | 14.09 |
| | Orientation | $F_{1,11}$ = 0.6075 | 0.4522 | 0.0523 | [0.0000–0.3218] | | 0.33 |
| Two-way interaction | Test × Orientation | $F_{1,11}$ = 7.6880 | 0.0181 | 0.4114 | [0.0196–0.1005] | * | 0.79 |
| **Experiment 6** | | | | | | | |
| **Two-way ANOVA for HS image** | | **F value** | **P value** | **Partial $\eta^2$** | **95% CI of partial $\eta^2$** | | **$BF_{10}$** |
| Main effects | Test | $F_{1,11}$ = 10.6427 | 0.0076 | 0.4917 | [0.0709–0.7758] | ** | 23.81 |
| | Orientation | $F_{1,11}$ = 0.7964 | 0.3913 | 0.0675 | [0.0000–0.4207] | | 0.46 |
| Two-way interaction | Test × Orientation | $F_{1,11}$ = 6.6033 | 0.0261 | 0.3751 | [0.0279–0.6807] | * | 1.14 |
| **Experiment 7** | | | | | | | |
| **Two-way ANOVA for GP image** | | **F value** | **P value** | **Partial $\eta^2$** | **95% CI of partial $\eta^2$** | | **$BF_{10}$** |
| Main effects | Test | $F_{1,11}$ = 0.0326 | 0.8600 | 0.0030 | [0.0000–0.0306] | | 0.16 |
| | Orientation | $F_{1,11}$ = 1.2493 | 0.2875 | 0.1020 | [0.0001–0.4695] | | 0.24 |
| Two-way interaction | Test × Orientation | $F_{1,11}$ = 0.9255 | 0.3567 | 0.0776 | [0.0000–0.4424] | | 0.07 |
| **Experiment 9** | | | | | | | |
| **Two-way ANOVA for lower-visibility KS image** | | **F value** | **P value** | **Partial $\eta^2$** | **95% CI of partial $\eta^2$** | | **$BF_{10}$** |
| Main effects | Test | $F_{1,11}$ = 2.7508 | 0.1254 | 0.2000 | [0.0001–0.5126] | | 1.42 |

| | | | | | | | |
|---|---|---|---|---|---|---|---|
| | Orientation | $F_{1,11}$ = 0.4085 | 0.5458 | 0.0358 | [0.0000–0.2980] | | 0.20 |
| Two-way interaction | Test × Orientation | $F_{1,11}$ = 1.3562 | 0.2688 | 0.1098 | [0.0001–0.4262] | | 0.19 |
| *P < 0.05, **P < 0.01, *** P < 0.005 | | | | | | | |

#### Table S5: Results of ANOVA on bias of the pre- and post-test stages in Experiments 1-7, 9

| **Experiment 1** | | | | | | | |
|---|---|---|---|---|---|---|---|
| **Three-way ANOVA for NS and FS images** | | **F value** | **P value** | **Partial $\eta^2$** | **95% CI of partial $\eta^2$** | | **$BF_{10}$** |
| Main effects | Group | $F_{1,22}$ = 0.5330 | 0.4731 | 0.0237 | [0.0000–0.1461] | | 0.16 |
| | Test | $F_{1,22}$ = 2.3391 | 0.1404 | 0.0961 | [0.0001–0.2927] | | 0.51 |
| | Orientation | $F_{1,22}$ = 0.3027 | 0.5877 | 0.0136 | [0.0000–0.0952] | | 0.07 |
| Two-way interactions | Group × Test | $F_{1,22}$ = 0.1203 | 0.7320 | 0.0054 | [0.0000–0.0562] | | 0.08 |
| | Group × Orientation | $F_{1,22}$ = 0.0532 | 0.8196 | 0.0024 | [0.0000–0.0196] | | 0.02 |
| | Test × Orientation | $F_{1,22}$ = 0.0006 | 0.9799 | 0.0000 | [0.0000–0.0000] | | 0.04 |
| Three-way interaction | Group × Test × Orientation | $F_{1,22}$ = 0.0442 | 0.8354 | 0.0020 | [0.0000–0.0172] | | 0.001 |
| **Experiment 2** | | | | | | | |
| **Three-way ANOVA for NS and FS images** | | **F value** | **P value** | **Partial $\eta^2$** | **95% CI of partial $\eta^2$** | | **$BF_{10}$** |
| Main effects | Group | $F_{1,22}$ = 0.0729 | 0.7897 | 0.0033 | [0.0000–0.0284] | | 0.11 |
| | Test | $F_{1,22}$ = 0.1670 | 0.6868 | 0.0075 | [0.0000–0.0446] | | 0.09 |
| | Frequency | $F_{1,22}$ = 0.1736 | 0.6809 | 0.0078 | [0.0000–0.0578] | | 0.07 |
| Two-way interactions | Group × Test | $F_{1,22}$ = 2.2284 | 0.1497 | 0.0920 | [0.0011–0.2210] | | 0.10 |
| | Group × Frequency | $F_{1,22}$ = 1.7310 | 0.2018 | 0.0729 | [0.0001–0.2085] | | 0.03 |
| | Test × Frequency | $F_{1,22}$ = 0.0322 | 0.8591 | 0.0015 | [0.0000–0.0235] | | 0.01 |
| Three-way interaction | Group × Test × Frequency | $F_{1,22}$ = 0.4600 | 0.5047 | 0.0205 | [0.0000–0.0901] | | 0.003 |
| **Experiment 3** | | | | | | | |
| **Three-way ANOVA for NS and FS images** | | **F value** | **P value** | **Partial $\eta^2$** | **95% CI of partial $\eta^2$** | | **$BF_{10}$** |
| Main effects | Group | $F_{1,22}$ = 0.3111 | 0.5826 | 0.0139 | [0.0000–0.1226] | | 0.86 |
| | Test | $F_{1,22}$ = 2.3851 | 0.1368 | 0.0978 | [0.0000–0.3770] | | 0.86 |
| | Orientation | $F_{1,22}$ = 0.6617 | 0.4247 | 0.0292 | [0.0000–0.2238] | | 0.42 |
| Two-way interactions | Group × Test | $F_{1,22}$ = 3.4048 | 0.0785 | 0.1340 | [0.0003–0.3861] | | 1.55 |
| | Group × Orientation | $F_{1,22}$ = 13.7908 | 0.0012 | 0.3853 | [0.0602–0.5927] | *** | 1.56 |
| | Test × Orientation | $F_{1,22}$ = 4.3829 | 0.0480 | 0.1661 | [0.0033–0.3275] | * | 0.66 |
| Three-way interaction | Group × Test × Orientation | $F_{1,22}$ = 0.0242 | 0.8778 | 0.0011 | [0.0000–0.0077] | | 0.63 |
| **Experiment 4** | | | | | | | |

| Two-way ANOVA for KS image | | F value | P value | Partial η$^2$ | 95% CI of partial η$^2$ | $BF_{10}$ |
|---|---|---|---|---|---|---|
| Main effects | Test | $F_{1,11} = 0.4037$ | 0.5382 | 0.0354 | [0.0000–0.3215] | 0.18 |
| | Orientation | $F_{1,11} = 0.0026$ | 0.9599 | 0.0002 | [0.0000–0.0008] | 0.15 |
| Two-way interaction | Test × Orientation | $F_{1,11} = 0.5782$ | 0.4630 | 0.0499 | [0.0000–0.3806] | 0.05 |
| **Experiment 5** | | | | | | |
| **Two-way ANOVA for PS image** | | **F value** | **P value** | **Partial η$^2$** | **95% CI of partial η$^2$** | **$BF_{10}$** |
| Main effects | Test | $F_{1,11} = 1.1858$ | 0.2995 | 0.0973 | [0.0000–0.3771] | 0.42 |
| | Orientation | $F_{1,11} = 1.5730$ | 0.2358 | 0.1251 | [0.0001–0.3845] | 0.22 |
| Two-way interaction | Test × Orientation | $F_{1,11} = 0.8386$ | 0.3794 | 0.0708 | [0.0005–0.3281] | 0.12 |
| **Experiment 6** | | | | | | |
| **Two-way ANOVA for HS image** | | **F value** | **P value** | **Partial η$^2$** | **95% CI of partial η$^2$** | **$BF_{10}$** |
| Main effects | Test | $F_{1,11} = 0.1067$ | 0.7500 | 0.0096 | [0.0000–0.1052] | 0.17 |
| | Orientation | $F_{1,11} = 1.3999$ | 0.2617 | 0.1129 | [0.0008–0.3893] | 0.45 |
| Two-way interaction | Test × Orientation | $F_{1,11} = 0.1331$ | 0.7221 | 0.0120 | [0.0000–0.1239] | 0.08 |
| **Experiment 7** | | | | | | |
| **Two-way ANOVA for GP image** | | **F value** | **P value** | **Partial η$^2$** | **95% CI of partial η$^2$** | **$BF_{10}$** |
| Main effects | Test | $F_{1,11} = 3.7718$ | 0.0782 | 0.2553 | [0.0002–0.6057] | 1.74 |
| | Orientation | $F_{1,11} = 0.1493$ | 0.7065 | 0.0134 | [0.0000–0.1352] | 0.19 |
| Two-way interaction | Test × Orientation | $F_{1,11} = 0.6241$ | 0.4462 | 0.0537 | [0.0000–0.3016] | 0.17 |
| **Experiment 9** | | | | | | |
| **Two-way ANOVA for lower-visibility KS image** | | **F value** | **P value** | **Partial η$^2$** | **95% CI of partial η$^2$** | **$BF_{10}$** |
| Main effects | Test | $F_{1,11} = 0.2442$ | 0.6309 | 0.0217 | [0.0000–0.1838] | 0.20 |
| | Orientation | $F_{1,11} = 0.0771$ | 0.7864 | 0.0070 | [0.0000–0.0681] | 0.16 |
| Two-way interaction | Test × Orientation | $F_{1,11} = 0.7715$ | 0.3985 | 0.0655 | [0.0000–0.3848] | 0.05 |

*P < 0.05, **P < 0.01, *** P < 0.005

#### Table S6: Results of ANOVA on functional connectivity in Experiment 12

| Four-way ANOVA | | F value | P value | Partial η$^2$ | 95% CI of partial η$^2$ | | $BF_{10}$ |
|---|---|---|---|---|---|---|---|
| Main effects | Task | $F_{1,27} = 0.0698$ | 0.7937 | 0.0026 | [0.0000–0.0284] | | 1.46 |
| | Image type | $F_{1,27} = 1.2121$ | 0.2806 | 0.0430 | [0.0001–0.2693] | | $< 10^{-5}$ |
| | Visual area | $F_{2.64,27} = 2.3652$ | 0.0857 | 0.0805 | [0.0097–0.2007] | | $< 10^{-5}$ |
| | Source region | $F_{2.49,27} = 6.0336$ | 0.0020 | 0.1827 | [0.0239–0.3390] | *** | 0.01 |
| Two-way interactions | Task × Image type | $F_{1,27} = 1.8562$ | 0.1843 | 0.0643 | [0.0000–0.2884] | | 10.50 |
| | Task × Visual area | $F_{2.79,75.25} = 0.4274$ | 0.7196 | 0.0156 | [0.0005–0.0373] | | $< 10^{-4}$ |

| | | | | | | |
|---|---|---|---|---|---|---|
| | Task × Source region | $F_{2.38,64.2} = 0.5794$ | 0.5919 | 0.0210 | [0.0001–0.0559] | $< 10^{-4}$ |
| | Image type × Visual area | $F_{2.6,70.23} = 0.7275$ | 0.5206 | 0.0262 | [0.0014–0.0649] | $< 10^{-5}$ |
| | Image type × Source region | $F_{3,81} = 0.2338$ | 0.8726 | 0.0086 | [0.0002–0.0200] | $< 10^{-5}$ |
| | Visual area × Source region | $F_{8.2,221.43} = 1.8989$ | 0.0596 | 0.0657 | [0.0232–0.0928] | $< 10^{-10}$ |
| Three-way interactions | Task × Image type × Visual area | $F_{2.89,77.95} = 0.9215$ | 0.4315 | 0.0330 | [0.0025–0.0969] | $< 10^{-10}$ |
| | Task × Image type × Source region | $F_{3,81} = 0.3087$ | 0.8190 | 0.0113 | [0.0001–0.0233] | $< 10^{-10}$ |
| | Task × Visual area × Source region | $F_{6.59,177.92} = 0.6167$ | 0.7319 | 0.0223 | [0.0101–0.0213] | $< 10^{-22}$ |
| | Image type × Visual area × Source region | $F_{6.9,186.26} = 0.5994$ | 0.7535 | 0.0217 | [0.0102–0.0177] | $< 10^{-22}$ |
| Four-way interaction | Task × Image type × Visual area × Source region | $F_{5.99,161.73} = 0.9893$ | 0.4343 | 0.0353 | [0.0149–0.0506] | $< 10^{-45}$ |

*P < 0.05, **P < 0.01, *** P < 0.005

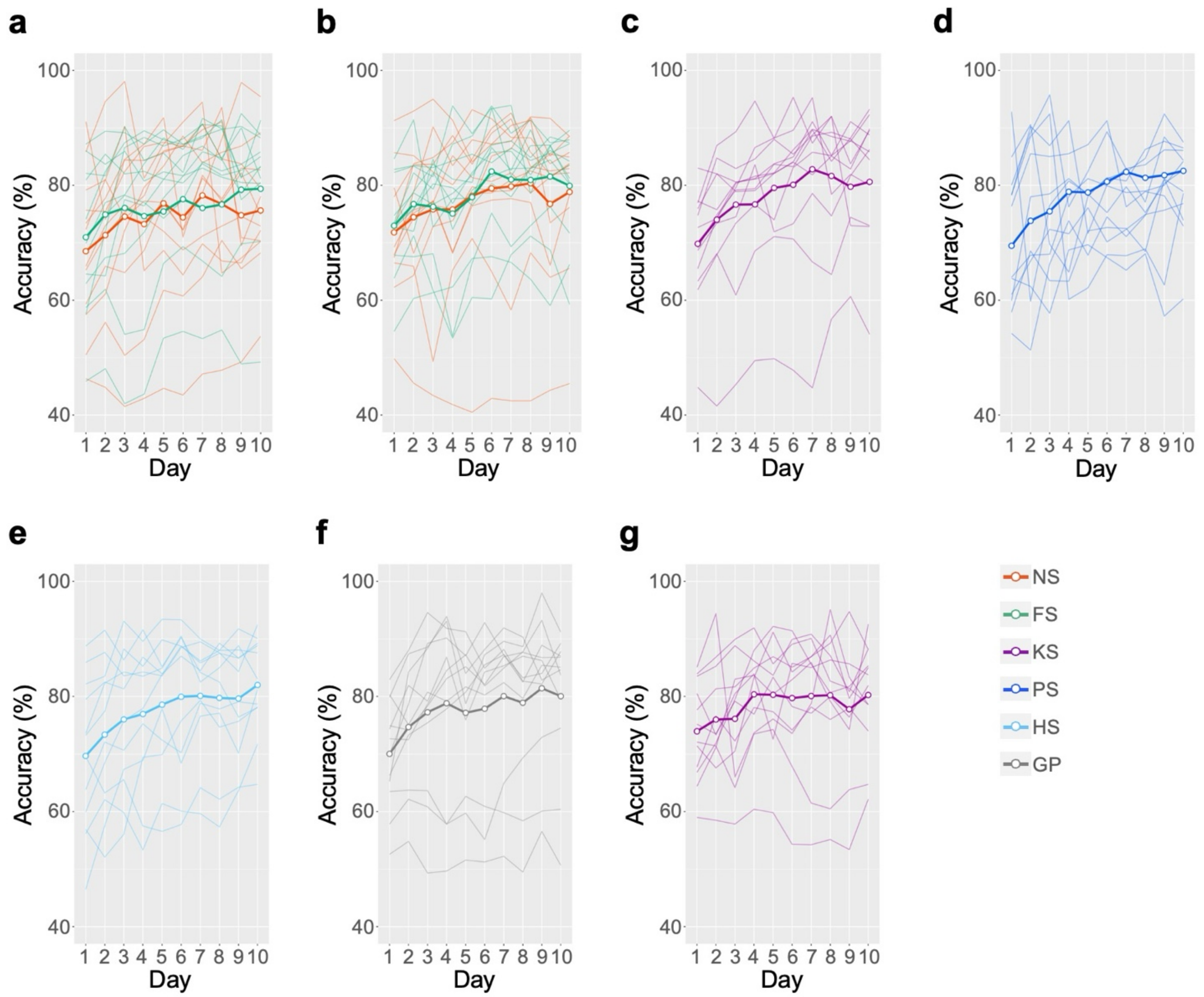


**Fig S1: Supplementary results of Experiments 1, 2, 4-7, 9**

**a-g**, Accuracies of RSVP task during the exposure+task stages for Experiments 1 (a), 2 (b), 4 (c), 5 (d), 6 (e), 7 (f), and 9 (g). Thick lines indicate mean accuracies across participants, and thin lines represent individual accuracies. NS, FS, KS, PS, HS, and GP represent natural scene, Fourier-scrambled, kurtosis- & skewness-matched, Portilla-Simoncelli, higher-order statistics, and Gabor patch, respectively.

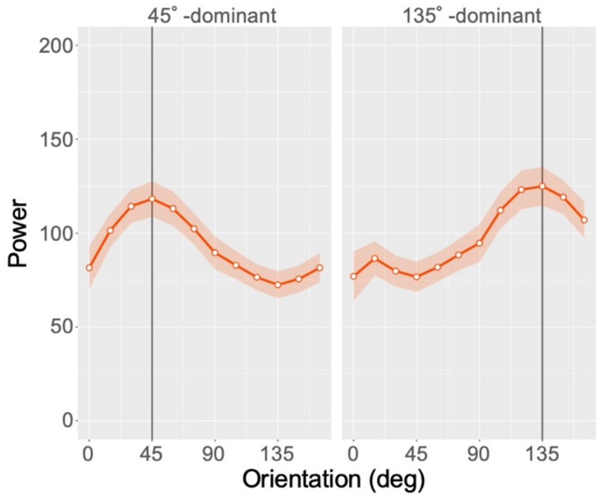


**Fig. S2: Orientation power distribution of NS images**

Left and right panels show orientation power distribution computed using a Sobel filter based on 45°-dominant and 135°-dominant NS images, respectively. Solid lines indicate average values across 40 images, and shading represents 95% confidence interval.

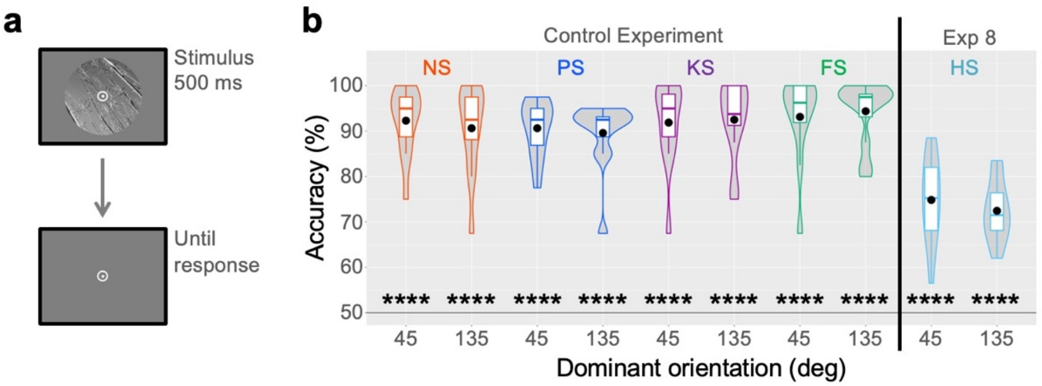


**Fig. S3: Procedures and results of Control Experiment and Experiment 8**

**a**, Procedure of the orientation discrimination task. In each trial, participants were asked to report whether the dominant orientation of a presented image was rotated in a clockwise or counterclockwise direction from the vertical axis. A different group of participants was employed for each of Control Experiment (N = 12) and Experiment 8 (N = 12). **b**, Accuracy of the orientation discrimination task in Control Experiment (NS, PS, KS, and FS images) and Experiment 8 (HS image). In Control Experiment, mean accuracies were significantly higher than the chance level for all image types (one-sample *t*-tests with Bonferroni correction for multiple comparisons; $t_{11} > 17.4770$, corrected $P < 10^{-4}$, Cohen's $d > 5.0452$, $BF_{10} > 10^{6}$), indicating that the dominant orientations of these images are supra-threshold. We also tested whether the orientation information in images differs across the types of images (NS, PS, KS, vs FS images) and orientations (45°- vs 135°-dominant). A two-way repeated-measures ANOVA on accuracies revealed no significant effects of Image ($F_{1.89,20.77} = 1.5060$, $P = 0.2449$, partial $\eta^2 = 0.1204$, 95% CI of partial $\eta^2$ = [0.0061–0.2845], $BF_{10} = 0.17$), Orientation ($F_{1,11} = 0.0158$, $P = 0.9023$, partial $\eta^2 = 0.0014$, 95% CI of partial $\eta^2$ = [0.0000–0.0112], $BF_{10} = 0.11$), or interaction between them ($F_{1,11} = 0.8849$, $P = 0.4041$, partial $\eta^2 = 0.0745$, 95% CI of partial $\eta^2$ = [0.0018–0.2068], $BF_{10} = 0.01$). Thus, the differences in learning based on exposure to the task-irrelevant images among the groups in Experiments 1-5 cannot be attributed to differences in orientation information across the image types. In Experiment 8, mean accuracies were significantly higher than the chance level for the 45°-dominant HS images (one-sample *t*-test with Bonferroni correction for multiple comparisons; $t_{11} = 9.2317$, corrected $P < 10^{-4}$, Cohen's $d = 8.0307$, 95% CI of accuracy = [68.9127–80.7540] %, $BF_{10} = 10726.38$) and the 135°-dominant HS images ($t_{11} = 11.6124$, corrected $P < 10^{-4}$, Cohen's $d = 10.8153$, 95% CI of accuracy = [68.2016–76.7151] %, $BF_{10} = 83865.74$). No significant difference was found between the orientations (paired *t*-test; $t_{11} = 0.7901$, $P < 0.4461$, Cohen's $d = 0.2281$, 95% CI of the difference in accuracies = [-4.2406–8.9910] %, $BF_{10} = 0.375$). The mean accuracies appear to be different between the HS image (Experiment

8) and the other types of images (Control Experiment). We confirmed this observation, as follows. First, based on no significant differences in mean accuracies among the images in Control Experiment, these accuracies were averaged across the images within Control Experiment. Second, we applied a two-way mixed-model ANOVA to accuracies with Group (Control Experiment vs Experiment 8) as the between-participant factor and Orientation (45°- vs 135°-dominant) as the within-participant factor. The ANOVA revealed significant effects of Group ($F_{1,22}$ = 48.9962, $P < 10^{-4}$, partial $\eta^2$ = 0.6901, 95% CI of partial $\eta^2$ = [0.3880–0.8203], $BF_{10} > 10^4$), but no main effect of Orientation ($F_{1,1}$ = 0.5664, $P < 0.4597$, partial $\eta^2$ = 0.0251, 95% CI of partial $\eta^2$ = [0.0000–0.1792], $BF_{10}$ = 0.25) or interaction between Group and Orientation ($F_{1,22}$ = 0.3984, P = 0.5344, partial $\eta^2$ = 0.0178, 95% CI of partial $\eta^2$ = [0.0000–0.1577], $BF_{10}$ = 0.29). These results suggest that the orientation information differs between the images in Control Experiment and Experiment 8. NS, PS, KS, FS, and HS represent natural scene, Portilla-Simoncelli, kurtosis- & skewness-matched, Fourier-scrambled, and higher-order statistics, respectively. Box plots are overlaid on violin plots. Black dots represent mean accuracies across participants, and violin plots show kernel probability densities of individual accuracies. ****P < 0.0001.

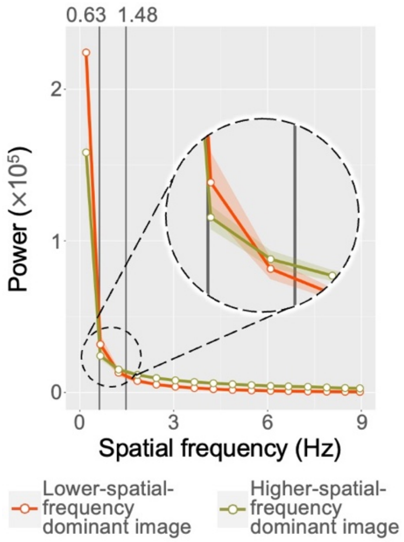


**Fig. S4: Supplementary methods of Experiment 2**

Frequency power distributions computed using Fourier transformation for lower-spatial-frequency dominant (orange) and higher-spatial-frequency dominant (yellow) NS images. Thin gray vertical lines indicate the mean normalized spatial frequencies for the lower- (0.63 Hz) and higher- (1.48 Hz) spatial-frequency dominant images. Solid lines indicate average frequency power across 10 images, and shading represents 95% confidence interval.

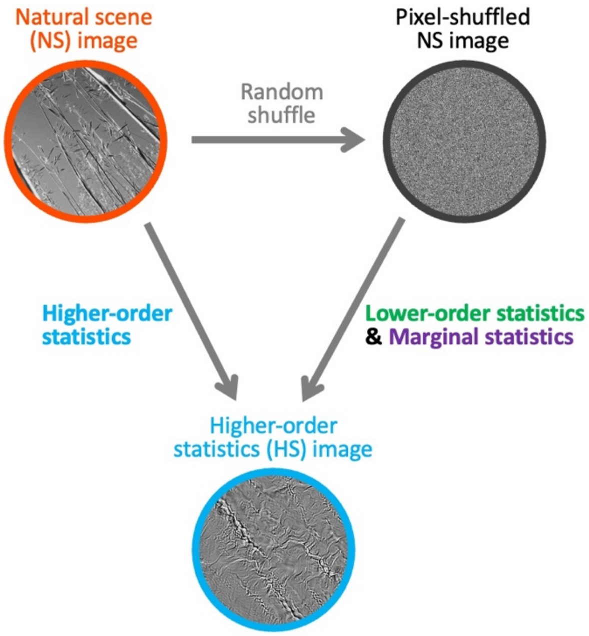


**Fig. S5: Supplementary methods of Experiment 6**

Natural scene images are characterized by higher-order statistics, marginal statistics and lower-order statistics. Higher-order statistics convey complex spatial relationships such as edges, textures, and contours typical of natural scenes (e.g., [1,2]). Marginal statistics include kurtosis, which reflects the "tailedness" or peakedness, and skewness, which indicates asymmetry of the luminance histogram of natural scenes (e.g., [3,4]). Lower-order statistics refer to properties such as orientation and spatial frequency that are locally and independently processed in the visual system (e.g., [5,6]). An HS image was generated as follows. First, higher-order statistics were calculated based on a target NS image. Second, the pixels of the target NS image were spatially shuffled. Third, lower-order statistics and marginal statistics were calculated based on the shuffled NS image. Finally, the Portilla-Simoncelli algorithm[2] (see Stimuli in Methods for details) was used to synthesize the HS image based on the higher-order statistics from the target NS image and the lower-order and marginal statistics from the shuffled NS image. This procedure effectively removed the original spatial frequency and orientation structure of the target NS image from the HS image, while preserving the higher-order and marginal statistics.

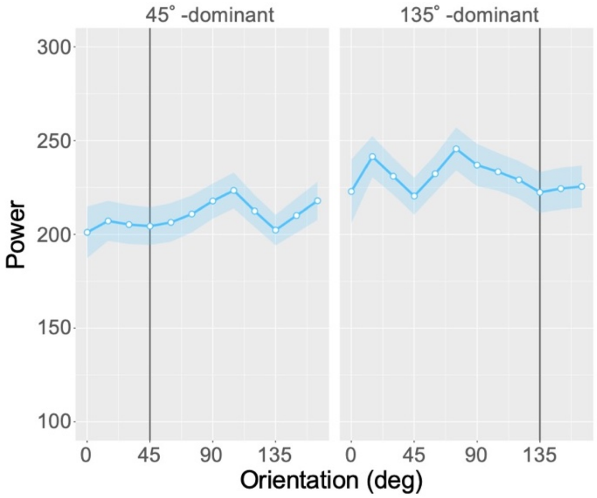


**Fig. S6: Orientation power distribution of HS images**

Left and right panels show orientation power distribution computed using a Sobel filter based on 45°-dominant and 135°-dominant HS images, respectively. Solid lines indicate average values across 40 images, and shading represents 95% confidence interval.

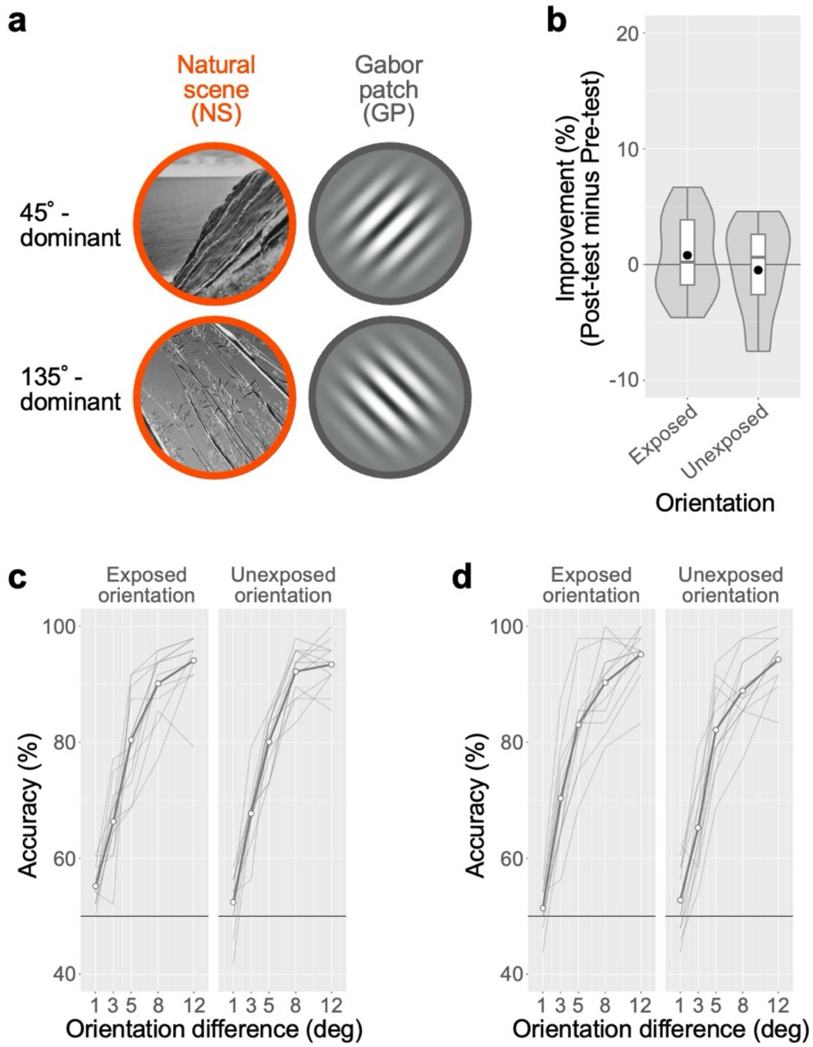


**Fig. S7: Methods and results of Experiment 7**

**a**, Examples of NS images (highlighted with orange rings) and Gabor patch images (GP, highlighted with gray rings). **b**, Improvement in performance on the orientation discrimination task (post-test minus pre-test) for the GP group (N = 12). A two-way repeated-measures ANOVA revealed no significant effects of Test (pre vs post), Orientation (exposed vs unexposed), or their interaction (See Table S1 for details). Box plots are overlaid on violin plots. Black dots represent mean performance improvements across participants, and violin plots show kernel probability densities of individual performance improvements. **c**, Performance in the orientation discrimination task in the pre-test stage. **d**, Performance in the post-test stage. Thick lines indicate mean accuracies across participants, and thin lines represent individual accuracies.

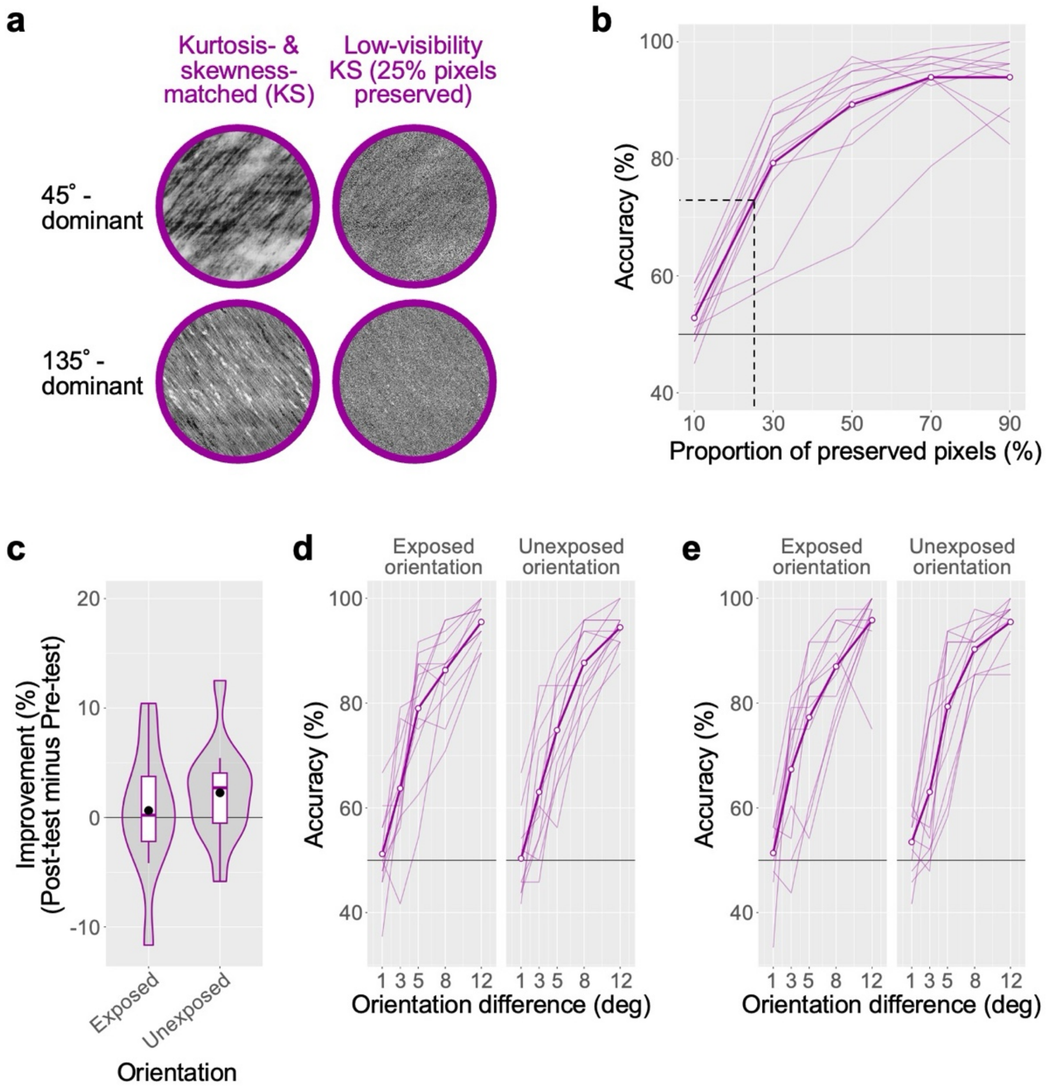


**Fig. S8: Methods and results of Experiment 9**

**a**, Examples of KS and low-visibility KS images. **b**, Accuracy of the orientation discrimination task in a pilot Experiment (N = 12). Dotted lines indicate the proportion of preserved pixels (25%) at which the mean accuracy in the orientation discrimination task with the low-visibility KS images was equivalent to that with the HS images (73.7%; Fig. S3b). Thick lines represent the mean accuracy across participants, and thin lines indicate individual data. **c**, Improvement in performance on the orientation discrimination task (post-test minus pre-test) for the low-visibility KS group (N = 12). A two-way repeated-measures ANOVA revealed no significant effects of Test (pre vs post) or Orientation (exposed vs unexposed), and no significant interaction between them (See Table S1 for details). **d**, Performance in the orientation discrimination task in the pre-test stage. **e**, Performance in the post-test stage. In panels d and e, thick lines represent mean accuracies across participants, and thin lines indicate individual accuracies. Box plots are overlaid on violin plots. Black dots represent mean performance improvements across participants, and violin plots show kernel probability densities of individual performance improvements. Thick lines indicate mean accuracies across participants, and thin lines represent individual accuracies.

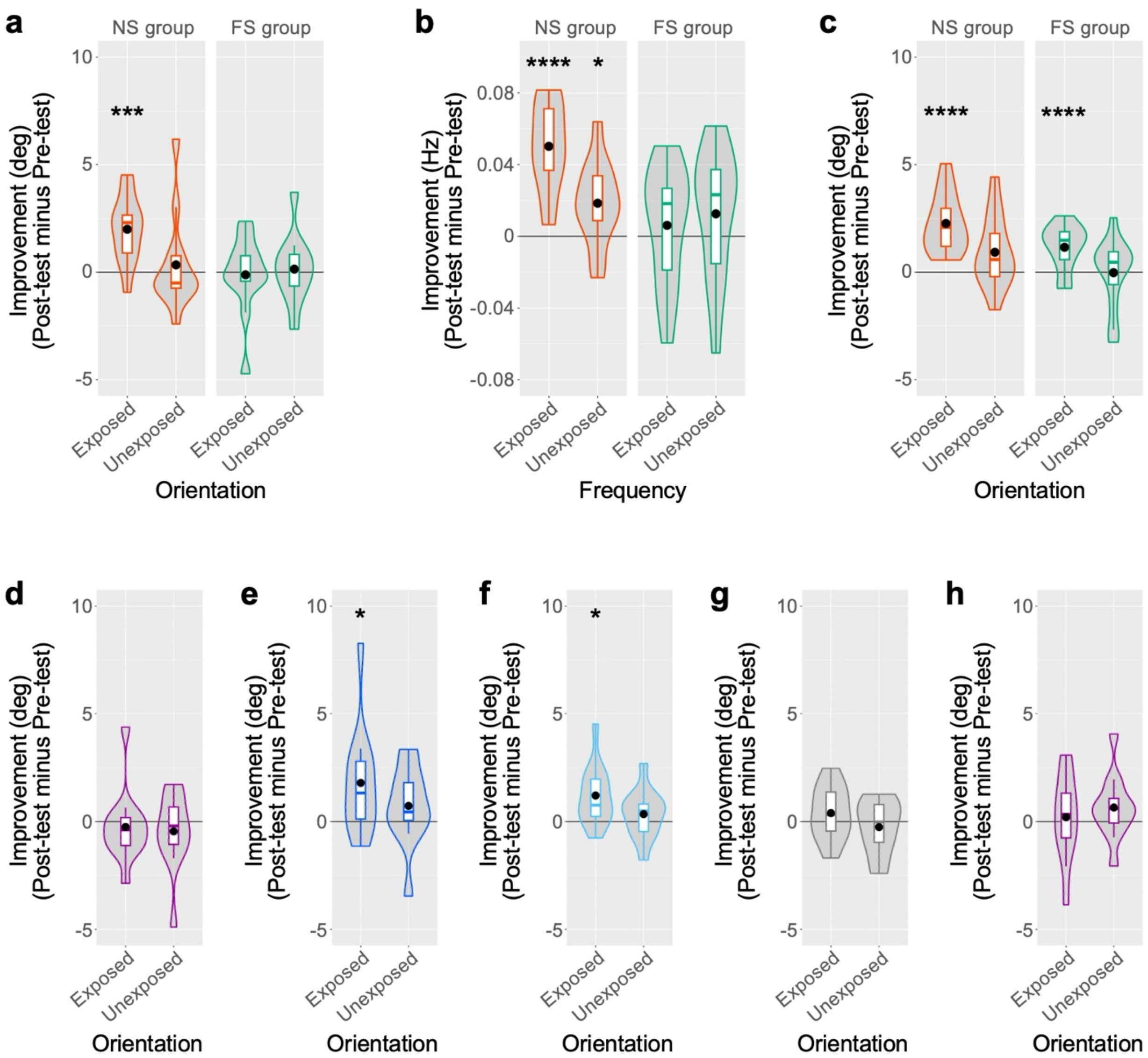


**Fig. S9: Changes in perceptual thresholds calculated by curve fitting**

In Experiments 1-7 and 9 (**a**-**h**), an ANOVA was applied to perceptual thresholds in the pre- and post-test stages. Detailed results of the ANOVAs are provided in Table S2. In the pre- and post-test stages, participants performed the spatial frequency discrimination task in Experiment 2 (**b**) and the orientation discrimination task in the other experiments. In Experiments 1 (**a**), 2 (**b**), 3 (**c**), 5 (**e**), and 6 (**f**), we found a significant main effect and/or interaction. Accordingly, the results of subsequent simple effect tests are shown in the figure captions. In the remaining experiments, no significant main effects or interactions were observed. **a**, In Experiment 1, participants were exposed to natural scene (NS) and Fourier-scrambled (FS) images in the exposure+task stage. For the NS group, we found a significant simple effect of Test at the exposed orientation ($F_{1,11}$ = 19.9255, P = 0.0010, partial $\eta^2$ = 0.6443, 95% CI of partial $\eta^2$ = [0.1985–0.8371]). **b**, In Experiment

2, participants were exposed to NS and FS images in the exposure+task stage. For the NS group, we found a significant simple effect of Test at the exposed ($F_{1,11}$ = 49.7857, $P < 10^{-4}$, partial $\eta^2$ = 0.8190, 95% CI of partial $\eta^2$ = [0.5975–0.9191]) and unexposed ($F_{1,11}$ = 7.4117, P = 0.0198, partial $\eta^2$ = 0.4026, 95% CI of partial $\eta^2$ = [0.0049–0.7165]) frequencies. **c**, In Experiment 3, participants were exposed to NS and FS images in the exposure stage. We found a significant interaction between Test and Orientation. **d**, In Experiment 4, participants were exposed to kurtosis- and skewness-matched (KS) images in the exposure+task stage. **e**, In Experiment 5, participants were exposed to the Portilla-Simoncelli (PS) images in the exposure+task stage. We found a significant simple effect of Test at the exposed orientation ($F_{1,11}$ = 6.2277, P = 0.0297, partial $\eta^2$ = 0.3615, 95% CI of partial $\eta^2$ = [0.0741–0.5987]). **f**, In Experiment 6, participants were exposed to higher-order statistics (HS) images in the exposure+task stage. We found a significant simple effect of Test at the exposed orientation ($F_{1,11}$ = 9.1053, P = 0.0117, partial $\eta^2$ = 0.4529, 95% CI of partial $\eta^2$ = [0.0909–0.6611]). **g**, In Experiment 7, participants were exposed to Gabor patch (GP) images in the exposure+task stage. **h**, In Experiment 9, participants were exposed to low-visibility KS images in the exposure+task stage. Box plots are overlaid on violin plots. Black dots represent mean threshold improvements across participants, and violin plots show kernel probability densities of individual performance improvements. *P < 0.05, ***P < 0.005, ****P < 0.001.

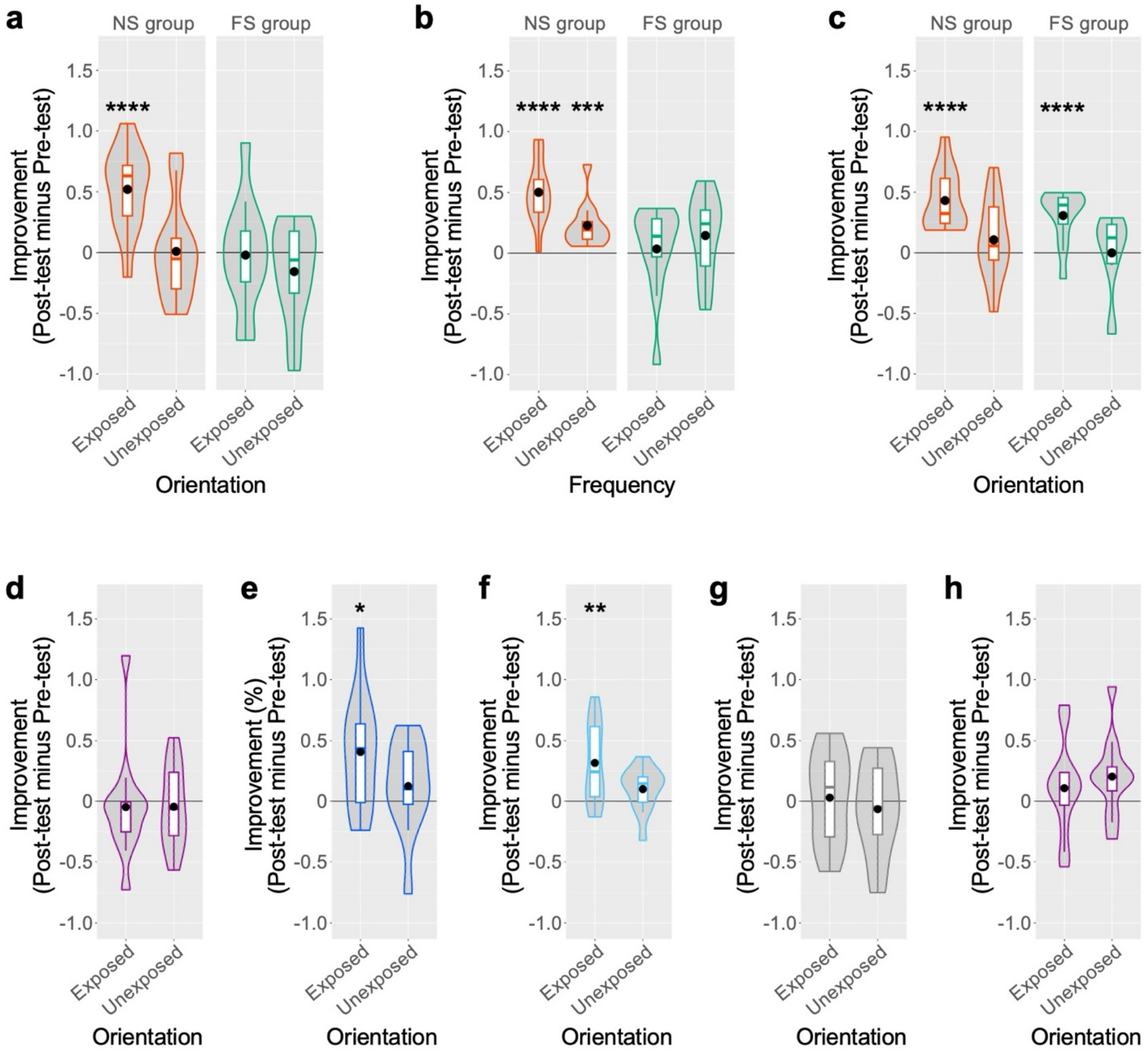


**Fig. S10: Changes in d’**

In Experiments 1-7 and 9 (**a**-**h**), an ANOVA was applied to d’ in the pre- and post-test stages. Detailed results of the ANOVAs are provided in Table S4. In the pre- and post-test stages, participants performed the spatial frequency discrimination task in Experiment 2 (**b**) and the orientation discrimination task in the other experiments. In Experiments 1 (**a**), 2 (**b**), 3 (**c**), 5 (**e**), and 6 (**f**), we found a significant main effect and/or interaction. Accordingly, the results of subsequent simple effect tests are shown in the figure captions. In the remaining experiments, no significant main effects or interactions were observed. **a**, In Experiment 1, participants were exposed to natural scene (NS) and Fourier-scrambled (FS) images in the exposure+task stage. For the NS group, we found a significant simple effect of Test at the exposed orientation ($F_{1,11}$ = 27.5581, P = 0.0003, partial $\eta^2$ = 0.7147, 95% CI of partial $\eta^2$ = [0.2949–0.8794]). **b**, In Experiment

2, participants were exposed to NS and FS images in the exposure+task stage. For the NS group, we found a significant simple effect of Test at the exposed ($F_{1,11}$ = 45.1154, $P < 10^{-4}$, partial $\eta^2$ = 0.8040, 95% CI of partial $\eta^2$ = [0.5312–0.8928]) and unexposed ($F_{1,11}$ = 18.1174, P = 0.0014, partial $\eta^2$ = 0.6222, 95% CI of partial $\eta^2$ = [0.4136–0.8002]) frequencies. **c**, In Experiment 3, participants were exposed to NS and FS images in the exposure stage. We found a significant interaction between Test and Orientation. **d**, In Experiment 4, participants were exposed to kurtosis- and skewness-matched (KS) images in the exposure+task stage. **e**, In Experiment 5, participants were exposed to the Portilla-Simoncelli (PS) images in the exposure+task stage. We found a significant simple effect of Test at the exposed orientation ($F_{1,11}$ = 9.0781, P = 0.0118, partial $\eta^2$ = 0.4521, 95% CI of partial $\eta^2$ = [0.0905–0.7010]). **f**, In Experiment 6, participants were exposed to higher-order statistics (HS) images in the exposure+task stage. We found a significant simple effect of Test at the exposed orientation ($F_{1,11}$ = 11.2595, P = 0.0064, partial $\eta^2$ = 0.5058, 95% CI of partial $\eta^2$ = [0.2193–0.7474]). **g**, In Experiment 7, participants were exposed to Gabor patch (GP) images in the exposure+task stage. **h**, In Experiment 9, participants were exposed to low-visibility KS images in the exposure+task stage. Box plots are overlaid on violin plots. Black dots represent mean threshold improvements across participants, and violin plots show kernel probability densities of individual performance improvements. *P < 0.05, **P < 0.01, ***P < 0.005, ****P < 0.001.

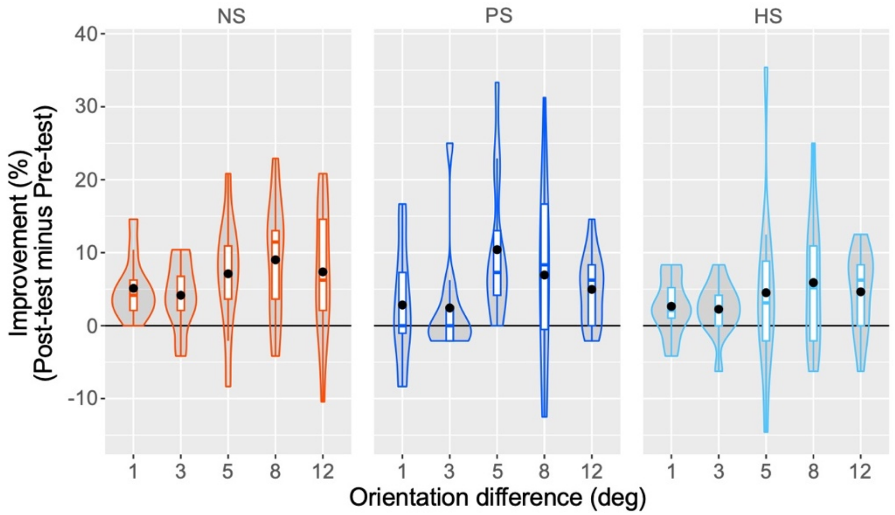


**Fig. S11: Comparison of performance improvements across Experiments 1, 5, and 6**

Left, middle, and right panels show performance improvements on the orientation discrimination task (post-test minus pre-test) for the exposed orientation at the different levels of task difficulties (the Δ values of 12, 8, 5, 1 degree; see Experiment 1 and Methods for details) from NS group (Experiment1), PS group (Experiment 5), and HS group (Experiment 6), respectively. We applied a three-way mixed-model ANOVA to accuracies with Test (pre vs post) and Difficulty (12, 8, 5, vs 1) as the within-participant factors and Group (NS, PS, vs HS) as the between-participant factor. We found significant effects of Test ($F_{1,33} = 46.8927$, $P < 10^{-4}$, partial $\eta^2 = 0.5869$, 95% CI of partial $\eta^2$ = [0.3544–0.7105], $BF_{10} > 10^{11}$) and Difficulty ($F_{2.61,86.09} = 433.2889$, $P < 10^{-4}$, partial $\eta^2 = 0.9292$, 95% CI of partial $\eta^2$ = [0.9029–0.9419], $BF_{10} > 10^{139}$) and a significant interaction between Test and Difficulty ($F_{3.1,102.2} = 3.6159$, $P = 0.0148$, partial $\eta^2 = 0.0988$, 95% CI of partial $\eta^2$ = [0.0167–0.1719], $BF_{10} = 0.67$). We found no significant effect of Group ($F_{2,33} = 1.0688$, $P = 0.3550$, partial $\eta^2 = 0.0608$, 95% CI of partial $\eta^2$ = [0.0006–0.2165], $BF_{10} = 0.08$), interaction between Test and Group ($F_{2,33} = 0.8974$, $P = 0.4173$, partial $\eta^2 = 0.0516$, 95% CI of partial $\eta^2$ = [0.0001–0.1663], $BF_{10} = 0.03$), interaction between Difficulty and Group ($F_{5.22,86.09} = 1.0487$ $P = 0.3959$, partial $\eta^2 = 0.0598$, 95% CI of partial $\eta^2$ = [0.0107–0.0918], $BF_{10} = 0.01$), or interaction between the three factors ($F_{6.19,102.2} = 0.3993$, $P = 0.8828$, partial $\eta^2 = 0.0236$, 95% CI of partial $\eta^2$ = [0.0089–0.0266], $BF_{10} < 10^{-5}$). Subsequent analyses revealed significant simple effects of Test at all difficulty levels ($F_{1,33} > 11.2911$, $P < 0.0020$, partial $\eta^2 < 0.2549$, $BF_{10} > 20.18$). Since the NS, PS, and HS images all contained the higher-order statistics, these results suggest that VPL of task-irrelevant images that contained higher-order statistics alters processing of the dominant orientation across a broad range of difficulty levels. Box plots are overlaid on violin plots. Black dots represent mean performance improvements across participants, and violin plots show kernel probability densities of individual performance improvements.

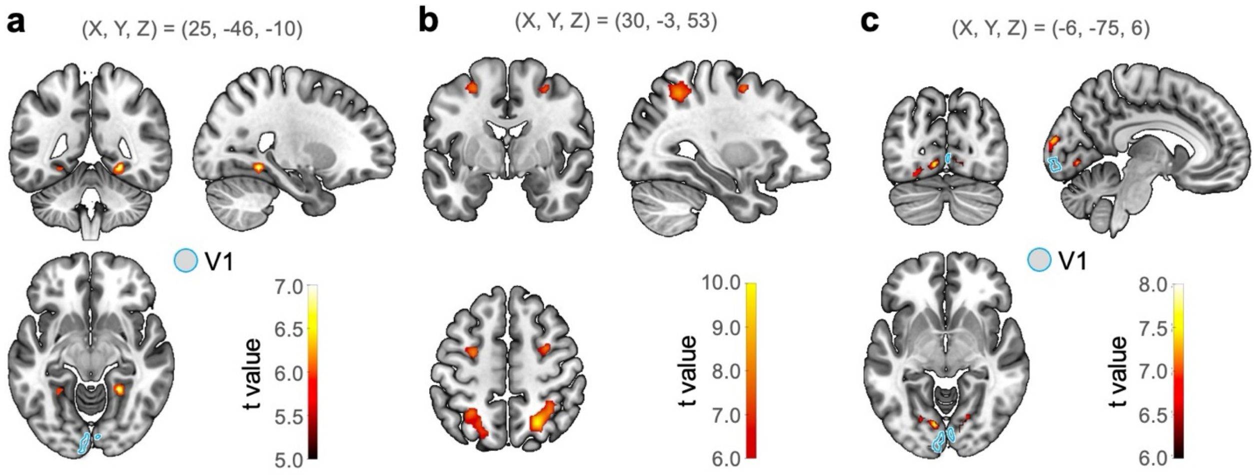


**Fig. S12: Supplementary results of whole-brain analyses in Experiment 11 (N = 28)**

**a**, Colored voxels show areas where BOLD signal amplitudes in response to task-irrelevant KS images were significantly higher than those to HS images (paired t-test, $P < 0.05$ after Bonferroni correction for multiple comparisons). **b**, BOLD signal amplitudes during the harder RSVP task were significantly higher than those during the easy RSVP task (corrected $P < 0.05$). **c**, BOLD signal amplitudes during the easy RSVP task were significantly higher than those during the harder RSVP task condition (corrected $P < 0.05$). Regions enclosed by blue lines represent voxels identified as belonging to V1 in more than half of the 28 participants.

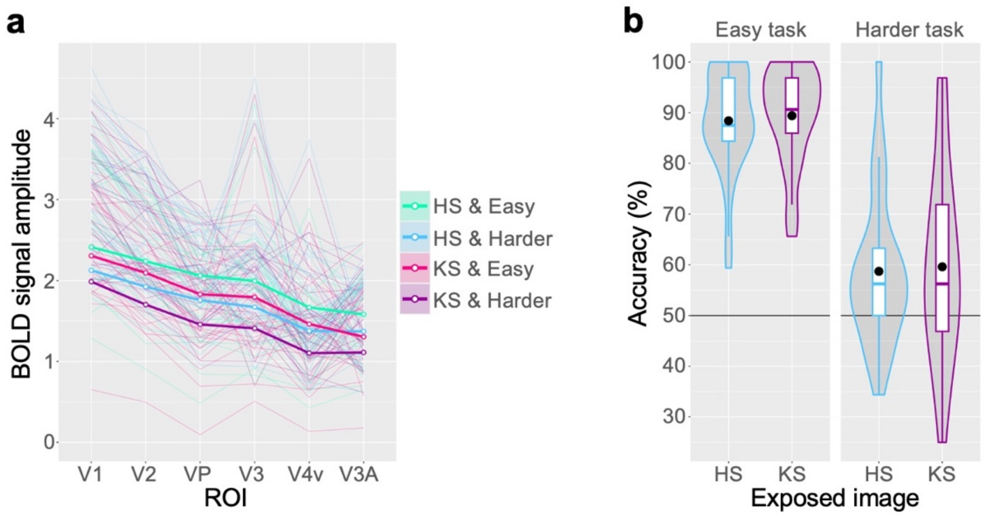


**Fig. S13: Supplementary results of ROI analyses in Experiment 11**

**a**, BOLD signal amplitudes in HS image & Easy RSVP task, HS image & Harder RSVP task, KS image & Easy RSVP task, and KS image & Harder RSVP task conditions are shown. Thick lines indicate mean BOLD signal amplitudes across participants, and thin lines represent individual BOLD signal amplitudes. **b**, Accuracies of easy (left panel) and harder (right panel) RSVP tasks. To compare accuracies across the conditions and images, we applied a two-way repeated-measures ANOVA to accuracies with the factors of Task (easy vs harder RSVP) and Image (HS vs KS). As expected, we found a significant main effect of Task ($F_{1,27}$ = 140.5582, $P < 10^{-4}$, partial $\eta^2$ = 0.8389, 95% CI of partial $\eta^2$ = [0.7062–0.9155], $BF_{10}$ = 140.56), but no significant main effect of Image ($F_{1,27}$ = 0.5232, P = 0.4757, partial $\eta^2$ = 0.0190, 95% CI of partial $\eta^2$ = [0.0000–0.1795], $BF_{10}$ = 0.13) or interaction between the factors ($F_{1,27}$ = 0.0019, P = 0.9653, partial $\eta^2$ = 0.0001, 95% CI of partial $\eta^2$ = [0.0000–0.0001], $BF_{10}$ = 0.11). Box plots are overlaid on violin plots. Black dots represent mean accuracies across participants, and violin plots show kernel probability densities of individual accuracies.

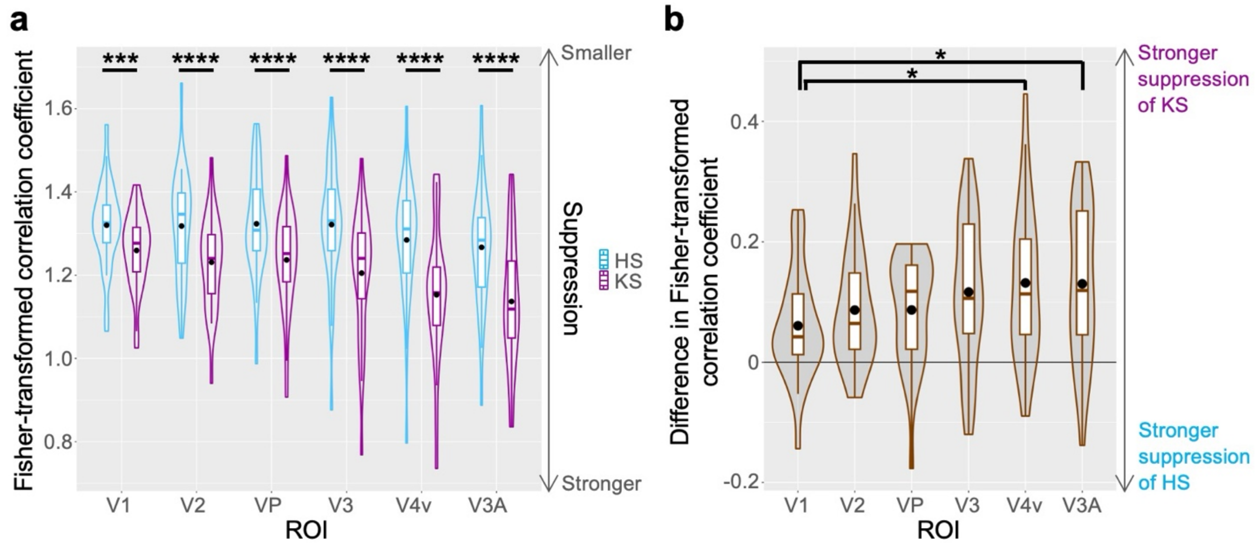


**Fig. S14: Results of RSA in Experiment 11**

**a**, Fisher-transformed correlation coefficients of BOLD signal patterns between easy and harder RSVP tasks (see Method for detailed calculation) for HS and KS images in each of regions of interest (ROIs). We applied a two-way repeated-measures ANOVA to correlation coefficients with the factors of ROI (V1, V2, VP, V3, V4v, vs V3A) and Image (HS vs KS). The ANOVA revealed significant effects of ROI ($F_{3.54,95.47}$ = 7.2060, P = 0.0001, partial $\eta^2$ = 0.2017, 95% CI of partial $\eta^2$ = [0.0667–0.3341], $BF_{10} > 10^5$), Image ($F_{1,27}$ = 40.1152, P <$10^{-4}$, partial $\eta^2$ = 0.5977, 95% CI of partial $\eta^2$ = [0.3751–0.7292], $BF_{10} > 10^{19}$), and their interaction ($F_{3.64,98.25}$ = 3.9294, P = 0.0068, partial $\eta^2$ = 0.1270, 95% CI of partial $\eta^2$ = [0.0363–0.2122], $BF_{10}$ = 0.27). Subsequent analyses revealed significant simple effects of Image at V1 ($F_{1,27}$ = 12.4953, P = 0.0015, partial $\eta^2$ = 0.3164, 95% CI of partial $\eta^2$ = [0.0688–0.5156], $BF_{10}$ = 23.72), V2 ($F_{1,27}$ = 23.2399, P <$10^{-4}$, partial $\eta^2$ = 0.4626, 95% CI of partial $\eta^2$ = [0.2717–0.6088], $BF_{10}$ = 498.65), VP ($F_{1,27}$ = 25.5884, P <$10^{-4}$, partial $\eta^2$ = 0.4866, 95% CI of partial $\eta^2$ = [0.1330–0.7030], $BF_{10}$ = 890.80), V3 ($F_{1,27}$ = 25.9898., P <$10^{-4}$, partial $\eta^2$ = 0.4905, 95% CI of partial $\eta^2$ = [0.2400–0.6752], $BF_{10}$ = 981.15), V4v ($F_{1,27}$ = 29.8736, P <$10^{-4}$, partial $\eta^2$ = 0.5253, 95% CI of partial $\eta^2$ = [0.3296–0.6689], $BF_{10}$ = 2412.30), and V3a ($F_{1,27}$ = 29.5797, P <$10^{-4}$, partial $\eta^2$ = 0.5228, 95% CI of partial $\eta^2$ = [0.2633–0.6938], $BF_{10}$ = 2258.39). **b**, Subtraction of the fisher-transformed correlation coefficients for the KS images from those for the HS images in each ROI. A one-way repeated-measures ANOVA revealed a significant effect of ROI ($F_{3.64,98.25}$ = 3.9294, P = 0.0068, partial $\eta^2$ = 0.1270, 95% CI of partial $\eta^2$ = [0.0335–0.2154], $BF_{10}$ = 7.57). Subsequent analyses revealed significant differences in the subtractions between V1 and V4v (Shaffer's Modified Sequentially Rejective Bonferroni Procedure; $t_{27}$ = 3.3161, corrected P = 0.0345, $BF_{10}$ = 14.57) and between V1 and V3A ($t_{27}$ = 3.3661, corrected P = 0.0345, $BF_{10}$ = 16.26). Box plots are overlaid on violin plots. Black dots indicate mean values across participants, and violin plots show kernel probability densities of individual data. VP: ventral posterior area. *P < 0.05, ***P < 0.01, ****P < 0.001.

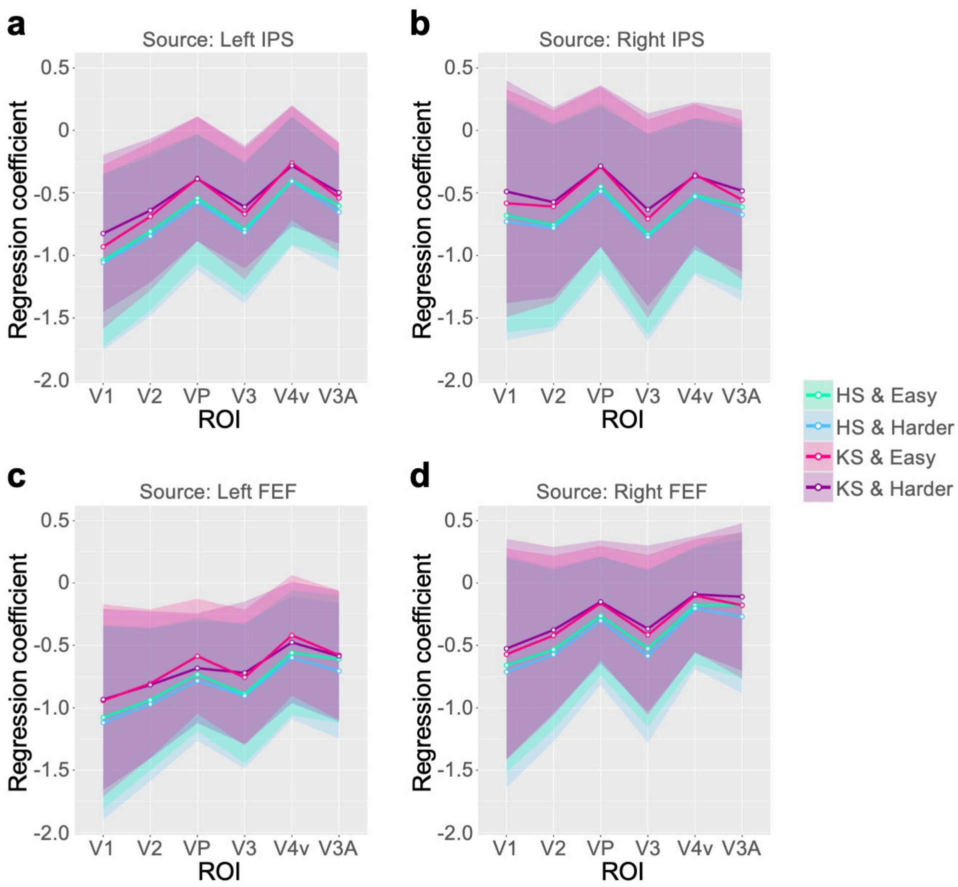


**Fig. S15: Results of functional connectivity analyses in Experiment 11**

**a-d**, Functional connectivity, defined as regression coefficients, between the left intraparietal sulcus (IPS) and visual areas (a), between the right IPS and visual areas (b), between the left frontal eye field (FEF) and visual areas (c), and between the right FEF and visual areas (d). We applied a four-way repeated-measures ANOVA to regression coefficients with the factors of Source regions (left IPS, right IPS, left FEF, vs right FEF), Visual area (V1, V2, VP, V3, V4v, vs V3A), Task (easy vs harder RSVP), and Image type (HS vs KS). We found a significant effect of Source region, but no significant effects of the other factors or interactions (see Table S2 for details). Shading represents 95% confidence intervals.

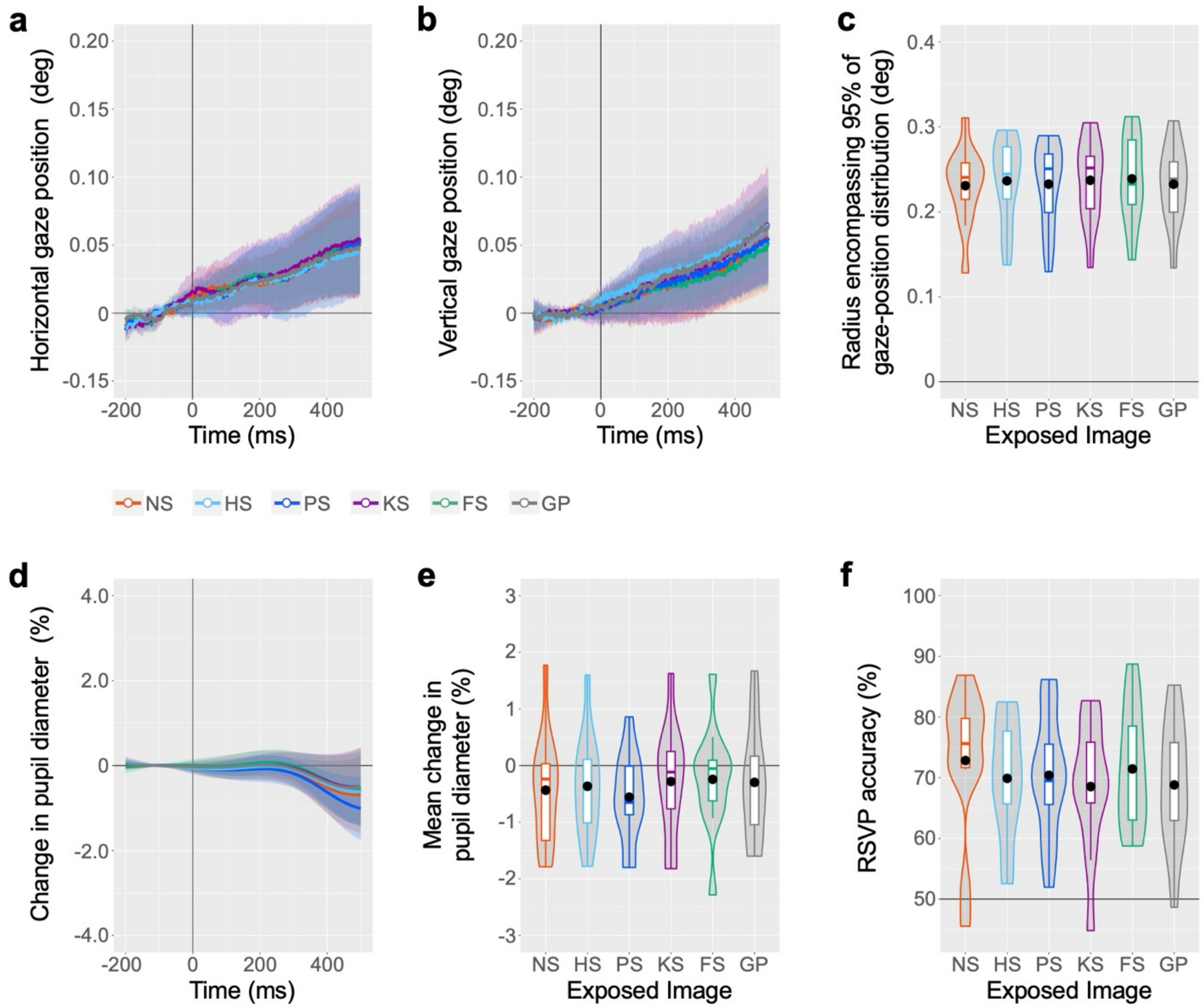


**Fig. S16: Results of Experiment 13**

**a**, Horizontal gaze-position time series when the natural scene (NS, orange), higher-order statistics (HS, cyan), Portilla-Simoncelli (PS, blue), kurtosis- and skewness-matched (KS, purple), Fourier-scrambled (FS, green), and Gabor patch (GP, gray) images were presented in the background. Solid lines indicate mean gaze positions, and shading represents 95% confidence interval across participants. The onset of the 500-ms stimulus period was set at 0 ms. **b**, Vertical gaze-position time series. **c**, Radius encompassing the gaze-position distribution with a 95% confidence range during the 500-ms stimulus period. A one-way repeated-measures ANOVA on radius revealed no significant effect of Image (NS, HS, PS, KS FS, vs GP; $F_{4.02,44.21}$ = 0.4153, P = 0.7976, partial $\eta^2$ = 0.0364, 95% CI of partial $\eta^2$ = [0.0037–0.0619], $BF_{10}$ = 0.0346). **d**, Time series of changes in pupil diameter. Solid lines indicate mean changes in pupil diameters, and shading represents 95% confidence interval across participants. **e**, Mean changes in pupil diameter during the 500-ms stimulus period. A one-way repeated-measures ANOVA on changes in pupil diameter revealed no significant effect of Image ($F_{3.37,37.10}$ = 1.2619, P = 0.3022, partial $\eta^2$ = 0.1029, 95% CI of partial $\eta^2$ = [0.0178–0.1921], $BF_{10}$ = 0.1374). **f**, RSVP task performance with each of the six background images. A one-way repeated-measures ANOVA revealed no significant effect of Image ($F_{3.36,36.99}$ = 1.4036, P = 0.2554, partial $\eta^2$ = 0.1132, 95% CI of partial $\eta^2$ = [0.0113–0.1985], $BF_{10}$ = 0.1725).
